# *Atp1a2* and *Kcnj9* are candidate genes underlying sensitivity to oxycodone-induced locomotor activation and withdrawal-induced anxiety-like behaviors in C57BL/6 substrains

**DOI:** 10.1101/2024.04.16.589731

**Authors:** Lisa R. Goldberg, Britahny M. Baskin, Yahia Adla, Jacob A. Beierle, Julia C. Kelliher, Emily J. Yao, Stacey L. Kirkpatrick, Eric R. Reed, David F. Jenkins, Jiayi Cox, Alexander M. Luong, Kimberly P. Luttik, Julia A. Scotellaro, Timothy A. Drescher, Sydney B. Crotts, Neema Yazdani, Martin T. Ferris, W. Evan Johnson, Megan K. Mulligan, Camron D. Bryant

## Abstract

Opioid use disorder is heritable, yet its genetic etiology is largely unknown. C57BL/6J and C57BL/6NJ mouse substrains exhibit phenotypic diversity in the context of limited genetic diversity which together can facilitate genetic discovery. Here, we found C57BL/6NJ mice were less sensitive to oxycodone (OXY)-induced locomotor activation versus C57BL/6J mice in a conditioned place preference paradigm. Narrow-sense heritability was estimated at 0.22-0.31, implicating suitability for genetic analysis. Quantitative trait locus (QTL) mapping in an F2 cross identified a chromosome 1 QTL explaining 7-12% of the variance in OXY locomotion and anxiety-like withdrawal in the elevated plus maze. A second QTL for EPM withdrawal behavior on chromosome 5 near *Gabra2* (alpha-2 subunit of GABA-A receptor) explained 9% of the variance. To narrow the chromosome 1 locus, we generated recombinant lines spanning 163-181 Mb, captured the QTL for OXY locomotor traits and withdrawal, and fine-mapped a 2.45-Mb region (170.16-172.61 Mb). Transcriptome analysis identified five, localized striatal cis-eQTL transcripts and two were confirmed at the protein level (KCNJ9, ATP1A2). *Kcnj9* codes for a potassium channel (GIRK3) that is a major effector of mu opioid receptor signaling. *Atp1a2* codes for a subunit of a Na+/K+ ATPase enzyme that regulates neuronal excitability and shows functional adaptations following chronic opioid administration. To summarize, we identified two candidate genes underlying the physiological and behavioral properties of opioids, with direct preclinical relevance to investigators employing these widely used substrains and clinical relevance to human genetic studies of opioid use disorder.

## INTRODUCTION

The high prevalence of opioid use disorder (**OUD**) in the United States was fueled by overprescribing and misuse of the highly addictive mu opioid receptor agonist oxycodone (**OXY**; active ingredient in Oxycontin®) (Kibaly *et al*. 2021). The switch to an abuse-deterrent formulation led to a surge in illicit use of heroin (Cicero & Ellis 2015) and now fentanyl (Perdue *et al*. 2024). Opioid-related deaths surpassed 100,000/year (U.S. CDC, 2023; https://www.cdc.gov/opioids/basics/epidemic.html). Twin studies estimate heritability of OUD is ∼50% (Ho *et al*. 2010; Ducci & Goldman 2012), however the genetic basis remains largely unknown (Gelernter & Polimanti 2021; Deak *et al*. 2022; Gaddis *et al*. 2022). Both shared and distinct genetic factors and biological mechanisms likely contribute to various neurobehavioral responses and adaptations associated with opioid use disorder, including initial neurobehavioral and subjective sensitivity, tolerance, withdrawal, and agonist-induced alleviation of withdrawal.

Genetic factors influencing drug behavioral sensitivity can predict additional model behaviors for OUD. Sensitivity to locomotor stimulation induced by addictive drugs is a behavior mediated by shared neural circuitry and neurochemistry with reward and reinforcement, namely mesostriatal dopamine release (Wise & Bozarth 1987; Di Chiara & Imperato 1988; Adinoff 2004). In support of shared genetics between acute drug sensitivity and other model traits for opioid use disorder, we mapped quantitative trait loci (**QTLs**) and candidate genes underlying sensitivity to acute psychostimulant- and opioid-induced locomotor activation (e.g., *Csnk1e*, *Hnrnph1*) and confirmed their involvement in drug reward (e.g., conditioned place preference; **CPP**) and operant reinforcement (e.g., self-administration) (Bryant *et al*. 2012, 2021; Yazdani *et al*. 2015; Goldberg *et al*. 2017; Ruan *et al*. 2020).

C57BL/6J(**B6J**) and C57BL/6NJ(**B6NJ**) are two common substrains in biomedical research and are nearly genetically identical, yet exhibit differences in model traits for substance use disorders (Bryant *et al*. 2008), including ethanol consumption (Mulligan *et al*. 2008; Warden *et al*. 2020; Jimenez Chavez *et al*. 2021), psychostimulant locomotion (Kumar *et al*. 2013; Goldberg *et al*. 2021a), nicotine-induced locomotor activity, hypothermia, antinociception, and anxiety-like behavior (Akinola *et al*. 2019), naloxone-induced conditioned place aversion (Kirkpatrick & Bryant 2015), and binge-like eating (Kirkpatrick *et al*. 2017; Babbs *et al*. 2020). While phenotypic differences between B6 substrains can be quite large (Bryant *et al*. 2008; Matsuo *et al*. 2010; Simon *et al*. 2013), genotypic diversity is extremely small (Keane *et al*. 2011; Yalcin *et al*. 2011; Simon *et al*. 2013; Mortazavi *et al*. 2022; Ferraj *et al*. 2023). Together, these characteristics are ideal for behavioral QTL and expression QTL (**eQTL**) analysis in an F2 reduced complexity cross (**RCC**) and can facilitate gene/variant identification (Bryant *et al*. 2018, 2020). For example, Kumar and colleagues mapped a QTL for cocaine locomotor sensitization to a missense mutation in *Cyfip2* (Kumar *et al*. 2013). We mapped this locus for methamphetamine locomotion (Goldberg *et al*. 2021a) and binge-like eating of sweetened food (Kirkpatrick *et al*. 2017). Mulligan and colleagues used the BXD recombinant inbred strains (a panel of genotyped inbred strains derived from crosses between C57BL/6J and DBA/2J parental strains) combined with analysis of C57BL/6 substrains to identify a functional intronic variant in *Gabra2* (alpha-2 subunit of the GABA-A receptor) that reduced Gabra2 mRNA and GABRA2 protein expression (Mulligan *et al*. 2019). Genetic mapping by our group followed by validation using genetically engineered lines identified the *Gabra2* variant in male-driven methamphetamine locomotion (Goldberg *et al*. 2021a). Phillips and colleagues used reduced genetic complexity between DBA/2J substrains as a confirmation step to identify a functional missense mutation in *Taar1* originally discovered in selected lines derived from C57BL/6J and DBA/2J alleles that influences methamphetamine-induced hyporthermia, aversion, and toxicity (Harkness *et al*. 2015; Shi *et al*. 2016; Miner *et al*. 2017; Reed *et al*. 2017; Phillips *et al*. 2021). We extended RCCs to BALB/c substrains and quickly identified candidate genes for nociceptive sensitivity, brain weight, and brain OXY metabolite levels (Beierle *et al*. 2022a, 2022b).

Here, we mapped the genetic basis of OXY behavioral sensitivity and spontaneous withdrawal in a B6J x B6NJ-F2 RCC. Upon identifying a major distal chromosome 1 QTL for oxycodone locomotion and withdrawal, we implemented an efficient fine mapping strategy using recombinant lines within the QTL interval (Bryant *et al*. 2018), and resolved a 2.45-Mb region. We then sourced a historical striatal expression QTL dataset from the same genetic cross (Bryant *et al*. 2019; Goldberg *et al*. 2021a) to home in on functionally plausible candidate genes. Finally, we conducted immunoblot analysis of eQTL genes within a recombinant line capturing the behavioral QTL, providing further support for the candidacy of *Atp1a2* and *Kcnj9*. The results provide multiple positional loci and highly plausible candidate genes underlying opioid behavioral sensitivity and withdrawal.

## MATERIALS AND METHODS

### Drugs

Oxycodone hydrochloride (**OXY**,Sigma-Aldrich, St. Louis, MO, USA) was dissolved in sterilized physiological saline (**SAL**;0.9%) prior to systemic injections (10 ml/kg, i.p.).

### Parental C57BL/6 substrains and F2 mice

All experiments were conducted in strict accordance with National Institute of Health guidelines for the Care and Use of Laboratory Animals and approved by the Boston University Institutional Animal Care and Use Committee. Details on acquisition of substrains and breeding are provided in **Supplementary Material**.

### Locomotor activity and OXY-CPP (Weeks 1 and 2)

Our nine-day OXY-CPP protocol is published (Kirkpatrick & Bryant 2015). Mice, including the parental substrains, all of the F2 mice, and one cohort of N6-1 mice, were trained in a two-chamber unbiased CPP protocol over nine days (D), with initial preference (D1, SAL i.p.), two alternating pairings of OXY (1.25 mg/kg, i.p.) and SAL (i.p.), separated by 48 h (D2-D5), a consolidation period (D6-D7), a drug-free CPP test (D8), and a state-dependent test for CPP (D9) following SAL (i.p.) or OXY (1.25 mg/kg, i.p.). The 1.25 mg/kg dose of OXY was chosen based on data in the Results section indicating substrain differences in OXY-induced locomotor activation at this dose. See **Supplementary Material** for further details.

### Baseline nociception, acute oxycodone-induced antinociception, and tolerance (Week 3) and elevated plus maze behaviors during spontaneous withdrawal from oxycodone (Week 4)

For Weeks 3 and 4, starting on Monday, a subset of 236 F2 mice (118 SAL, 118 OXY) that went through the OXY-CPP prototocol above were injected once daily with either SAL (i.p.) or OXY (20 mg/kg, i.p.) on Monday through Thursday at 1600 h. Sixteen hours later on Friday, the majority of these 236 F2 mice (160 F2 mice total: 77F, 83M) were tested for baseline nociceptive sensitivity on the 52.5°C hot plate assay and then assessed for OXY-induced antinociception (5 mg/kg, i.p.) at 30 min post-OXY (see **Supplementary Material**). The remaining 76 F2 mice (37F, 39M) received the same OXY regimen during Week 3 but were not exposed to hot plate testing. The 20 mg/kg dose of OXY was chosen for repeated dosing during Week 3 and Week 4 based on prior pilot data indicating antinociceptive tolerance in parental substrains (**Supplementary Material**) and and prior pilot data indicating an effect of OXY on EPM behaviors during withdrawal (data not shown). We report the hot plate procedure above to fully document the history of the mice leading up to the results for the elevated plus maze that are presented in this study (see below). A thermal nociception QTL for hot plate latency (s) and its associated cis-eQTLs are published from these F2 mice (Bryant *et al*. 2019).

For Week 4, we continued injecting all 236 F2 mice (118 SAL, 118 OXY) once daily with SAL (i.p.) or OXY (20 mg/kg, i.p.) for four days. On Day 5, 16 h post-injection, mice were tested for anxiety-like behavior in the elevated plus maze (**EPM**) (Schulteis *et al*. 1998). Each mouse was transported in a clean cage from the vivarium to a testing room with red-illuminating ceiling lights and placed into the center of the EPM (Stoelting Instruments, Wheaton, IL USA) and was video recorded for 5 min. Video recordings were tracked using AnyMaze. The 16 h time point for assessment of EPM behaviors was based on our previous studies with morphine tolerance assessment which ranged from 16-24 h post-final injection, an interval during which we observed baseline- as well as morphine-induced hyperalgesia (Eitan *et al*. 2003; Bryant *et al*. 2006a, 2006b). At the time of designing this OXY study in 2014, there was a very limited number of studies addressing the time course of OXY withdrawal in mice. However, a recent report examining the time course of onset of spontaneous OXY withdrawal in mice identified a peak between 6 h and 24 h which in retrospect, supports our choice of 16 h as a rational time point for assessment of EPM behaviors (Contreras *et al*. 2024). Our primary measure of the emotional-affective component of opioid withdrawal was % Open Arm Time [(open time)/(open time + closed time) *100)]. We also examined Open Arm Entries (#) and Open Arm Distance (m). Twenty-four hours after EPM assessment (Saturday of Week 4), striatum punches (Yazdani *et al*. 2015; Kirkpatrick *et al*. 2017) were collected and processed for RNA-seq analysis (GSE 119719), including eQTL analysis (Bryant *et al*. 2019; Goldberg *et al*. 2021a).

### Behavioral analysis in B6J and B6NJ substrains

Behaviors were analyzed in R (https://www.r-project.org/) or Graphpad Prism Inc. using two- or three-way ANOVAs, with Substrain, Sex, and sometimes Treatment as factors. Furthermore, some analyses include a repeated measure, namely Day or Time (5-min bin). Main effects and interactions (p<0.05) were pursued with Bonferroni or Tukey’s post-hoc tests (as indicated) to determine their sources.

### Genotyping and QTL analysis in B6J x B6NJ-F2 mice

The marker panel comprised 96 markers and is published (Kirkpatrick *et al*. 2017; Bryant *et al*. 2019; Goldberg *et al*. 2021b) (see **Supplementary Material** for details). We first performed Ordered Quantile (ORQ) normalization (Peterson & Cavanaugh 2019) and ran ANOVAs with Sex, Treatment, and Testing Location (data were collected in two different buildings on Boston University Medical Campus). QTL analysis was performed using R/qtl (Broman *et al*. 2003; Broman, & Sen 2009) as described (Kirkpatrick *et al*. 2017; Bryant *et al*. 2019; Goldberg *et al*. 2021b) (**Supplementary Material**). QTL models were finalized based on main effects and interactions. For OXY locomotion traits, Sex and Location were included as additive covariates, and Tx as an interactive covariate. For EPM phenotypes, Location was included as an additive covariate, and Tx was included as an interactive covariate. Additional analyses were performed as described below, including, e.g., sex-specific analyses to assess whether QTLs were being driven by females or males and analysis of SAL mice only to further assess the specificity of the QTLs for OXY treatment.

### Interval-specific recombinant lines for fine mapping the distal chromosome 1 QTL for OXY locomotion

We exploited the near-isogenic F2 background and simple genetic architecture of complex traits in RCCs (Kumar *et al*. 2013; Kirkpatrick *et al*. 2017; Bryant *et al*. 2019; Goldberg *et al*. 2021b; Beierle *et al*. 2022a, 2022b) to devise a rapid fine mapping approach. We first backcrossed an F2 mouse heterozygous (J/N) across the distal chromosome 1 QTL interval to parental B6J mice and identified offspring with new recombination events spanning the 163-181 Mb region. We propagated these lines by again backcrossing to B6J and monitoring genotypes to ensure retention of heterozygosity within each sub-interval. New recombination events were similarly propagated as new lines. Great care was taken to ensure lines were maintained separately, even when they appeared to have the same break points. Individuals within and among lines at each generation of backcrossing are equally unrelated outside of the recombinant interval and continual backcrossing avoids population structure among families as the genetic background progresses toward isogenic B6J. Details of recombinant lines (genotyping, lineage, analysis) are provided in **Supplementary Material**.

### Power analysis of required sample size for recombinant lines

To estimate sample size per genotype for recombinant lines to capture the QTL, we used means and standard deviations of oxycodone-induced locomotor activity for J/J versus J/N genotypes at the peak chromosome 1 QTL marker (181.32 Mb). Based on Cohen’s d= 0.79, a sample size of 27 homozygous J/J and 27 heterozygous J/N mice was required to achieve 80% power (2-tailed test, p < 0.05). Therefore, we aimed for n=27 per genotype per recombinant line.

### Behavioral phenotyping of recombinant lines

For N6-1, we generated two cohorts to replicate and fine-map the distal chromosome 1 QTL. For the first cohort (n=26-27 J/J SAL, 30 J/N SAL, 34 J/J OXY, 43-44 J/N OXY), mice were run through the full nine-day OXY-CPP protocol (Monday-Thursday: Weeks 1 and 2). A subset of the N6-1 mice (n=18 J/J SAL, 19-20 J/N SAL), 14 J/J OXY, 23 J/N OXY) that were run through CPP were then run through two additional, four-day weeks (Weeks 3 and 4: Monday-Thursday) of four daily OXY injections (4x 20 mg/kg, i.p.). For Week 3, 16 h post-final OXY injection, mice were assessed baseline nociception and then subsequently for antinociceptive tolerance on the 52.5degC hot plate at 10, 20, 30, and 40 min post-OXY (5 mg/kg, i.p. - see **Supplementary Material** for methodological details). For Week 4, on Friday, 16 h following the final injection of the daily 4x 20 mg/kg OXY regimen (Monday-Thursday), mice were tested on the EPM during OXY withdrawal.

For the second cohort of the N6-1 line and for the seven additional recombinant lines, because robust genetic results were observed on D2 following the first OXY injection, we used an abbreviated, two-day protocol to facilitate fine mapping. Specifically, to precisely replicate the experimental conditions that led to the identification of the distal chromosome 1 QTL, mice received SAL on D1 (i.p.; open access) and OXY on D2 (1.25 mg/kg, i.p.; OXY-paired side). Details of statistical analyses are provided in the **Supplementary Material**.

### Striatal tissue collection from F2 mice, RNA-seq and striatal eQTL mapping

Procedures for RNA extractions, RNA-seq, and eQTL analysis of F2 samples are published (Bryant *et al*. 2019; Goldberg *et al*. 2021b) (see **Supplementary Material** for details). We also conducted Pearson’s correlation between transcript levels (normalized read counts) and OXY behaviors and fit a linear model to regress transcript levels onto behavior as described in **Supplementary Material**. Striatum punches were harvested from 23 F2 mice (7F, 16M, 76-126 days old; all OXY-treated mice who went through the four-week protocol) 24 h after assessment of EPM behaviors. Because we already had the genotypes and information regarding the distal chromosome 1 QTL (including location and knowledge of dominant mode of inheritance), we “cherry-picked” 11 samples that were homozygous J/J (4 females, 7 males) and 12 samples that were heterozygous J/N (3 females, 9 males) throughout the 163-181 Mb region to ensure that we had a balanced sample size of these two critical genotypes.

### Protein analysis of positionally cloned candidate genes

For protein analysis of candidate genes/transcripts, we harvested striatal punches (2.5 mm diameter, 3 mm thick) from 24 experimentally naïve N6-1 mice (all littermates), including 13 J/J (5 females, 8 males) and 11 J/N (3 females, 8 males). Due to the limited sample sizes of the females, data were sex-collapsed for each genotype. Genotypic differences were determined via Student’s t-tests. Because each immunoblot was an independent assessment of a different protein (unlike the recombinant lines assessed for behavior) and because each protein was chosen based on eQTL results that already implemented a false-discovery rate to correct for multiple comparisons, we report proteins as differentially expressed using both unadjusted p-values as well as Bonferonni-adjusted p-values for the seven comparisons. Information on immunoblot procedures is provided in **Supplementary Material**.

### Identification of genetic variants between C57BL/6 substrains

Genomic variation (e.g., SNPs, indels, structural variants) among nine C57BL/6 and five C57BL/10 substrains has recently been documented (Mortazavi *et al*. 2022) and supplementary data tables for this publication are available at Mendeley: (https://data.mendeley.com/datasets/k6tkmm6m5h/3). No homozygous coding variants or structural variants that differentiate J and NJ were found within this region. Fifteen non-coding homozygous SNPs or InDels were located within the distal Chr1 interval (**Supplementary Material**). There are many heterozygous variants that differentiate substrains in this region, possibly the result of segmental duplication. However, validation and assessment of these variants has been difficult (Mortazavi *et al*. 2022) and thus, heterozygous variants were not considered or reported in the analysis. Variant position was converted from GRCm38 to GRCm39 using the Assembly Converter tool in Ensembl and the Variant Effect Predictor tool (W *et al*. 2016) was used to assess variant effect. Repeating elements that overlapped with variants were annotated using the RepeatMasker track available in the USCS Genome Browser (Nassar *et al*. 2023).

## RESULTS

### Reduced OXY locomotor sensitivity in B6NJ versus B6J substrains

The OXY-CPP protocol is illustrated in **Fig.1A**. We tested for Sex x Substrain interactions for each phenotype. If no interaction, Sex was removed from the model. If there was a significant interaction, we analyzed the sexes separately. In examining OXY locomotion on D2 and D4 alongside SAL counterparts (0 mg/kg), there was a dose-dependent increase in OXY-induced locomotor activity, that, when considering all doses, did not interact with Substrain (**Fig.1B,C**). Nevertheless, we wanted to know if there was a particular dose where the substrains differed on D2 and D4 that would be amenable to genetic analysis. We focused on 1.25 mg/kg OXY, given that it was an intermediate dose for which there was evidence for a substrain difference. For both D2 and D4, two-way ANOVA with the inclusion of the SAL group (0 mg/kg) vs. 1.25 mg/kg OXY in the model indicated a Genotype x Treatment interaction that was explained by decreased OXY-induced locomotor activity in B6NJ versus B6J, with no difference between SAL groups (i.p., **Fig.1D,E**). Importantly, we also examined Substrain and Dose effects on SAL training days for OXY-CPP (D3, D5) and found little evidence for substrain differences, although there was evidence for a conditioned locomotor effect at the two highest prior OXY doses (**Fig.S1**). These results support decreased OXY locomotion in B6NJ versus B6J mice, providing the impetus for forward genetic analysis. Accordingly, narrow-sense h2 estimates (Hegmann & Possidente 1981) based on between-strain (genetic), within-strain (environmental), and total variance were 0.22 and 0.31 for D2 and D4 OXY locomotion (1.25 mg/kg, i.p.), respectively.

**Figure 1.**
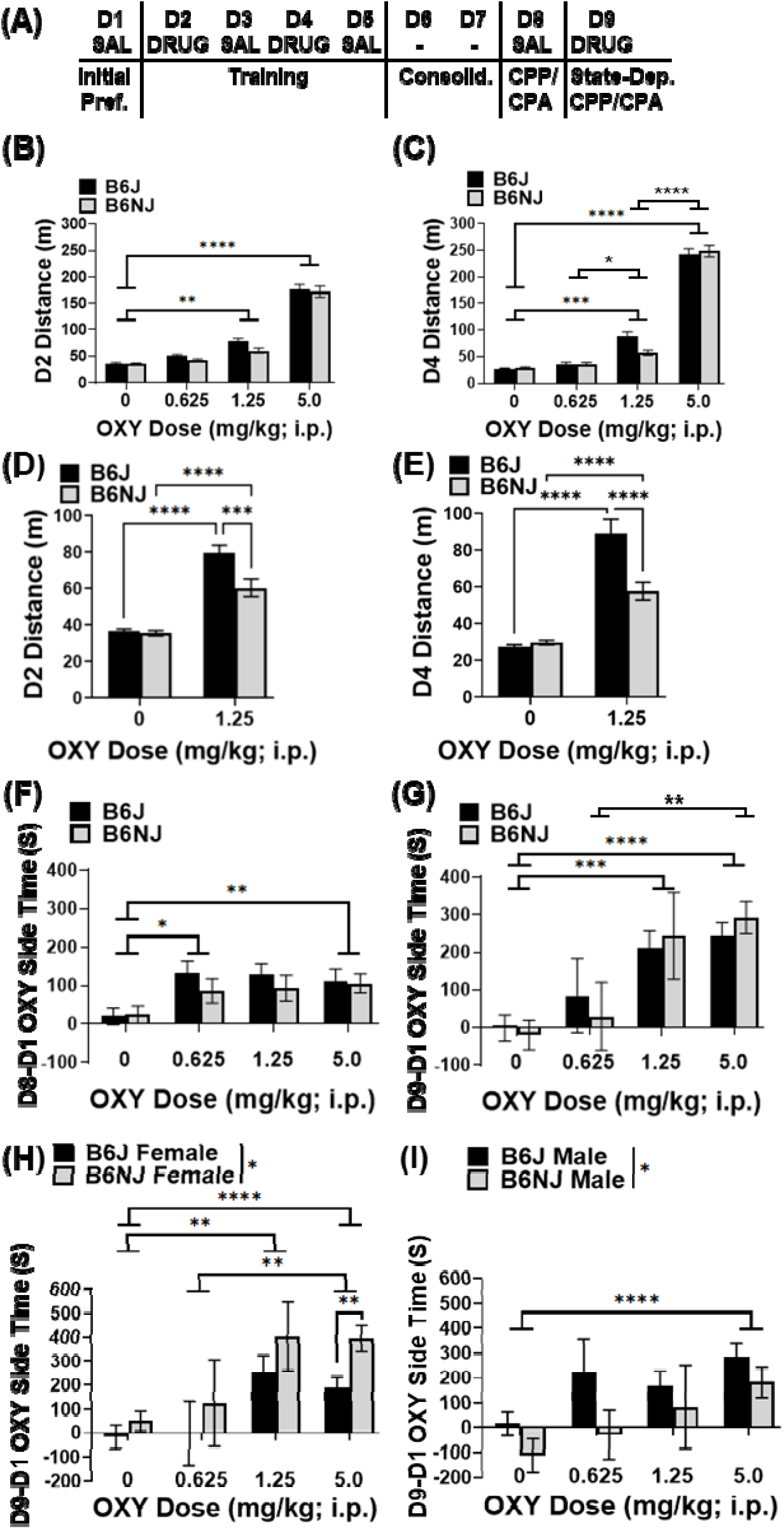
OXY-induced locomotor activity and conditioned place preference in B6J vs. B6NJ substrains. **(A):** Schematic of OXY-CPP protocol. D=Day. For all phenotypes below with the exception of state-dependent CPP (panel G), inclusion of Sex in the model revealed no effect of Sex or interactions with Genotype (ps>0.05); thus, data were collapsed across sexes. **(B):** Two-way ANOVA indicated a Dose effect (F3,293=176.2; p<0.0001) that was explained by an increase at 1.25 vs. 0 mg/kg (**p<0.01) and at 5.0 vs. 0 mg/kg (****p<0.0001) (Bonferroni). There was no Substrain effect (F1,293=1.66; p=0.20) and no interaction (F3,293<1). Unpaired t-test between Genotypes at the 1.25 mg/kg dose indicated a significant decrease in OXY distance in B6NJ vs. B6J (t30=2.84; p=0.008), prompting an additional analysis with 0 mg/kg (SAL) in panel D. **(C):** Two-way ANOVA indicated a Dose effect (F3,291=386.2; p<0.0001) that was explained by an increase at 1.25 vs. 0 mg/kg (***p<0.001), 5.0 vs. 0 mg/kg (****p<0.0001), 1.25 vs. 0.625 mg/kg (*p < 0.05), and 5.0 vs. 1.25 mg/kg (****p<0.0001) (Bonferroni). There was no Substrain effect (F1,291<1) and no interaction (F3,291=1.11; p=0.35). Unpaired t-test between Genotypes at the 1.25 mg/kg dose indicated a significant decrease in OXY distance in B6NJ vs. B6J (t30=2.91; p=0.0068), prompting an additional analysis with 0 mg/kg (SAL) in panel E. **(D):** In narrowing the focus of D2 distance to 1.25 mg/kg OXY vs. 0 mg/kg (SAL), two-way ANOVA indicated an effect of Substrain (F1,138=14.67; p<0.001), Treatment (F1,138=167.9; p <0.0001), and a Substrain x Treatment interaction (F1,138=12.15; p<0.001). Both substrains showed an increase in distance traveled following OXY vs. SAL (****p<0.0001) (Tukey’s). Furthermore, B6NJ showed a significant decrease in OXY-induced distance traveled compared to B6J (***p<0.001), with no difference between SAL groups (p>0.05) (Tukey’s). **(E):** In narrowing the focus of D4 distance to SAL vs. 1.25 mg/kg OXY for D4, there was an effect of Substrain (F1,136=20.23; p<0.0001), Treatment (F1,136=189.6; p<0.0001), and an interaction (F1,136=26.80; p<0.0001). Both substrains showed an increase in distance traveled following OXY vs. SAL (***p<0.0001) (Tukey’s). Furthermore, B6NJ showed a significant decrease in OXY-induced distance traveled compared to B6J (****p<0.0001), with no difference between SAL groups (p>0.05). **(F):** For drug-free OXY-CPP, there was a Dose effect (F3,293=5.52; p< 0.01) that was explained by increased OXY-CPP at 0.6125 vs. 0 mg/kg (*p<0.05) and at 5 vs. 0 mg/kg (**p<0.01) (Bonferroni). There was no Substrain effect (F3,293<1) and no interaction (F3,293<1). **(G):** For state-dependent OXY-CPP, two-way ANOVA indicated a Dose effect (F3,269=17.56; p<0.0001) that was explained by increased OXY-CPP at 1.25 vs. 0 mg/kg (***p<0.001), 5 vs. 0 mg/kg (****p<0.0001) and 5 vs. 0.6125 mg/kg (**p<0.01) (Bonferroni). There was no Substrain effect (F1,269<1) and no interaction (F3,269<1). (**H**, **I**): Inclusion of Sex in the ANOVA model for state-dependent CPP revealed a Dose effect and a Substrain x Sex interaction. Females-only analysis indicated a Dose effect (F3,134=11.11; p<0.0001) that was explained by an increase at 1.25 vs. 0 mg/kg (**p<0.01), at 5 vs. 0 mg/kg (****p<0.0001), and at 5 vs. 0.6125 mg/kg (*p<0.05) (Bonferroni) and Substrain effect (F1,134=5.59; p<0.05) that was explained by a significant increase in OXY-CPP in B6NJ vs. B6J at the 5.0 mg/kg OXY dose (**p<0.01) (Bonferroni). Males-only analysis indicated a Dose effect (F3,127=7.88; p<0.0001) that was explained by a significant increase in preference at 5 vs. 0 mg/kg (****p<0.0001) (Bonferroni) and a Substrain effect (F1,127=5.53; p<0.05) that was not accounted for by any specific dose (ps>0.05). Sample sizes include the following: B6J 0 mg/kg: n=56-60 (30F, 26M-30M); B6NJ 0 mg/kg: n=46-50 (26F-30F, 19M-20M); B6J 0.6125 mg/kg: n=16-24 (n=10F-14F, 6M-10M); B6NJ 0.6125 mg/kg: n=16-24 (6-10F, 10-14M); B6J 1.25 mg/kg: n=20 (10F, 10M); B6NJ 1.25 mg/kg: n=12 (6F, 6M); B6J 5 mg/kg: n=55 (25F, 30M): n=55; B6NJ 5 mg/kg n=56 (29F, 27M).

In examining drug-free OXY-CPP as the difference between final (D8) and initial preference (D1; D8-D1 OXY Side Time, s), there was a significant Dose effect that was explained by a significant CPP at the lowest and highest prior OXY doses (**Fig.1F**). For state-dependent CPP (D9-D1 OXY Side Time, s), sex-collapsed analysis also indicated a Dose effect that was explained by a dose-dependent increase in CPP (**Fig.1G**). Furthermore, inclusion of Sex in the model revealed a Sex x Substrain interaction that prompted separate analyses in females and males. For females, there was a Dose effect that was explained by an increased preference at 1.25 mg/kg and 5.0 mg/kg relative to 0 mg/kg (SAL) and by an increase in preference from 0.625 mg/kg to 5.0 mg/kg (**Fig.1H**). There was also a Substrain effect that was explained by increased preference in B6NJ versus B6J mice at 5.0 mg/kg (**Fig.1H**). For males, there was also a Dose effect that was explained by a significant increased preference at 5.0 mg/kg versus 0 mg/kg (SAL) (**Fig.1I**). The Substrain effect was due to an overall decrease in preference in B6NJ versus B6J, irrespective of the dose (**Fig.1I**).

### QTL mapping of OXY locomotion (Week 1)

A list of genetic markers is provided in **Table S1**. The timeline of OXY regimen, behavioral assessment, and sample sizes are provided in **Fig.2A**. Starting with 425 F2 mice (213 SAL, 213 OXY), we mapped a genome-wide significant QTL on distal chromosome 1 for D2 and D4 OXY locomotor traits using a treatment-interactive model, where both treatments were included in a single analysis (**Fig.2B-D**). Like the parental substrains, mice containing 1-2 copies of the B6NJ allele (J/N, N/N) at the peak-associated marker showed a decrease in D2 OXY-induced distance traveled (**Fig.2E**) as well as D4 distance and other D2 and D4 OXY locomotor traits (**Fig.S2**), with no change in D2 or D4 distance traveled in SAL-treated mice as a function of Genotype (**Fig.2E; Fig.S2**). These observations support a dominant inheritance of the N allelic effect and provide strong support for a causal gene/variant within this locus underlying OXY locomotion. To address specificity of this QTL for OXY treatment, QTL analysis of D3 and D5 locomotor traits (SAL treatment days) indicated that while there was a trending QTL peak for distal chromosome 1 on D3, it was not significant (**Fig.S3A**) and is likely mediated by a carry-over, conditioned locomotor response from the first OXY injection (**Fig.S1**). Additionally, QTL analysis of D1 distance revealed no genome-wide significant QTLs (**Fig.3SB**). Finally, QTL analysis of only the SAL F2 mice revealed no genome-wide significant QTLs for D2 or D4 locomotor traits (**Fig.S3C**), providing further evidence for specificity of the distal chromosome 1 for OXY-induced behaviors.

**Figure 2.**
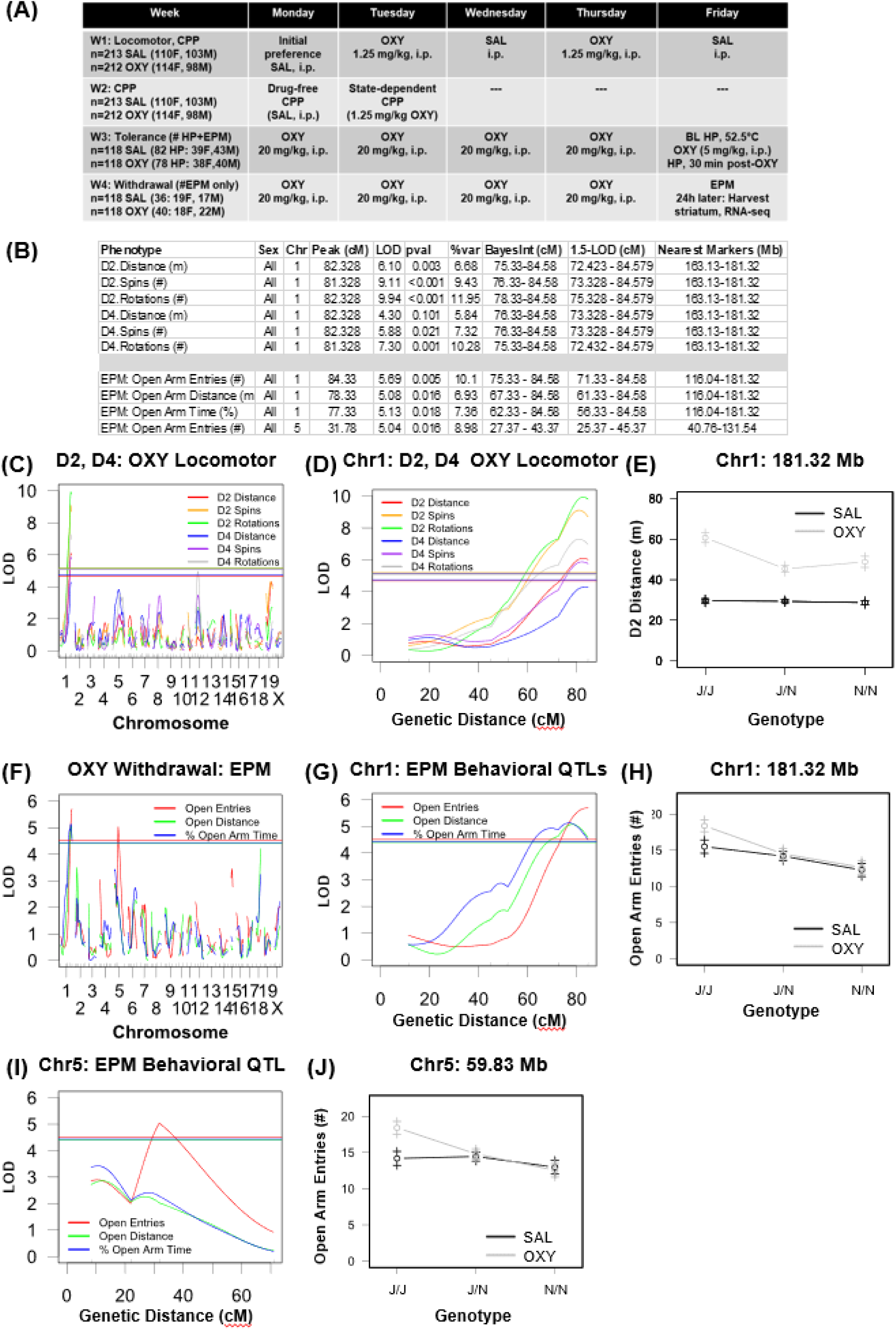
QTLs on chromosomes 1 and 5 for OXY locomotion and spontaneous withdrawal. **(A):** Summary of 4-week (W) protocol for F2 mice (sample sizes as indicated) and the N6-1 recombinant line (cohort in Fig.3). **(B):** Summary of QTL results for six OXY-induced locomotor traits and three EPM withdrawal traits during spontaneous OXY withdrawal. **(C):** Genome-wide plot of D2 and D4 OXY locomotor traits (Distance, Spins, Rotations) following 1.25 mg/kg OXY (i.p.) during CPP training. **(D):** Chromosome 1 QTL plot for D2 and D4 OXY locomotor traits. **(E):** Chromosome 1 effect plot for D2 OXY Distance (see Fig.S2 for additional effect plots). **(F):** Genome-wide QTL plot of EPM traits during spontaneous OXY withdrawal. **(G):** Chromosome 1 QTL plot for EPM withdrawal traits. **(H):** Chromosome 1 effect plot at peak-associated marker for Open Arm Entries (#) (see Fig.S5 for additional effect plots). **(I):** Chromosome 5 plot for Open Arm Entries (#)**. (J):** Chromosome 5 effect plot at peak-associated marker for Open Arm Entries. Sample sizes for F2 mice are as follows: D2, D4 SAL: n= 213 (110F, 103M). D2, D4 OXY: n=212 (114F, 98M). EPM SAL: n=118 (58F, 60M). EPM OXY: n=118 (56F, 62M).

**Figure 3:**
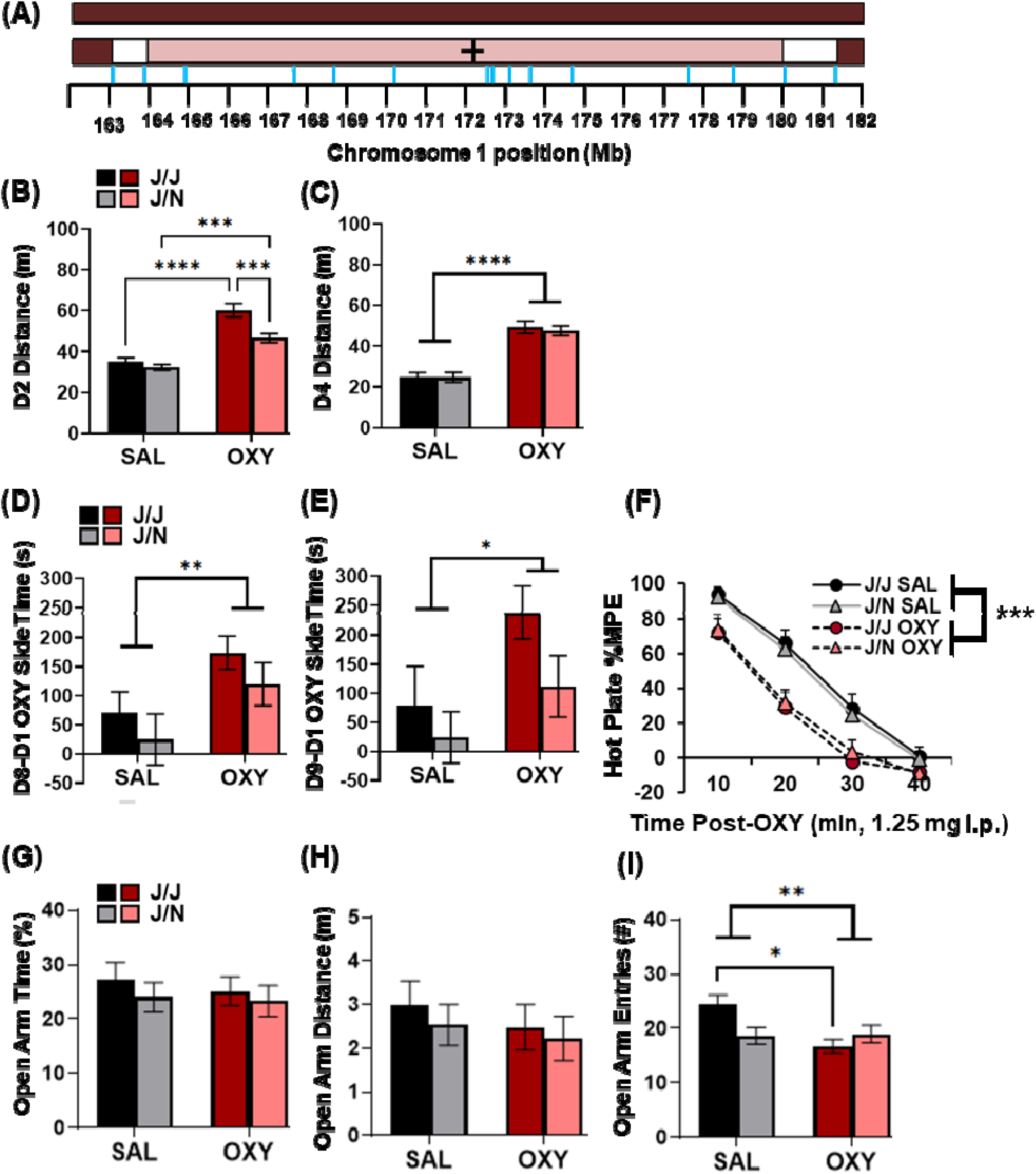
Capture of the distal chromosome 1 QTL for OXY locomotion and withdrawal in the N6-1 recombinant line. **(A):** Schematic of the N6-1 recombinant interval spanning 163-181 Mb. Blue ticks on the x-axis indicate marker location (Mb). Burgundy color = homozygous J/J. Mauve color = heterozygous J/N. White color = region of uncertainty for recombination breakpoint. **(B):** For D2 locomotor activity (OXY), three-way ANOVA revealed no effect of Sex or interactions of Sex with Genotype (ps >0.05); thus, data were collapsed across sexes. There was an effect of Genotype (F1,31=11.36; p<0.01), Treatment (F1,131=63.66; ***p<0.0001), and an interaction (F1,131=4.56; p<0.05). There was a decrease in OXY-induced locomotor activity in J/N versus J/J (***p<0.001) (Bonferroni). Both J/J and J/N OXY groups showed an increase in locomotor activity versus their SAL counterparts (****p<0.0001,***p<0.001, respectively) (Bonferroni). **(C):** In examining D4 OXY locomotion, three-way ANOVA revealed no effect of Sex or interactions of Sex with Genotype, so data were collapsed across sexes. There was a Treatment effect (F1,131=80.00; ****p<0.0001) but no Genotype effect (F1,131<1) and no interaction (F1,131<1). **(D):** In examining drug-free OXY-CPP (D8-D1,s), there was no effect of Sex or interactions with Sex, so data were collapsed across sexes. There was a Treatment effect (F1,130=6.91;**p<0.01) but no Genotype effect (F1,130=1.76;p=0.19) and no interaction (F1,130<1). **(E):** In examining state-dependent OXY-CPP (D9-D1,s), there was no effect of Sex or interactions, so data were collapsed across sexes. There was a Treatment effect (F1,131=5.23; *p<0.05) but no Genotype effect (F1,131=2.84; p=0.095) and no interaction (F1,131<1), together indicating significant drug-free and state-dependent OXY-CPP versus SAL, irrespective of Genotype. **(F):** In examining antinociceptive tolerance following a challenge dose of OXY (1.25 mg/kg, i.p.) in a subset of the same mice as used for CPP, there was a main effect of Treatment (F1,268=41.97; ***p<0.0001) and Time (F3,268=133.54; p<0.0001), but no effect of N6-1 Genotype or Sex (F1,268<1), and no interactions of Genotype or Sex with any other factors, indicating significant tolerance, irrespective of N6-1 Genotype or Sex. **(G-I):** In examining EPM behaviors during OXY withdrawal in the same mice as used for CPP->tolerance, there were no effects of Sex and no Genotype x Sex interactions; thus, data were collapsed across sexes. For % Open Arm Time and Open Arm Distance (mc), there was no effect of Genotype (F1,70<1), Treatment (F1,70<1), or interaction (F1,70<1; **panel G,H**). For Open Arm Entries, while there was no Genotype effect (F1,70=1.10; p=0.30), there was a Treatment effect (F1,70=4.90; *p=0.021) and a Genotype x Treatment interaction (F1,70=5.79; p=0.015) that was driven by a decrease in Open Arm Entries in OXY J/J mice vs SAL J/J (*p<0.05 **panel I**), with no significant difference between J/N treatment groups (p>0.05) (Bonferroni). Sample sizes for N6-1 genotypes for panels B-E are as follows: J/J SAL: n=26-27 (17F, 9M-10M); J/N SAL: n=30 (15F, 15M); J/J OXY: n=34 (13F, 21M); J/N OXY: n=43-44 (18F, 25-26M). Sample sizes for N6-1 genotypes for panels F-I (subset of mice used in panels B-E) are as follows: J/J SAL: n=18 (7F, 11M); J/N SAL: n=19-20 (8-9F, 11M); J/J OXY: n=14 (7F, 7M); J/N OXY: n=23 (12F, 11M).

### QTL mapping of OXY withdrawal behaviors in the EPM (Week 4)

Next, a subset (N=236) of the 425 F2 mice underwent repeated, daily i.p. SAL (n=118; 58F, 60M) or high-dose OXY treatment (20 mg/kg, i.p.; n=118; 56F, 62M; **Fig.2A**) on Monday-Thursday of Weeks 3 and 4 (“4x 20 mg/kg”). The majority of these 236 F2 mice (N=160; **Fig.2A**) were examined for antinociceptive tolerance on Day 5 (Friday) of Week 3, while the remaining 76 mice (**Fig.2A**) were only exposed to the 4x 20 mg/kg OXY regimen during Weeks 3 and 4 and not to hot plate testing prior to assessment of EPM withdrawal behaviors on Day 5 (Friday) of Week 4. The 4x 20 mg/kg OXY regimen was chosen based on initial observations of significant antinociceptive tolerance in the C57BL/6 substrains (**Fig.S4A**) and induce significant effects on EPM behaviors during spontaneous withdrawal in previous pilot studies in naïve C57BL/6 substrains (data not shown).

For Week 3, the 160 F2 mice (77F, 83M) that underwent hot plate testing first underwent four days of daily i.p. SAL (39F, 43M) or 20 mg/kg OXY (38F, 40M) and 16 h later on on Day 5 (Friday), following baseline hot plate nociceptive assessment, mice were injected with a challenge dose of 5 mg/kg OXY and assessed for OXY antinociception 30 min later. However, in F2 mice, there was no significant effect of Treatment, Sex, or interaction on OXY antinociception (**new Fig.S4B**), indicating no tolerance and no sex-dependent effects. Possible sources of discrepancy could be due to the fact that the parental substrains from the pilot study had no prior exposure to any experimental procedures such as CPP, injections, etc. whereas F2 mice did. Indeed, the %MPE was much lower in F2 mice versus the parental substrains, suggesting that prior experimental training/handling reduced OXY-induced antinociception (e.g., via habituation to concomitant stress-induced antinociception or cross-tolerance of stress-induced antinociception with OXY antinociception).

For assessment of EPM withdrawal behaviors at the end of Week 4, all 160 F2 mice that underwent hot plate testing at the end of Week 3 resumed the 4x 20 mg/kg OXY regimen during Week 4 (Monday-Thursday) and EPM testing 16 h later on Friday. Furthermore, an additional 76 F2 mice (36 SAL, 40 OXY; **Fig.2A**) that received the Week 3 OXY regimen but not hot plate exposure also resumed the 4x 20 mg/kg OXY regimen during Week 4 (Monday-Thursday) and were tested on the EPM on Friday, thus yielding a total of 236 F2 mice (118 SAL, 118 OXY) for assessment of EPM traits during OXY withdrawal.

Means and S.E.M. of the EPM traits in SAL- and OXY-trained mice is provided in **Fig.S5A-C**. Overall there was no Treatment effect. Nevertheless, segregation of alleles let to the identification of a similarly localized, genome-wide significant, treatment-interactive distal chromosome 1 QTL for EPM withdrawal traits (Open Arm Entries, Open Arm Distance, % Open Arm Time; **Fig.2F-G**), suggesting a common causal gene/variant for D2 and D4 OXY locomotor traits and EPM withdrawal traits. Again, like with OXY locomotor traits, the N genotype at the peak-associated marker (181.32 Mb) was less responsive to OXY as indicated by the effect plot, with only the J genotype showing an increase in Open Arm Entries versus SAL (**Fig.2H**) as well as Open Arm Distance (m) and Open Arm Time (%) (**Fig.S5D,E**). Analysis of only the F2 mice that had been exposed to the hot plate revealed the same distal chromosome 1 locus for Open Arm Time (%) and a trending medial chromosome 5 QTL for Open Arm Entries (#) (**Fig.S6**), indicating that inclusion of the small subset of 76 F2 mice that were not exposed to the hot plate (the addition of which strengthened the already significant LOD scores) did not confound the results.

We mapped a second QTL on medial chromosome 5 for Open Arm Entries (**Fig.2F,I**). Again, OXY-withdrawn mice with the J/J allele at the peak-associated marker (59.83 Mb) showed an increase in Open Arm Entries, while the N allele was unresponsive to OXY (**Fig.2J**). Removal of the 76 F2 mice who were not exposed to the hot plate reduced LOD score of the QTL to a non-significant peak (**Fig.S6**), again supporting the contention that these mice simply added power to detect the QTL and that lack of hot plate exposure did not confound the results. The chromosome 5 QTL is near *Gabra2* (codes for alpha-2 subunit of GABA-A receptor), a well-known cis-eQTL mediated by a single nucleotide intronic deletion near a splice acceptor site that is private to B6J (Mulligan *et al*. 2019) that is genetically linked to and validated for methamphetamine stimulant sensitivity (Goldberg *et al*. 2021a) and multiple seizure models (Hawkins *et al*. 2021; Yu *et al*. 2022). *Gabra2* is a high-priority candidate gene for opioid withdrawal, as the alpha-2 subunit contributes to GABA-A receptor-mediated anxiogenesis and anxiolysis, alcohol dependence, and polydrug misuse (Engin *et al*. 2012). Analysis of SAL-treated F2 mice only revealed no genome-wide significant QTLs for EPM traits (**Fig.S3D**), providing evidence for the specificity of the chromosome 1 and chromosome 5 QTLs for OXY withdrawal.

### Separate QTL analyses of OXY locomotor and EPM withdrawal traits in females and males

In addition to including Sex as an additive covariate in the above analyses, we also performed QTL mapping separately in F2 females and F2 males for D2 and D4 OXY locomotor traits and EPM withdrawal traits using the same QTL model (with Sex removed as an additive covariate). Interestingly, the genome-wide significant QTL at distal chromosome 1 was significant in females but not males (**Fig.7A-E**), indicating that females clearly drove the original sex-combined distal chromosome 1 QTL signal (**Fig.2B-E**). The effect plots for D2 and D4 OXY locomotor traits indicate a larger effect for females versus males (**Fig.S8A-F**).

Sex-specific QTL analysis of EPM behaviors indicated that neither females nor males showed a significant genome-wide significant QTL on distal chromosome 1, though females showed a higher LOD score for chromosome 1 (**Fig.S7F,G**). Sex-specific analysis of the medial chromosome 5 QTL for EPM Open Arm Entries indicated that it was only significant in males (**Fig.S7G,I**), again with only the J allele being responsive to OXY (**Fig.S8G**).

### Distal chromosome 1 recombinant lines spanning 163-181 Mb

Because the distal chromosome 1 QTL for D2 and D4 OXY locomotor traits was large and spanned ∼20 Mb, we sought to rapidly narrow this locus by breeding and phenotyping interval-specific recombinant lines spanning 163-181 Mb at each generation of backcrossing to B6J (Bryant *et al*. 2018). Details of recombinant lines, including genetic markers, their recombination break points, and their pedigree/lineage are described in **Supplementary Material** (see description along with **Tables S2,S3; Fig.S9**).

### N6-1 recombinant line captures distal chromosome 1 QTL for OXY locomotion and withdrawal

A schematic for the recombinant region in the N6-1 line is illustrated in **Fig.3A**. The heterozygous (J/N) region spanned 163.13-181.32 Mb, meaning N6-1 mice were homozygous J/J at these coordinates and beyond. For D2 but not D4, N6-1 showed a significant decrease in OXY locomotion (**Fig.3B-C**). For drug-free and state-dependent OXY-CPP, N6-1 trended toward a reduction, although the results were not statistically significant (**Fig.3D,E**).

In examining antinociceptive tolerance in the same mice at the end of the Week 3 regimen (4x 20 mg/kg OXY, i.p.) and following a challenge dose of 5 mg/kg OXY (i.p.), significant tolerance was evident regardless of Genotype (**Fig.3F**).

For OXY withdrawal on the EPM in the same mice, there was no effect of OXY or Genotype for % Open Arm Time or Open Arm Distance (**Fig.3G,H**); however, for Open Arm Entries there was a Treatment x Genotype interaction that was explained by the OXY-responsive J/J Genotype showing a *decrease* in open arm entries, with no response to Treatment in the J/N Genotype (**Fig.3I**).

### Fine mapping the distal chromosome 1 behavioral QTL for D2 OXY locomotion in recombinant lines

Given there were eight recombinant lines (**Fig.4A; Table S3; Fig.S9**), to expedite fine mapping, we employed a truncated, two-day protocol (D1,D2; **Fig.1A**), focusing on D1 locomotor traits following SAL (i.p.) and open access to the two sides of the CPP chamber and D2 OXY locomotor traits following 1.25 mg/kg (i.p.) of OXY (**Fig.2A-E**). Student’s t-tests with a Bonferroni-corrected p-value were first employed for D1 and D2 locomotor to adjust for comparisons across eight lines (p<0.05/8=0.00625). In separate analyses, we also incorporated Sex with two-way ANOVAs and report Genotype x Sex interactions and post-hoc analyses from in **Fig.S10**. Notably, Genotype x Sex interactions were only observed for D1 locomotor traits (SAL) and not for D2 (OXY) traits]. See **Figure Legends** and **Supplementary Material** for additional details regarding two-way ANOVA results.

**Figure 4.**
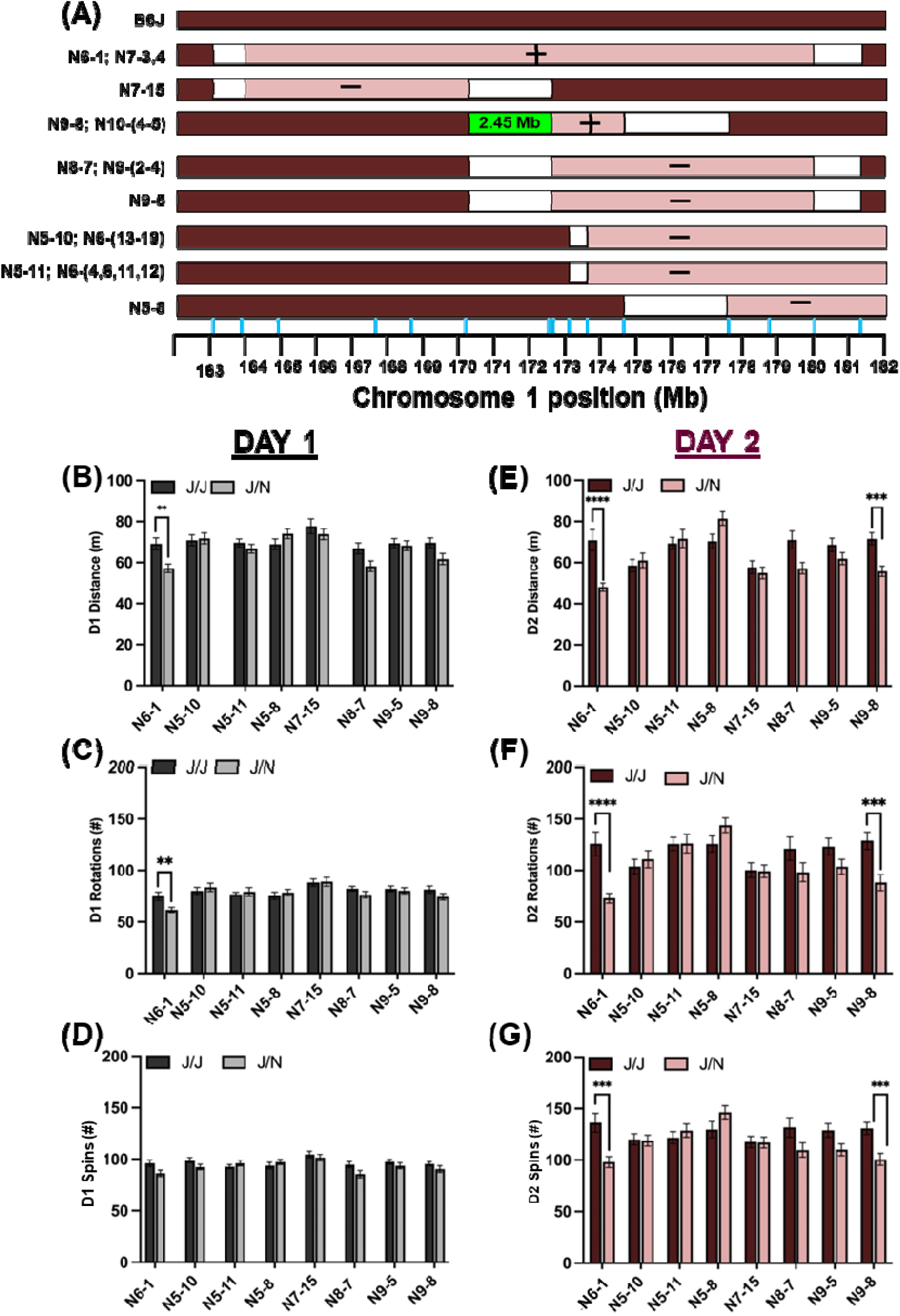
Positional cloning of a 2.45 Mb region on distal chromosome 1 underlying OXY locomotor traits in recombinant lines backcrossed to B6J. **(A):** Schematic of the eight congenic lines that were phenotyped to deduce a 2.45 Mb region spanning 170.16-172.61 Mb. Blue ticks on the x-axis indicate marker location (Mb). Burgundy color: homozygous J/J genotype. Mauve color: heterozygous J/N genotype. White color = region of uncertainty for recombination breakpoint. “+” = captured the QTL for reduced D2 OXY distance. “-” = failed to capture the QTL for reduced D2 OXY distance. **(B-D, vertically):** D1 Distance, D1 Rotations, and D1 Spins following SAL (i.p.) in eight recombinant lines. **p<0.00625. For D1 Distance (SAL, i.p.) in **N6-1**, J/N showed a decrease (t48=3.35; **p=0.0016; **panel B**). For D1 Rotations in **N6-1**, J/N showed a decrease (t48=2.87; **p=0.0061; **panel C**; vertically below panel B). For D1 Spins, none of the lines showed a Genotype effect (ps > 0.038; **panel D**). **(E-G, vertically):** For D2 OXY locomotion, J/N from **N6-1** and **N9-8** showed a decrease (t48=4.29; ****p<0.0001, t49=4.09;***p=0.0002; **panel E**). For D2 OXY rotations, J/N from **N6-1** and **N9-8** showed a decrease (t48=4.36; ****p<0.0001; t49=3.54;***p=0.0009; **panel F**; below panel E). For D2 OXY spins, J/N from **N6-1** and **N9-8** showed a decrease (t48=3.65; ***p=0.0006, t49=3.60; ***p=0.0007; **panel G**; vertically below panel F). Sample sizes for genotypes are as follows: **N6-1:** n=24 J/J (12F, 12M), 24 J/N (9F, 15M); **N5-10:** n=31 J/J (13F, 18M), 26 J/N (15F, 11M); **N5-11:** n=43 J/J (15F, 28M), 29 J/N (9F, 20M); **N5-8:** N=23 J/J (14F, 9M), 30 J/N (14F, 16M); **N7-15:** n=35 J/J (23F, 12M), 26 J/N (11F, 15M); N8-7 (N9-2,3,4): n=22 J/J (12F, 10M), 24 J/N (12F, 12M); **N9-5:** n=25 J/J (14F, 11M), 43 J/N (20F, 23M); **N9-8 (N10-4,5):** n=24 J/J (10F, 14M), 27 J/N (10F, 14M).

For D1 locomotion measures (SAL, i.p.), only N6-1 showed a reduction in Distance and Rotations, and none of the lines showed a difference in Spins (**Fig.4B-D**). For D2 (OXY, 1.25 mg/kg i.p.), both N6-1 and N9-8 showed significant reductions in Distance, Rotations, and Spins (**Fig.4E-G**). In considering the recombination breakpoints of N9-8 that captured the QTL for D2 OXY traits (“**+**”) versus the six congenic lines that did not capture the QTL (“**-**“: N5-10, N5-11, N5-8, N7-15, N8-7, and N9-5), the N9-8(+) J/N segment ended proximally at 172.61 Mb and the distal breakpoint of N7-15 line (-) was 170.16 Mb (**Fig.4A**). These key observations, combined with multiple distal recombinant lines that failed to capture the QTL (N5-10,-11,-8) reduced the causal locus to a 2.45-Mb region (170.16-172.61 Mb; **Fig.4A**). Note that although N8-7(-) and N9-5(-) appear to possess the same proximal breakpoint as N9-8(+) (**Fig.4A**), these two lines comprise two different, independent recombination events from N6-1 (**Fig.S4A; Table S3**) which in turn, differ from the N9-8(+) proximal breakpoint. A snapshot of the genes within the 2.45 Mb interval from UCSC genome browser is provided in **Fig.S11**.

### Striatal cis-expression QTLs, Pearson’s correlation, and regression analysis of transcript levels versus behavior

Expression QTLs (eQTLs) provide functional, causal support for quantitative trait genes/variants underlying behavior (Goldberg *et al*. 2021a; Yao *et al*. 2021; Beierle *et al*. 2022a, 2022b). We focused on chromosome 1-localized transcripts (170.16-172.61 Mb) showing a peak association with the peak for OXY locomotion (rs51237371; 181.32 Mb), including Pcp4l1, Ncstn, Atp1a2, Kcnj9, and Igsf9 (**Fig.5A**). We also note three striatal cis-eQTLs just distal to the 2.45-Mb region that could potentially be regulated by one or more noncoding variants within the 2.45-Mb region, including Cadm3, Aim2, and Rgs7 (**Fig.5A**). **Table S4** comprises 58 transcripts showing a peak association at 181.32 Mb. Because of the high linkage disequilibrium in an F2 cross, some of these transcripts are located tens of Mb away from this marker.

**Figure 5.**
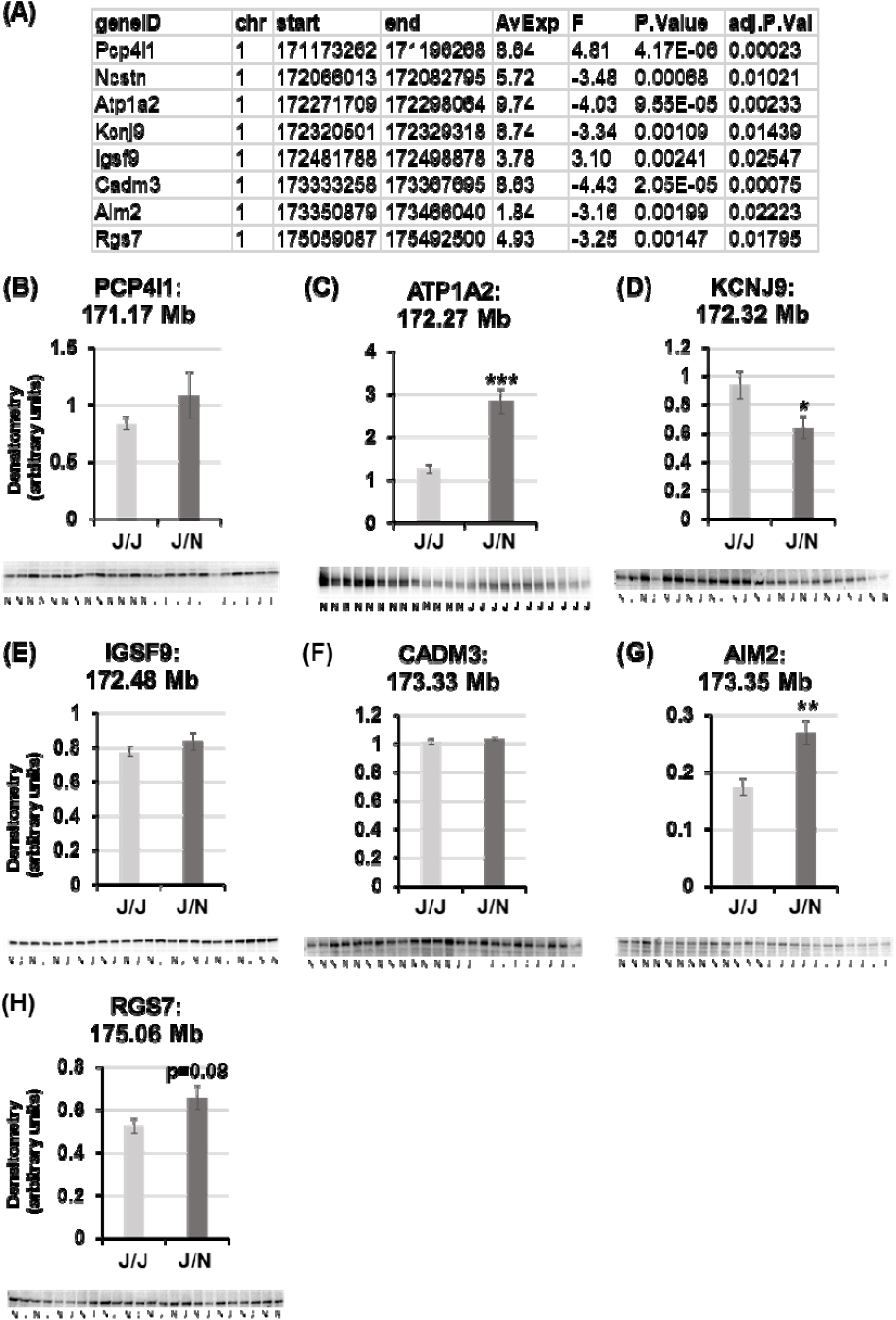
Protein analysis of striatal cis-eQTL transcripts within or distal to the fine-mapped 2.45 Mb region (chromosome 1: 170.16-172.61 Mb) for D2 OXY locomotor traits. **(A):** Striatal cis-eQTL transcripts within the 170.16-172.61 Mb interval (Pcprl1, Atp1a2, Kcnj9, and Igsf9) and just distal to this region (Cadm3, Aim2, and Rgs7). See **Table S4** for the full list of cis-eQTLs showing a peak association with rs51237371 (181.32 Mb). **(B-H):** Immunoblot analysis of proteins coded by candidate genes within the 2.45 Mb region on distal chromosome 1. There was an increase in ATP1A2 immunostaining in J/N (t22=4.59; ***p=0.00014; **panel C**) and a decrease in KCNJ9 immumostaining in J/N (t18=2.21; *p=0.04; **panel D**). There was no difference with PCP4L1 (t22=1.02; p=0.32) or IGSF9 (t22<1; **panel E**). We assayed 3 additional proteins encoded by transcripts with cis-eQTLs that were distal to the fine-mapped 2.45 Mb locus (CADM3, AIM2, RGS7; **panels F-H**). There was an increase in AIM2 in J/N (t22=3.57; **p=0.0017; **panel G**) and a trend for RGS7 (t19=1.80; p=0.088; **panel H;** discussed in **Supplementary Material**). There was no genotypic difference with CADM3 (t22=1.45; p=0.16; **panel F**). For ATP1A2, PCP4L1, IGSF9, AIM2, and CADM3: n=11 J/J (3F, 8M); n=13 J/N (5F, 8M). For KCNJ9, n=11J/J (3F, 8M), n=12 J/N (5F, 7M). For RGS7, n=8 J/J (3F, 5M), n=12 J/N (5F, 7M).

We next focused on chromosome 5 cis-eQTLs for OXY withdrawal (**Fig.2B,F,I,J**). **Table S5** comprises a list of transcripts with a peak association with the peak chromosome 5 QTL marker for OXY withdrawal (rs32809545; 59.83 Mb). A block of five eQTL transcripts was localized to less than 20 Mb of the peak marker, including Cpeb2, N4bp2, Gabra4, Pdgfra, and Clock. Notably, the chromosome 5 QTL for OXY withdrawal lies adjacent to the peak cis-eQTL (rs29547790; 70.93 Mb) for Gabra2 expression.

To potentially aid in further prioritizing candidate genes in the fine-mapped, 2.45-Mb interval for D2 OXY distance, we examined the correlation between gene expression and behavior via simple Pearson’s r and via fitting a linear model and regressing gene expression (normalized read counts) onto D2 OXY distance using the limma package (Ritchie *et al*. 2015).). Simple Pearson’s r identified correlations for Pcp4l1 (r=-0.53; p=0.0088), Igsf9 (r=-0.48; p=0.021), and Rgs7: r=+0.58; p=0.0037) whereas Atp1a2 was not quite significant (r=+0.38; p=0.082). However, results of the linear model support all four transcripts (Pcp4l1, Atp1a2, Igsf9, and Rgs7) as predictors of D2 OXY distance (p_adjusted_<0.05).

### Striatal protein analysis in the N6-1 recombinant line supports Atp1a2 and Kcnj9 as candidate genes underlying OXY locomotion and withdrawal

We ran striatal immunoblot analysis from naive N6-1 mice on proteins encoded by four out of the five transcripts with cis-eQTLs in the 2.45 Mb locus (PCP4L1, ATP1A2, KCNJ9 (GIRK3), and IGSF9; **Fig.5B-E**) and three additional proteins distal to the locus (CADM3, AIM2, RGS7). We could not obtain reliable blots for fifth protein, nicastrin (NCSTN). There was a significant increase in immunostaining of ATP1A2 (within 2.45-Mb region) and AIM2 (distal to 2.45-Mb region) and a trending decrease in KCNJ9 of J/N versus J/J genotypes. Thus, we identified two compelling candidate genes within the fine-mapped locus (Atp1a2, Kncj9) that show genotype-dependent changes in expression at both the transcript and protein levels. Note that out of the two statistically significant proteins within the 2.45-Mb region (ATP1A2, KCNJ9/GIRK3), only Atp1a2 survived significance after the p-value threshold was adjusted for the seven protein comparisons (p<0.05/7 = 0.0071).

### Genetic variants between substrains within the causal QTL interval on distal chromosome 1

Fifteen noncoding homozygous variants distinguished J and N substrains within the distal Chr 1 interval (**Table S6**). None of these variants had predicted high impact. Eight variants were located within introns of *Uap1*, *Nos1ap*, *Dusp12*, *Cfap126*, *Nectin4*, and *Cd48*. An additional three variants were associated with pseudogenes or lncRNAs (*e.g*., *Gm37030*, *Gm10522*, *Gm37010*) and the remaining variants were located in intergenic regions with one variant annotated as an enhancer. Comparison of variation across C57BL/6 and C57BL/10 substrains (Mortazavi *et al*. 2022) revealed that five variants in the region have a pattern consistent with being J private alleles while one variant was consistent with an N private allele. All but one of the remaining variants show clear lineage specific patterns differentiating the J and N lineages.

## DISCUSSION

We employed a C57BL/6 RCC combined with fine-mapping, cis-eQTL mapping, and protein analysis to triangulate on two compelling candidate genes underlying OXY locomotion and withdrawal – *Atp1a2* and *Kcnj9*. We identified a second locus and candidate gene on chromosome 5 for opioid withdrawal, namely *Gabra2*. For the distal chromosome 1 locus, we implemented a novel fine-mapping approach that we previously proposed (Bryant *et al*. 2018) and efficiently narrowed the locus to 2.45 Mb. Typically, congenic lines are backcrossed for at least 10 generations to remove genetic variation outside of QTL intervals (Davis *et al*. 2005). Here, the genetic background of C57BL/6 substrains is nearly isogenic and thus, the remote possibility for epistasis is even less likely, especially after just a single generation of backcrossing, let alone the three to eight additional generations of backcrossing implemented from N5-N10 generations.

The overlapping QTL intervals for OXY locomotion and withdrawal suggest a common underlying genetic basis. This hypothesis is supported by the N6-1 recombinant line (161-183 Mb) capturing a phenotype for OXY locomotion and withdrawal (**Fig.3B,I**). Intriguingly, however, the OXY-responsive allelic effect of J/J for EPM behavior was opposite in N6-1 (decrease in Open Arm Entries; **Fig.3I**) versus F2 mice (increase in Open Arm Entries; **Fig.2H**). Nevertheless, in both cases, the J/J genotype was the OXY-responsive allele to changes in EPM behavior. Once explanation is that a different experimenter phenotyped the N6-1 versus F2 (Chesler *et al*. 2002). Also, previous studies with C57BL/6 have observed both a decrease (Ozdemir *et al*. 2023) and an increase in EPM behaviors (Hodgson *et al*. 2008; Buckman *et al*. 2009; Hofford *et al*. 2009) during opioid withdrawal.

The distal chromosome 1 locus for OXY locomotor traits and withdrawal was previously associated with phenotypic variation in crosses that include B6J as a parental strain. A region on distal chromosome 1—aptly named QTL-rich region 1 (*Qrr1*)—extends from approximately 170 Mb (rs8242852; GRCm38) to 175.4 Mb (rs4136041; GRCm38) and contains alleles (likely private to B6J) that contribute to variation in transcript levels and many behavioral, pharmacological, metabolic, physiological, and immunological phenotypes (Mozhui *et al*. 2008). It is probable that *Qrr1* contains multiple variants that contribute to different types of trait variation. For example, a B6J private mutation (C50T) in a member of the tRNA730-ArgTCT (*n-Tr20*) was found to reduce the total tRNAArgUCU pool in B6J brain and contribute to B6 substrain differences in seizure susceptibility (Ishimura *et al*. 2014). This tRNA mutation likely underlies seizure susceptibility QTLs (Ferraro *et al*. 2001, 2004) and contributes to variance in transcript levels associated with the *Qrr1* locus (Kapur *et al*. 2020) in crosses that include B6J as a progenitor strain. It is worth noting that the location of nTr20 (Chr1:173.39-173.39 Mb) is just distal to our fine-mapped 2.45-Mb interval and is therefore unlikely to contribute to OXY behavioral traits. A microdeletion in *Ifi202b* (Chr1:173.96-173.98 Mb) has been proposed to regulate activity, exploration, and adiposity traits (Vogel *et al*. 2013). However, B6 substrains are not known to be polymorphic for the *Ifi202b* mutation (Vogel *et al*. 2013) and similar to the tRNA mutation, this mutation lies distal to our 2.45-Mb interval. More recently, *Aim2* (Chr1:173.35-173.46 Mb), which is also outside our critical interval, has been proposed as another candidate for modulating physiological/metabolic traits mapping to *Qrr1*, especially hypercholesterolemia and diet-induced obesity, based on fine-mapping (Kim *et al*. 2024) and phenotyping in knock-out mice (Gong *et al*. 2019).

Based on our approach combining fine-mapping and eQTL analysis, we propose that OXY trait variation may be mediated by genetic variation in *Atp1a2* and *Kcnj9*. We and others have found eQTLs and transcript and/or protein level differences in several *Qrr1* genes, including *Atp1a2* and *Kcnj9* (Hitzemann *et al*. 2003; Mozhui *et al*. 2008; Thifault *et al*. 2008; Boughter *et al*. 2012). Unfortunately, the latest available DNA sequencing data (Mortazavi *et al*. 2022) within the region that we have fine-mapped contains an unusual number of heterozygous calls that we suspect may be due to segmental one or more segmental duplications. For simplicity’s sake, we have only reported the homozygous variant calls, none of which are located within or near *Kcnj9* or *Atp1a2*. In fact, none of the protein-coding genes associated with the 15 homozygous variants within the 2.45-Mb fine-mapped region (*Uap1*, *Nos1ap*, *Dusp12*, *Cfap126*, *Nectin4*, *Cc48*, *Cd48*, *Cd84* - **Table S6**) matched with the eQTL transcripts we identified (**Fig.5A**). Future long-read sequencing studies in C57BL/6 substrains should help to clarify the suspected structural variation within this region and provide insight into the genomic source of the eQTLs we identified within this region.

One major candidate gene in which knockout affects opioid behavioral sensitivity and withdrawal is *Kcnj9*, which codes for <u>G</u> protein-inwardly <u>R</u>ectifying <u>K</u>+ channel 3 (GIRK3, a.k.a. Kir3.3). Multiple addictive substances acting directly (e.g., morphine) or indirectly through GPCRs to modulate GIRK3, thus unleashing the dopaminergic reward pathway and stimulating acute and reinforcing behaviors (Rifkin *et al*. 2017). Evidence indicates that morphine activates mu opioid receptors in VTA GABAergic neurons, leading to GIRK3-mediated neuronal inhibition, disinhibition of mesolimbic dopamine neurons, and locomotor activation (Kotecki *et al*. 2015). *Kcnj9*/GIRK3 (172.32 Mb) lies within the 170.16-172.61 Mb region, has a cis-eQTL that peaked at the behavioral QTL, and shows a decrease in KCNJ9/GIRK3 protein in J/N (**Figs.4,5**). Reduced KCNJ9 protein in J/N is predicted to reduce OXY locomotion and withdrawal, given that GIRK3 is a major effector of mu opioid receptor signaling and is necessary for opioid-induced locomotion (Kotecki *et al*. 2015), opioid antinociception and multiple opioid withdrawal phenotypes (Cruz *et al*. 2008; Kozell *et al*. 2009). Another F2 study between C57BL/6J and 129P3/J mapped a similar distal chromosome 1 locus for antinociception induced by multiple Gi/Go-coupled GPCR drug classes (opioids, alpha-2 adrenergics, cannabinoids); however, when testing *Kcnj9* knockouts, they found no difference in morphine antinociception (Smith *et al*. 2008). A portion of our 2.45-Mb region (172,138,540-172,571,795 bp; mm10) was also identified for withdrawal induced by sedative/hypnotics acting at the GABA-A receptor and *Kcnj9* knockouts showed reduced withdrawal (handling-induced convulsions) (Kozell *et al*. 2009). To summarize, *Kcnj9* is a compelling positional and functional candidate gene underlying OXY behavioral sensitivity and withdrawal.

A second positional/functional candidate within the 2.45-Mb region is *Atp1a2* (sodium/potassium-transporting ATPase subunit alpha-2) which has a cis-eQTL that we validated as showing a robust increase in protein in J/N. Atp1a2 codes for a plasma membrane protein enzyme that is primarily expressed in astrocytes and regulates Na+ and K+ concentrations across the cell membrane, thus potentially influencing resting potential, depolarization, neurotransmitter release, and excitability local neurons. An adaptation in brain dopamine- and norepinephrine Na+/K+ ATPase activity following chronic opioid administration is hypothesized to contribute to changes in neuronal firing underlying opioid dependence (Desaiah & Ho 1979). For example, short-term morphine treatment in vivo can stimulate striatal Na+/K+ ATPase activity and inhibit neuronal depolarization via a D2 dopamine receptors and long-term morphine treatment can decrease synaptic protein expression and activity of Na+/K+ ATPase (Prokai *et al*. 2005) and increase neuronal depolarization through a D1 dopamine receptor-dependent mechanism both through cAMP/PKA-dependent mechanisms and which ultimately depend on mu opioid receptor activation (Wu *et al*. 2006, 2007). Thus, repeated high-dose OXY could induce neuronal hyperexcitability and depolarization accompanying opioid dependence (Kong *et al*. 2001) through an Na+/K+ ATPase-dependent mechanism (Kong *et al*. 1997), a physiological effect whose magnitude could depend on genotype-dependent differences in ATP1A2 protein levels. Also, the causal genes underlying the distal chromosome 1 QTL could be different for acute (e.g., *Kcnj9*) versus chronic (e.g., *Atp1a2*) behavioral effects of OXY or that both genes could contribute to both OXY behaviors.

We identified a second medial chromosome 5 QTL for OXY withdrawal that overlapped with the *Gabra2*-containing QTL (Mulligan *et al*. 2019) we mapped and validated for methamphetamine stimulant sensitivity (Goldberg *et al*. 2021a). *Gabra2* is a compelling candidate gene for anxiety-like behavior during opioid withdrawal, given its importance in the anxiogenic and anxiolytic properties benzodiazepine-site drugs, alcohol dependence, and polydrug misuse (Löw *et al*. 2000; Engin *et al*. 2012). GABRA2-linked variants are associated with a disrupted connectome of reward circuits and cognitive deficits in heroin users (Sun *et al*. 2018), polysubstance use, and alcohol dependence (Nudmamud-Thanoi *et al*. 2020). The Gabra2 eQTL is mediated by a single intronic nucleotide deletion near a splice site in B6J that causes a loss-of-function (expression) at the mRNA and protein levels (Mulligan *et al*. 2019). Thus, *Gabra2* is a high priority candidate gene for opioid withdrawal. Nevertheless, the peak QTL marker (rs33209545; 59.83 Mb) was more proximally located versus *Gabra2* (70.93 Mb) and thus we also considered peak cis-eQTLs at 59.83 Mb (**Table S5**). Candidate genes include *Cpeb2*, *N4bp2*, *Gabra4*, *Pdgfra*, and *Clock*. A human linkage scan of comorbid dependence on multiple substances, including opioids, identified significant linkage with a locus containing *GABRA4*, *GABRB1*, and *CLOCK* (Yang *et al*. 2012). Furthermore, clock genes have long been implicated in model traits for OUD (Forde & Kalsi 2017), including opioid withdrawal (Wang *et al*. 2006; Li *et al*. 2009; Perreau-Lenz *et al*. 2010; Hood *et al*. 2011; Roy *et al*. 2018; Rahmati-Dehkordi *et al*. 2021).

While there are many strengths to this study (fine mapping, functional analysis at RNA and protein level), there are some limitations. First, despite our most valiant effort, we were unable to resolve the 2.45-Mb locus any further, due to reasons described above. Second, we only conducted cis-eQTL analysis from one tissue (striatum) and thus, additional cis-eQTLs could exist in other brain regions. We chose striatum for practical reasons as it is large and readily amenable to RNA-seq analysis and it is rich in mu opioid receptors that contribute to opioid-induced locomotor sensitivity (Severino *et al*. 2020). Third, while we obtained functional evidence for causal sources of behavior, we do not have any obvious candidate causal quantitative trait variants. At the moment, future validation studies of candidate genes will require modulation of gene expression (e.g., virally-mediated) rather than germline gene editing of candidate variants.

In summary, we extended C57BL/6 substrain differences to include opioid behavioral sensitivity and withdrawal and identified strong candidate genes on distal chromosome 1 (*Kcnj9*, *Atp1a2*) through a novel positional cloning strategy unique to reduced complexity crosses combined with multi-level functional analyses. We also identified a chromosome 5 for opioid withdrawal, where *Gabra2* is a compelling candidate gene. Given that C57BL/6 is the most widely used mouse strain in biomedical research and given the huge focus of preclinical addiction research on opioids (which largely employs C57BL/6 mice), investigators should be aware of these genetic sources of variance in opioid behaviors as they ponder which C57BL/6 substrain to use in their opioid studies.

## ACKNOWLEDGMENTS

Pieter Faber, Technical Director for the University of Chicago Genomics Facility.

## AUTHOR CONTRIBUTIONS

L.R.G.: Data collection, analysis, and writing of manuscript; B.M.B: Data analysis and writing of manuscript Y.A.: Data analysis; J.A.B.: Data analysis and writing of the manuscript; J.C. K.: Data collection and analysis; E.J.Y.: Data collection and analysis; S.L.K.: Data collection and analysis; E.R.R.: Data collection and analysis; D.F.J.: Data analysis; J.C.: Data analysis; A.M.L.: Data collection and analysis; K.P.L.: Data collection; J.A.S.: Data collection; T.A.D.: Data collection; S.B.C.: Data collection; N.Y.: Data analysis; M.T.F.: Data analysis; W.E.J.: Data analysis; M.K.M.: Data collection, analysis, and writing of the manuscript. C.D.B.: Data collection, analysis, and writing of the manuscript.

## DATA AVAILABILITY STATEMENT

Data are available and will be provided by the corresponding author upon request.

## REFERENCES

Adinoff, B. (2004) Neurobiologic processes in drug reward and addiction. Harv Rev Psychiatry 12, 305–320.

Akinola, L.S., Mckiver, B., Toma, W., Zhu, A.Z.X., Tyndale, R.F., Kumar, V. & Damaj, M.I. (2019) C57BL/6 Substrain Differences in Pharmacological Effects after Acute and Repeated Nicotine Administration. Brain Sci 9.

Babbs, R.K., Beierle, J.A., Yao, E.J., Kelliher, J.C., Medeiros, A.R., Anandakumar, J., Shah, A.A., Chen, M.M., Johnson, W.E. & Bryant, C.D. (2020) The effect of the demyelinating agent cuprizone on binge-like eating of sweetened palatable food in female and male C57BL/6 substrains. Appetite 150, 104678.

Beierle, J.A., Yao, E.J., Goldstein, S.I., Lynch, W.B., Scotellaro, J.L., Shah, A.A., Sena, K.D., Wong, A.L., Linnertz, C.L., Averin, O., Moody, D.E., Reilly, C.A., Peltz, G., Emili, A., Ferris, M.T. & Bryant, C.D. (2022a) Zhx2 Is a Candidate Gene Underlying Oxymorphone Metabolite Brain Concentration Associated with State-Dependent Oxycodone Reward. J Pharmacol Exp Ther 382, 167–180.

Beierle, J.A., Yao, E.J., Goldstein, S.I., Scotellaro, J.L., Sena, K.D., Linnertz, C.A., Willits, A.B., Kader, L., Young, E.E., Peltz, G., Emili, A., Ferris, M.T. & Bryant, C.D. (2022b) Genetic basis of thermal nociceptive sensitivity and brain weight in a BALB/c reduced complexity cross. Mol Pain 18, 17448069221079540.

Boughter, J.D., Jr, Mulligan, M.K., St John, S.J., Tokita, K., Lu, L., Heck, D.H. & Williams, R.W. (2012) Genetic control of a central pattern generator: rhythmic oromotor movement in mice is controlled by a major locus near Atp1a2. PLoS One 7, e38169.

Broman, K.W. & Sen, S. (2009) A Guide to QTL Mapping with R/qtl., Statistics for Biology and Health. 1st edn. Springer-Verlag, Inc., New York.

Broman, K.W., Wu, H., Sen, S. & Churchill, G.A. (2003) R/qtl: QTL mapping in experimental crosses. Bioinformatics (Oxford, England) 19, 889–90.

Bryant, C.D., Bagdas, D., Goldberg, L.R., Khalefa, T., Reed, E.R., Kirkpatrick, S.L., Kelliiher, J.C., Chen, M.M., Johnson, W.E., Mulligan, M.K. & Damaj, M.I. (2019) C57BL/6 substrain differences in inflammatory and neuropathic nociception and genetic mapping of a major quantitative trait locus underlying acute thermal nociception. Mol Pain 1744806918825046.

Bryant, C.D., Eitan, S., Sinchak, K., Fanselow, M.S. & Evans, C.J. (2006a) NMDA receptor antagonism disrupts the development of morphine analgesic tolerance in male, but not female C57BL/6J mice. AmJPhysiolRegulIntegrCompPhysiol 291, R315–26.

Bryant, C.D., Ferris, M.T., De Villena, F.P.M., Damaj, M.I., Kumar, V. & Mulligan, M.K. (2018) Reduced complexity cross design for behavioral genetics. In Gerlai, R.T. (ed), Molecular-Genetic and Statistical Techniques for Behavioral and Neural Research, pp. 165–190.

Bryant, C.D., Healy, A.F., Ruan, Q.T., Coehlo, M.A., Lustig, E., Yazdani, N., Luttik, K.P., Tran, T., Swancy, I., Brewin, L.W., Chen, M.M. & Szumlinski, K.K. (2021) Sex-dependent effects of an Hnrnph1 mutation on fentanyl addiction-relevant behaviors but not antinociception in mice. Genes Brain Behav 20, e12711.

Bryant, C.D., Parker, C.C., Zhou, L., Olker, C., Chandrasekaran, R.Y., Wager, T.T., Bolivar, V.J., Loudon, A.S., Vitaterna, M.H., Turek, F.W. & Palmer, A.A. (2012) Csnk1e is a genetic regulator of sensitivity to psychostimulants and opioids. Neuropsychopharmacology 37, 1026–1035.

Bryant, C.D., Roberts, K.W., Byun, J.S., Fanselow, M.S. & Evans, C.J. (2006b) Morphine analgesic tolerance in 129P3/J and 129S6/SvEv mice. Pharmacol Biochem Behav 85, 769–779.

Bryant, C.D., Smith, D.J., Kantak, K.M., Nowak, T.S., Williams, R.W., Damaj, M.I., Redei, E.E., Chen, H. & Mulligan, M.K. (2020) Facilitating Complex Trait Analysis via Reduced Complexity Crosses. Trends Genet 36, 549–562.

Bryant, C.D., Zhang, N.N., Sokoloff, G., Fanselow, M.S., Ennes, H.S., Palmer, A.A. & McRoberts, J.A. (2008) Behavioral differences among C57BL/6 substrains: implications for transgenic and knockout studies. JNeurogenet 22, 315– 331.

Buckman, S.G., Hodgson, S.R., Hofford, R.S. & Eitan, S. (2009) Increased elevated plus maze open-arm time in mice during spontaneous morphine withdrawal. BehavBrain Res 197, 454–456.

Chesler, E.J., Wilson, S.G., Lariviere, W.R., Rodriguez-Zas, S.L. & Mogil, J.S. (2002) Identification and ranking of genetic and laboratory environment factors influencing a behavioral trait, thermal nociception, via computational analysis of a large data archive. Neurosci Biobehav Rev 26, 907–923.

Cicero, T.J. & Ellis, M.S. (2015) Abuse-Deterrent Formulations and the Prescription Opioid Abuse Epidemic in the United States: Lessons Learned From OxyContin. JAMA Psychiatry 72, 424–430.

Contreras, K.M., Buzzi, B., Vaughn, J., Caillaud, M., Altarifi, A.A., Olszewski, E., Walentiny, D.M., Beardsley, P.M. & Damaj, M.I. (2024) Characterization and validation of a spontaneous acute and protracted oxycodone withdrawal model in male and female mice. Pharmacol Biochem Behav 173795.

Cruz, H.G., Berton, F., Sollini, M., Blanchet, C., Pravetoni, M., Wickman, K. & Lüscher, C. (2008) Absence and rescue of morphine withdrawal in GIRK/Kir3 knock-out mice. J Neurosci 28, 4069–4077.

Davis, R.C., Schadt, E.E., Smith, D.J., Hsieh, E.W.Y., Cervino, A.C.L., van Nas, A., Rosales, M., Doss, S., Meng, H., Allayee, H. & Lusis, A.J. (2005) A genome-wide set of congenic mouse strains derived from DBA/2J on a C57BL/6J background. Genomics 86, 259–270.

Deak, J.D., Zhou, H., Galimberti, M., Levey, D.F., Wendt, F.R., Sanchez-Roige, S., Hatoum, A.S., Johnson, E.C., Nunez, Y.Z., Demontis, D., Børglum, A.D., Rajagopal, V.M., Jennings, M.V., Kember, R.L., Justice, A.C., Edenberg, H.J., Agrawal, A., Polimanti, R., Kranzler, H.R. & Gelernter, J. (2022) Genome-wide association study in individuals of European and African ancestry and multi-trait analysis of opioid use disorder identifies 19 independent genome-wide significant risk loci. Mol Psychiatry 27, 3970–3979.

Desaiah, D. & Ho, I.K. (1979) Effects of acute and continuous morphine administration on catecholamine-sensitive adenosine triphosphatase in mouse brain. J Pharmacol Exp Ther 208, 80–85.

Di Chiara, G. & Imperato, A. (1988) Drugs abused by humans preferentially increase synaptic dopamine concentrations in the mesolimbic system of freely moving rats. Proceedings of the National Academy of Sciences of the United States of America 85, 5274–8.

Ducci, F. & Goldman, D. (2012) The genetic basis of addictive disorders. PsychiatrClinNorth Am 35, 495–519.

Eitan, S., Bryant, C.D., Saliminejad, N., Yang, Y.C., Vojdani, E., Keith, D., Jr, Polakiewicz, R. & Evans, C.J. (2003) Brain region-specific mechanisms for acute morphine-induced mitogen-activated protein kinase modulation and distinct patterns of activation during analgesic tolerance and locomotor sensitization. JNeurosci 23, 8360–8369.

Engin, E., Liu, J. & Rudolph, U. (2012) α2-containing GABA(A) receptors: a target for the development of novel treatment strategies for CNS disorders. Pharmacol Ther 136, 142–152.

Ferraj, A., Audano, P.A., Balachandran, P., Czechanski, A., Flores, J.I., Radecki, A.A., Mosur, V., Gordon, D.S., Walawalkar, I.A., Eichler, E.E., Reinholdt, L.G. & Beck, C.R. (2023) Resolution of structural variation in diverse mouse genomes reveals chromatin remodeling due to transposable elements. Cell Genom 3, 100291.

Ferraro, T.N., Golden, G.T., Smith, G.G., Longman, R.L., Snyder, R.L., DeMuth, D., Szpilzak, I., Mulholland, N., Eng, E., Lohoff, F.W., Buono, R.J. & Berrettini, W.H. (2001) Quantitative genetic study of maximal electroshock seizure threshold in mice: evidence for a major seizure susceptibility locus on distal chromosome 1. Genomics 75, 35– 42.

Ferraro, T.N., Golden, G.T., Smith, G.G., Martin, J.F., Lohoff, F.W., Gieringer, T.A., Zamboni, D., Schwebel, C.L., Press, D.M., Kratzer, S.O., Zhao, H., Berrettini, W.H. & Buono, R.J. (2004) Fine mapping of a seizure susceptibility locus on mouse Chromosome 1: nomination of Kcnj10 as a causative gene. Mamm Genome 15, 239–251.

Forde, L.A. & Kalsi, G. (2017) Addiction and the Role of Circadian Genes. J Stud Alcohol Drugs 78, 645–653.

Gaddis, N., Mathur, R., Marks, J., Zhou, L., Quach, B., Waldrop, A., Levran, O., Agrawal, A., Randesi, M., Adelson, M., Jeffries, P.W., Martin, N.G., Degenhardt, L., Montgomery, G.W., Wetherill, L., Lai, D., Bucholz, K., Foroud, T., Porjesz, B., Runarsdottir, V., Tyrfingsson, T., Einarsson, G., Gudbjartsson, D.F., Webb, B.T., Crist, R.C., Kranzler, H.R., Sherva, R., Zhou, H., Hulse, G., Wildenauer, D., Kelty, E., Attia, J., Holliday, E.G., McEvoy, M., Scott, R.J., Schwab, S.G., Maher, B.S., Gruza, R., Kreek, M.J., Nelson, E.C., Thorgeirsson, T., Stefansson, K., Berrettini, W.H., Gelernter, J., Edenberg, H.J., Bierut, L., Hancock, D.B. & Johnson, E.O. (2022) Multi-trait genome-wide association study of opioid addiction: OPRM1 and beyond. Sci Rep 12, 16873.

Gelernter, J. & Polimanti, R. (2021) Genetics of substance use disorders in the era of big data. Nat Rev Genet 22, 712– 729.

Goldberg, L.R., Kirkpatrick, S.L., Yazdani, N., Luttik, K.P., Lacki, O.A., Babbs, R.K., Jenkins, D.F., Johnson, W.E. & Bryant, C.D. (2017) Casein kinase 1-epsilon deletion increases mu opioid receptor-dependent behaviors and binge eating1. Genes Brain Behav 16, 725–738.

Goldberg, L.R., Yao, E.J., Kelliher, J.C., Reed, E.R., Wu Cox, J., Parks, C., Kirkpatrick, S.L., Beierle, J.A., Chen, M.M., Johnson, W.E., Homanics, G.E., Williams, R.W., Bryant, C.D. & Mulligan, M.K. (2021a) A quantitative trait variant in Gabra2 underlies increased methamphetamine stimulant sensitivity. Genes Brain Behav 20, e12774.

Goldberg, L.R., Yao, E.J., Kelliher, J.C., Reed, E.R., Wu Cox, J., Parks, C., Kirkpatrick, S.L., Beierle, J.A., Chen, M.M., Johnson, W.E., Homanics, G.E., Williams, R.W., Bryant, C.D. & Mulligan, M.K. (2021b) A quantitative trait variant in Gabra2 underlies increased methamphetamine stimulant sensitivity. BioRxiv 450337.

Gong, Z., Zhang, X., Su, K., Jiang, R., Sun, Z., Chen, W., Forno, E., Goetzman, E.S., Wang, J., Dong, H.H., Dutta, P. & Muzumdar, R. (2019) Deficiency in AIM2 induces inflammation and adipogenesis in white adipose tissue leading to obesity and insulin resistance. Diabetologia 62, 2325–2339.

Harkness, J.H., Shi, X., Janowsky, A. & Phillips, T.J. (2015) Trace Amine-Associated Receptor 1 Regulation of Methamphetamine Intake and Related Traits. Neuropsychopharmacology 40, 2175–2184.

Hawkins, N.A., Nomura, T., Duarte, S., Barse, L., Williams, R.W., Homanics, G.E., Mulligan, M.K., Contractor, A. & Kearney, J.A. (2021) Gabra2 is a genetic modifier of Dravet syndrome in mice. Mamm Genome 32, 350–363.

Hegmann, J.P. & Possidente, B. (1981) Estimating genetic correlations from inbred strains. BehavGenet 11, 103–114.

Hitzemann, R., Malmanger, B., Reed, C., Lawler, M., Hitzemann, B., Coulombe, S., Buck, K., Rademacher, B., Walter, N., Polyakov, Y., Sikela, J., Gensler, B., Burgers, S., Williams, R.W., Manly, K., Flint, J. & Talbot, C. (2003) A strategy for the integration of QTL, gene expression, and sequence analyses. Mamm Genome 14, 733–47.

Ho, M.K., Goldman, D., Heinz, A., Kaprio, J., Kreek, M.J., Li, M.D., Munafo, M.R. & Tyndale, R.F. (2010) Breaking barriers in the genomics and pharmacogenetics of drug addiction. ClinPharmacolTher 88, 779–791.

Hodgson, S.R., Hofford, R.S., Norris, C.J. & Eitan, S. (2008) Increased elevated plus maze open-arm time in mice during naloxone-precipitated morphine withdrawal. BehavPharmacol 19, 805–811.

Hofford, R.S., Hodgson, S.R., Roberts, K.W., Bryant, C.D., Evans, C.J. & Eitan, S. (2009) Extracellular signal-regulated kinase activation in the amygdala mediates elevated plus maze behavior during opioid withdrawal. BehavPharmacol 20, 576–583.

Hood, S., Cassidy, P., Mathewson, S., Stewart, J. & Amir, S. (2011) Daily morphine injection and withdrawal disrupt 24-h wheel running and PERIOD2 expression patterns in the rat limbic forebrain. Neuroscience 186, 65–75.

Ishimura, R., Nagy, G., Dotu, I., Zhou, H., Yang, X.-L., Schimmel, P., Senju, S., Nishimura, Y., Chuang, J.H. & Ackerman, S.L. (2014) RNA function. Ribosome stalling induced by mutation of a CNS-specific tRNA causes neurodegeneration. Science 345, 455–459.

Jimenez Chavez, C.L., Bryant, C.D., Munn-Chernoff, M.A. & Szumlinski, K.K. (2021) Selective Inhibition of PDE4B Reduces Binge Drinking in Two C57BL/6 Substrains. Int J Mol Sci 22.

Kapur, M., Ganguly, A., Nagy, G., Adamson, S.I., Chuang, J.H., Frankel, W.N. & Ackerman, S.L. (2020) Expression of the Neuronal tRNA n-Tr20 Regulates Synaptic Transmission and Seizure Susceptibility. Neuron 108, 193–208.e9.

Keane, T.M., Goodstadt, L., Danecek, P., White, M.A., Wong, K., Yalcin, B., Heger, A., Agam, A., Slater, G., Goodson, M., Furlotte, N.A., Eskin, E., Nellaker, C., Whitley, H., Cleak, J., Janowitz, D., Hernandez-Pliego, P., Edwards, A., Belgard, T.G., Oliver, P.L., McIntyre, R.E., Bhomra, A., Nicod, J., Gan, X., Yuan, W., van der Weyden, L., Steward, C.A., Bala, S., Stalker, J., Mott, R., Durbin, R., Jackson, I.J., Czechanski, A., Guerra-Assuncao, J.A., Donahue, L.R., Reinholdt, L.G., Payseur, B.A., Ponting, C.P., Birney, E., Flint, J. & Adams, D.J. (2011) Mouse genomic variation and its effect on phenotypes and gene regulation. Nature 477, 289–294.

Kibaly, C., Alderete, J.A., Liu, S.H., Nasef, H.S., Law, P.-Y., Evans, C.J. & Cahill, C.M. (2021) Oxycodone in the Opioid Epidemic: High “Liking”, “Wanting”, and Abuse Liability. Cell Mol Neurobiol 41, 899–926.

Kim, J.H., Simpkins, M.A., Williams, N.T., Cimino, E., Simon, J., Richmond, T.R., Youther, J., Slutz, H. & Denvir, J. (2024) Tachol1 QTL on mouse chromosome 1 is responsible for hypercholesterolemia and diet-induced obesity. Mamm Genome.

Kirkpatrick, S.L. & Bryant, C.D. (2015) Behavioral architecture of opioid reward and aversion in C57BL/6 substrains. FrontBehavNeurosci 8, 450.

Kirkpatrick, S.L., Goldberg, L.R., Yazdani, N., Babbs, R.K., Wu, J., Reed, E.R., Jenkins, D.F., Bolgioni, A.F., Landaverde, K.I., Luttik, K.P., Mitchell, K.S., Kumar, V., Johnson, W.E., Mulligan, M.K., Cottone, P. & Bryant, C.D. (2017) Cytoplasmic FMR1-Interacting Protein 2 Is a Major Genetic Factor Underlying Binge Eating. BiolPsychiatry 81, 757–769.

Kong, J.Q., Leedham, J.A., Taylor, D.A. & Fleming, W.W. (1997) Evidence that tolerance and dependence of guinea pig myenteric neurons to opioids is a function of altered electrogenic sodium-potassium pumping. J Pharmacol Exp Ther 280, 593–599.

Kong, J.Q., Meng, J., Biser, P.S., Fleming, W.W. & Taylor, D.A. (2001) Cellular depolarization of neurons in the locus ceruleus region of the guinea pig associated with the development of tolerance to opioids. J Pharmacol Exp Ther 298, 909–916.

Kotecki, L., Hearing, M., McCall, N.M., Marron Fernandez de Velasco, E., Pravetoni, M., Arora, D., Victoria, N.C., Munoz, M.B., Xia, Z., Slesinger, P.A., Weaver, C.D. & Wickman, K. (2015) GIRK Channels Modulate Opioid-Induced Motor Activity in a Cell Type- and Subunit-Dependent Manner. J Neurosci 35, 7131–7142.

Kozell, L.B., Walter, N.A.R., Milner, L.C., Wickman, K. & Buck, K.J. (2009) Mapping a barbiturate withdrawal locus to a 0.44 Mb interval and analysis of a novel null mutant identify a role for Kcnj9 (GIRK3) in withdrawal from pentobarbital, zolpidem, and ethanol. J Neurosci 29, 11662–11673.

Kumar, V., Kim, K., Joseph, C., Kourrich, S., Yoo, S.H., Huang, H.C., Vitaterna, M.H., de Villena, F.P., Churchill, G., Bonci, A. & Takahashi, J.S. (2013) C57BL/6N mutation in Cytoplasmic FMRP interacting protein 2 regulates cocaine response. Science 342, 1508–1512.

Li, S., Liu, L., Jiang, W. & Lu, L. (2009) Morphine withdrawal produces circadian rhythm alterations of clock genes in mesolimbic brain areas and peripheral blood mononuclear cells in rats. J Neurochem 109, 1668–1679.

Löw, K., Crestani, F., Keist, R., Benke, D., Brünig, I., Benson, J.A., Fritschy, J.M., Rülicke, T., Bluethmann, H., Möhler, H. & Rudolph, U. (2000) Molecular and neuronal substrate for the selective attenuation of anxiety. Science 290, 131– 134.

Matsuo, N., Takao, K., Nakanishi, K., Yamasaki, N., Tanda, K. & Miyakawa, T. (2010) Behavioral profiles of three C57BL/6 substrains. FrontBehavNeurosci 4, 29.

Miner, N.B., Elmore, J.S., Baumann, M.H., Phillips, T.J. & Janowsky, A. (2017) Trace amine-associated receptor 1 regulation of methamphetamine-induced neurotoxicity. Neurotoxicology 63, 57–69.

Mortazavi, M., Ren, Y., Saini, S., Antaki, D., St Pierre, C.L., Williams, A., Sohni, A., Wilkinson, M.F., Gymrek, M., Sebat, J. & Palmer, A.A. (2022) SNPs, short tandem repeats, and structural variants are responsible for differential gene expression across C57BL/6 and C57BL/10 substrains. Cell Genom 2, 100102.

Mozhui, K., Ciobanu, D.C., Schikorski, T., Wang, X., Lu, L. & Williams, R.W. (2008) Dissection of a QTL hotspot on mouse distal chromosome 1 that modulates neurobehavioral phenotypes and gene expression. PLoS Genet 4, e1000260.

Mulligan, M.K., Abreo, T., Neuner, S.M., Parks, C., Watkins, C.E., Houseal, M.T., Shapaker, T.M., Hook, M., Tan, H., Wang, X., Ingels, J., Peng, J., Lu, L., Kaczorowski, C.C., Bryant, C.D., Homanics, G.E. & Williams, R.W. (2019) Identification of a functional non-coding variant in the GABAA receptor α2 subunit of the C57BL/6J mouse reference genome: Major implications for neuroscience research. Frontiers in Genetics 540211.

Mulligan, M.K., Ponomarev, I., Boehm, S.L., 2nd, Owen, J.A., Levin, P.S., Berman, A.E., Blednov, Y.A., Crabbe, J.C., Williams, R.W., Miles, M.F. & Bergeson, S.E. (2008) Alcohol trait and transcriptional genomic analysis of C57BL/6 substrains. Genes Brain Behav 7, 677–689.

Nassar, L.R., Barber, G.P., Benet-Pagès, A., Casper, J., Clawson, H., Diekhans, M., Fischer, C., Gonzalez, J.N., Hinrichs, A.S., Lee, B.T., Lee, C.M., Muthuraman, P., Nguy, B., Pereira, T., Nejad, P., Perez, G., Raney, B.J., Schmelter, D., Speir, M.L., Wick, B.D., Zweig, A.S., Haussler, D., Kuhn, R.M., Haeussler, M. & Kent, W.J. (2023) The UCSC Genome Browser database: 2023 update. Nucleic Acids Res 51, D1188–D1195.

Nudmamud-Thanoi, S., Veerasakul, S. & Thanoi, S. (2020) Pharmacogenetics of drug dependence: Polymorphisms of genes involved in GABA neurotransmission. Neurosci Lett 726, 134463.

Ozdemir, D., Allain, F., Kieffer, B.L. & Darcq, E. (2023) Advances in the characterization of negative affect caused by acute and protracted opioid withdrawal using animal models. Neuropharmacology 232, 109524.

Perdue, T., Carlson, R., Daniulaityte, R., Silverstein, S.M., Bluthenthal, R.N., Valdez, A. & Cepeda, A. (2024) Characterizing prescription opioid, heroin, and fentanyl initiation trajectories: A qualitative study. Soc Sci Med 340, 116441.

Perreau-Lenz, S., Sanchis-Segura, C., Leonardi-Essmann, F., Schneider, M. & Spanagel, R. (2010) Development of morphine-induced tolerance and withdrawal: involvement of the clock gene mPer2. Eur Neuropsychopharmacol 20, 509–517.

Peterson, R. & Cavanaugh, J. (2019) Ordered quantile normalization: a semiparametric transformation built for the cross-validation era. Journal of Applied Statistics 1–16.

Phillips, T.J., Roy, T., Aldrich, S.J., Baba, H., Erk, J., Mootz, J.R.K., Reed, C. & Chesler, E.J. (2021) Confirmation of a Causal Taar1 Allelic Variant in Addiction-Relevant Methamphetamine Behaviors. Front Psychiatry 12, 725839.

Prokai, L., Zharikova, A.D. & Stevens, S.M. (2005) Effect of chronic morphine exposure on the synaptic plasma-membrane subproteome of rats: a quantitative protein profiling study based on isotope-coded affinity tags and liquid chromatography/mass spectrometry. J Mass Spectrom 40, 169–175.

Rahmati-Dehkordi, F., Ghaemi-Jandabi, M., Garmabi, B., Semnanian, S. & Azizi, H. (2021) Circadian rhythm influences naloxone induced morphine withdrawal and neuronal activity of lateral paragigantocellularis nucleus. Behav Brain Res 414, 113450.

Reed, C., Baba, H., Zhu, Z., Erk, J., Mootz, J.R., Varra, N.M., Williams, R.W. & Phillips, T.J. (2017) A Spontaneous Mutation in Taar1 Impacts Methamphetamine-Related Traits Exclusively in DBA/2 Mice from a Single Vendor. Front Pharmacol 8, 993.

Rifkin, R.A., Moss, S.J. & Slesinger, P.A. (2017) G Protein-Gated Potassium Channels: A Link to Drug Addiction. Trends Pharmacol Sci 38, 378–392.

Ritchie, M.E., Phipson, B., Wu, D., Hu, Y., Law, C.W., Shi, W. & Smyth, G.K. (2015) limma powers differential expression analyses for RNA-sequencing and microarray studies. Nucleic Acids Res 43, e47.

Roy, K., Bhattacharyya, P. & Deb, I. (2018) Naloxone precipitated morphine withdrawal and clock genes expression in striatum: A comparative study in three different protocols for the development of morphine dependence. Neurosci Lett 685, 24–29.

Ruan, Q.T., Yazdani, N., Blum, B.C., Beierle, J.A., Lin, W., Coelho, M.A., Fultz, E.K., Healy, A.F., Shahin, J.R., Kandola, A.K., Luttik, K.P., Zheng, K., Smith, N.J., Cheung, J., Mortazavi, F., Apicco, D.J., Ragu Varman, D., Ramamoorthy, S., Ash, P.E.A., Rosene, D.L., Emili, A., Wolozin, B., Szumlinski, K.K. & Bryant, C.D. (2020) A Mutation in Hnrnph1 That Decreases Methamphetamine-Induced Reinforcement, Reward, and Dopamine Release and Increases Synaptosomal hnRNP H and Mitochondrial Proteins. J Neurosci 40, 107–130.

Schulteis, G., Yackey, M., Risbrough, V. & Koob, G.F. (1998) Anxiogenic-like effects of spontaneous and naloxone-precipitated opiate withdrawal in the elevated plus-maze. Pharmacol Biochem Behav 60, 727–731.

Severino, A.L., Mittal, N., Hakimian, J.K., Velarde, N., Minasyan, A., Albert, R., Torres, C., Romaneschi, N., Johnston, C., Tiwari, S., Lee, A.S., Taylor, A.M., Gavériaux-Ruff, C., Kieffer, B.L., Evans, C.J., Cahill, C.M. & Walwyn, W.M. (2020) μ-Opioid Receptors on Distinct Neuronal Populations Mediate Different Aspects of Opioid Reward-Related Behaviors. eNeuro 7, ENEURO.0146-20.2020.

Shi, X., Walter, N.A., Harkness, J.H., Neve, K.A., Williams, R.W., Lu, L., Belknap, J.K., Eshleman, A.J., Phillips, T.J. & Janowsky, A. (2016) Genetic Polymorphisms Affect Mouse and Human Trace Amine-Associated Receptor 1 Function. PLoS One 11, e0152581.

Simon, M.M., Greenaway, S., White, J.K., Fuchs, H., Gailus-Durner, V., Sorg, T., Wong, K., Bedu, E., Cartwright, E.J., Dacquin, R., Djebali, S., Estabel, J., Graw, J., Ingham, N.J., Jackson, I.J., Lengeling, A., Mandillo, S., Marvel, J., Meziane, H., Preitner, F., Puk, O., Roux, M., Adams, D.J., Atkins, S., Ayadi, A., Becker, L., Blake, A., Brooker, D., Cater, H., Champy, M.F., Combe, R., Danecek, P., di Fenza, A., Gates, H., Gerdin, A.K., Golini, E., Hancock, J.M., Hans, W., Holter, S.M., Hough, T., Jurdic, P., Keane, T.M., Morgan, H., Muller, W., Neff, F., Nicholson, G., Pasche, B., Roberson, L.A., Rozman, J., Sanderson, M., Santos, L., Selloum, M., Shannon, C., Southwell, A., Tocchini-Valentini, G.P., Vancollie, V.E., Wells, S., Westerberg, H., Wurst, W., Zi, M., Yalcin, B., Ramirez-Solis, R., Steel, K.P., Mallon, A.M., Hrab 283 de Angelis, M., Herault, Y. & Brown, S.D. (2013) A comparative phenotypic and genomic analysis of C57BL/6J and C57BL/6N mouse strains. Genome Biol 14, R82.

Smith, S.B., Marker, C.L., Perry, C., Liao, G., Sotocinal, S.G., Austin, J.-S., Melmed, K., Clark, J.D., Peltz, G., Wickman, K. & Mogil, J.S. (2008) Quantitative trait locus and computational mapping identifies Kcnj9 (GIRK3) as a candidate gene affecting analgesia from multiple drug classes. Pharmacogenet Genomics 18, 231–241.

Sun, Y., Zhang, Y., Zhang, D., Chang, S., Jing, R., Yue, W., Lu, L., Chen, D., Sun, Y., Fan, Y. & Shi, J. (2018) GABRA2 rs279858-linked variants are associated with disrupted structural connectome of reward circuits in heroin abusers. Transl Psychiatry 8, 1–10.

Thifault, S., Ondrej, S., Sun, Y., Fortin, A., Skamene, E., Lalonde, R., Tremblay, J. & Hamet, P. (2008) Genetic determinants of emotionality and stress response in AcB/BcA recombinant congenic mice and in silico evidence of convergence with cardiovascular candidate genes. HumMolGenet 17, 331–344.

Vogel, H., Montag, D., Kanzleiter, T., Jonas, W., Matzke, D., Scherneck, S., Chadt, A., Töle, J., Kluge, R., Joost, H.-G. & Schürmann, A. (2013) An interval of the obesity QTL Nob3.38 within a QTL hotspot on chromosome 1 modulates behavioral phenotypes. PLoS One 8, e53025.

W, M., L, G., Se, H., Hs, R., Gr, R., A, T., P, F. & F, C. (2016) The Ensembl Variant Effect Predictor. Genome biology 17.

Wang, X., Wang, Y., Xin, H., Liu, Y., Wang, Y., Zheng, H., Jiang, Z., Wan, C., Wang, Z. & Ding, J.M. (2006) Altered expression of circadian clock gene, mPer1, in mouse brain and kidney under morphine dependence and withdrawal. J Circadian Rhythms 4, 9.

Warden, A.S., DaCosta, A., Mason, S., Blednov, Y.A., Mayfield, R.D. & Harris, R.A. (2020) Inbred Substrain Differences Influence Neuroimmune Response and Drinking Behavior. Alcohol Clin Exp Res 44, 1760–1768.

Wise, R.A. & Bozarth, M.A. (1987) A psychomotor stimulant theory of addiction. Psychological review 94, 469–92.

Wu, Z.-Q., Chen, J., Chi, Z.-Q. & Liu, J.-G. (2007) Involvement of dopamine system in regulation of Na+,K+-ATPase in the striatum upon activation of opioid receptors by morphine. Mol Pharmacol 71, 519–530.

Wu, Z.-Q., Li, M., Chen, J., Chi, Z.-Q. & Liu, J.-G. (2006) Involvement of cAMP/cAMP-dependent protein kinase signaling pathway in regulation of Na+,K+-ATPase upon activation of opioid receptors by morphine. Mol Pharmacol 69, 866–876.

Yalcin, B., Wong, K., Agam, A., Goodson, M., Keane, T.M., Gan, X., Nellaker, C., Goodstadt, L., Nicod, J., Bhomra, A., Hernandez-Pliego, P., Whitley, H., Cleak, J., Dutton, R., Janowitz, D., Mott, R., Adams, D.J. & Flint, J. (2011) Sequence-based characterization of structural variation in the mouse genome. Nature 477, 326–329.

Yang, B.-Z., Han, S., Kranzler, H.R., Farrer, L.A., Elston, R.C. & Gelernter, J. (2012) Autosomal linkage scan for loci predisposing to comorbid dependence on multiple substances. Am J Med Genet B Neuropsychiatr Genet 159B, 361–369.

Yao, E.J., Babbs, R.K., Kelliher, J.C., Luttik, K.P., Borrelli, K.N., Damaj, M.I., Mulligan, M.K. & Bryant, C.D. (2021) Systems genetic analysis of binge-like eating in a C57BL/6J x DBA/2J-F2 cross. Genes Brain Behav e12751.

Yazdani, N., Parker, C.C., Shen, Y., Reed, E.R., Guido, M.A., Kole, L.A., Kirkpatrick, S.L., Lim, J.E., Sokoloff, G., Cheng, R., Johnson, W.E., Palmer, A.A. & Bryant, C.D. (2015) Hnrnph1 Is A Quantitative Trait Gene for Methamphetamine Sensitivity. PLoS Genet 11, e1005713.

Yu, W., Mulligan, M.K., Williams, R.W. & Meisler, M.H. (2022) Correction of the hypomorphic Gabra2 splice site variant in mouse strain C57BL/6J modifies the severity of Scn8a encephalopathy. HGG Adv 3, 100064.

