## Supplementary Material_R1 for "*Atp1a2* and *Kcnj9* are candidate genes underlying sensitivity to oxycodone-induced locomotor activation and withdrawal-induced anxiety-like behaviors in C57BL/6 substrains"

**TABLE OF CONTENTS**

**MATERIALS AND METHODS………………………………………………………………………………p.2-7**

**RESULTS………………………………………………………………………………………………………p.7-9**

**SUPPLEMENTARY TABLES……………………………………………………………………………….p.9-17**

**SUPPLEMENTARY FIGURES………………………………………………………………………………p.18-28**

**REFERENCES…………………………………………………………………………………………………p.29-30**

**MATERIALS AND METHODS (further details)**

**Parental C57BL/6 substrains and F2 mice**

Colony rooms were maintained on a 12:12 h light–dark cycle (lights on at 0630 h). Mice were housed same-sex (2-4/cage with standard laboratory chow (Teklad 18% Protein Diet, Envigo, Indianapolis, IN, USA) and water available *ad libitum* except while mice were in the testing chambers for 30 min. Mice were 50-100 days old on the first day of training.

C57BL/6J (**B6J**; #000664) and C57BL/NJ (**B6NJ**; #005304) mice were purchased from The Jackson Laboratory (Bar Harbor, ME USA) at 7 weeks old and were habituated in the vivarium one week prior to experimental testing next door. Behavioral testing was performed during the light phase (0800 h-1300 h). For QTL mapping, B6J females were crossed to B6NJ males to generate B6J x B6NJ-F_1_ mice and B6J x B6NJ F_1_ offspring were intercrossed to generate B6J x B6NJ F_2_ mice. All mice within a cage were assigned the same treatment. 425 F2 mice (213 SAL, 212 OXY) were phenotyped in the OXY locomotor/CPP protocol during Weeks 1 and 2 (**Fig.2A**). A subset (118 SAL, 118 OXY) went on to receive a high-dose, multi-week OXY regimen (see below) over Weeks 3 and 4 (**Fig.2A**) followed by assessment of spontaneous withdrawal behaviors in the elevated plus maze (**EPM**). The majority of these F2 mice (82 SAL, 78 OXY) underwent hot plate testing for tolerance during Week 3; however a subset of the 236 F2 mice (76 total: n=36 SAL, 40 OXY) received the exact same multi-week high-dose OXY regimen without exposure to the hot plate during Week 3 (**Fig.2A**).

**Locomotor activity traits and conditioned place preference (Weeks 1 and 2)**

The Plexiglas apparatus for assessing locomotor activity and conditioned place preference (**CPP**) is a rectangular open field (40 cm length x 20 cm width x 45 cm tall; Lafayette Instruments, Lafayette, IN, USA) surrounded by a sound-attenuating chamber (MedAssociates, St. Albans, VT, USA). For place conditioning, mice were trained during Week 1 (Monday-Friday: Days 1-5), allowed a two-day consolidation period (Saturday-Sunday: Days 6-7), and assessed drug-free and state-dependent CPP during the first two days of Week 2 (Monday-Tuesday: Days 8-9) (Kirkpatrick & Bryant 2015). The apparatus was partitioned into two equally sized compartments with an infrared-transmitting plastic black divider containing a mouse entryway (5 cm x 6.25 cm) that was flipped upside down during training to confine mice to one side. We used distinct, removable floor textures to differentiate each side (Plaskolite Inc., Columbus, OH, USA). The left side has a wavy, smooth texture and the right side has a rougher, pointy texture (Kirkpatrick & Bryant 2015). Behavioral data were recorded using a security camera system (Swann Communications, Melbourne, Australia) and the recordings were subjected to video tracking analysis (Anymaze, Stoelting, Wood Dale, IL, USA). The conditioned place preference (**CPP**) procedure has been described previously in detail (Kirkpatrick & Bryant 2015; Goldberg *et al.* 2021). On Day(**D**) 1 (D1), initial preference for the drug-paired side (right side) was assessed; mice received SAL (i.p.), were placed into the left side (saline-paired side), and were allowed free access to both sides of the chamber for 30 min. On training days (D2-D5), mice received either drug (i.p.; D2, D4) or SAL (i.p.; D3, D5) and were confined to either the drug-paired (right side) or saline-paired side (left side) for 30 min, respectively. The main outcome measures were Distance (m), Spins (#; 360 degree rotations of the mouse nose and body) and Rotations (#; sequential crosses and thus circles traversing four, equally sized quadrants within an enclosed side of the CPP chamber) (Kirkpatrick & Bryant 2015). Mice were left undisturbed in their home cages on D6 and D7. On D8, drug-free CPP for the drug-paired side was assessed over 30 min using identical procedures to D1 (i.e., i.p. SAL injection, placement into left side, open access for 30 min). On D9, state-dependent CPP for the drug-paired side was similarly assessed as on D8, with the exception that mice once again received their training treatment/dose [OXY (i.p.) or SAL (i.p.)]. Mice were then left undisturbed in their home cages for the rest of Week 2. We quantified various locomotor activity measures (e.g., distance traveled, rotations around the perimeter, body spins) during initial preference, training days, and during final CPP (both drug-free and state-dependent). The main outcome measure for OXY-CPP was the difference in time (s) spent on the drug-paired side between D1 and D8 (drug-free) or D9 (state-dependent), based on the three main CPP dependent variables which were D1 OXY Side Time (s), D8 OXY Side Time (s), and D9 OXY Side Time (s).

**Procedures for baseline nociception and oxycodone-induced antinociception (Week 3)**

For F2 mice, following 4 daily injections of OXY (20 mg/kg, i.p.; Monday-Thursday; 1600 h), 16 h later (Friday, 0800 h), mice were habituated to the testing room for 1 h. Mice were then placed in a Plexiglas cylinder (15 cm diameter; 33.0 cm tall) on the 52.5degC hot plate (IITC Life Science Inc., Woodland Hills, CA, USA) and the latency to lick the hind paw was recorded, with a 60 s cut-off latency. Thirty min post-assessment of baseline pain sensitivity, mice were injected with a challenge dose of OXY (5 mg/kg, i.p.) and assessed for post-injection latencies at 30 min. Percent maximum possible effect (%MPE) at 30 min post-OXY was quantified as the measurement of antinociception using the following formula: %MPE = (Post-injection latency – Baseline Latency) / (60 – Baseline Latency) *100. Mice were left undisturbed in their home cages for Days 6 and 7 of Week 3.

Because we did not observe significant antinociceptive tolerance to OXY in F2 mice following the challenge dose of OXY (5 mg/kg, i.p.) and antinociceptive assessment at 30 min (**Fig.S4B**), for N6-1 mice, we modified the procedure for tolerance assessment by including multiple time points for assessment of tolerance on test day, including earlier time points where OXY behavioral effects are expected to peak based on pilot data at the time in C57BL/6 substrains indicating peak OXY locomotor effects at 10 min post-OXY (i.p.). We also included a second assessment of baseline latency to ensure a reliable baseline measurement and report the average of the two values. Thirty min after the second baseline assessment, mice were injected with the same challenge dose of 5 mg/kg OXY (i.p.) as in F2 mice and were assessed for post-injection latencies at 10, 20, 30, and 40 min post-OXY. Percent MPE was calculated for each of the four time points (see formula above) and repeated measures ANOVA was conducted with Time as the repeated measure and Genotype, Treatment, and Sex as factors.

**Genotyping and QTL analysis in B6J x B6NJ-F2 mice**

Samples were genotyped using a custom Fluidigm array (San Francisco, CA USA) as previously described (Kirkpatrick *et al.* 2017; Bryant *et al.* 2019). The marker panel consisted of 96 SNP markers (**Table S1**) spaced at approximately 30 Mb. Markers with a failure rate of greater than 5% fail rate were excluded from mapping. Following QC, there were 425 mice (213 SAL, 213 OXY) and 79 markers used to map OXY induced locomotor traits, and 236 mice (118 SAL, 118 OXY) and 76 markers used to map EPM phenotypes. Genomic DNA samples were diluted to 100 ng/uL in low TE buffer (Teknova, Hollister, CA, USA). Single nucleotide polymorphisms (SNPs) were called using the Fluidigm SNP Genotyping Analysis Software and SNPtype Normalization.

QTL analysis was performed in R/qtl (Broman *et al.* 2003; Broman, & Sen 2009). The scanone function was used to calculate LOD scores. The marker position (cM) was estimated using the sex-averaged position using the Mouse Map Converter that was previously available via The Jackson Laboratory (<http://cgd.jax.org/mousemapconverter>) and an updated version is now available as an R package provided by Dr. Karl Broman (<https://cran.r-project.org/web/packages/mmconvert/mmconvert.pdf>; Github: https://github.com/rqtl/mmconvert). QTL analysis was performed in r/QTL using the “scanone” function and Haley-Knott (HK) regression in R/qtl (Broman *et al.* 2003). Permutation analysis (perm=1000) was used to compute genome-wide significance thresholds (p < 0.05). For each significant QTL, both the Bayes credible interval and 1.5-LOD intervals are reported.

**Genotyping and generation of recombinant lines**

To genotype recombinant lines within the distal chromosome 1 region (163-181 Mb) and screen for new recombination events, we used the Sanger dataset (REL-1505, <http://www.sanger.ac.uk/>) to retrieve polymorphic SNPs underlying distal chromosome 1 (163-181 Mb) that distinguished B6NJ from B6J (Keane *et al.* 2011; Yalcin *et al.* 2011). Recombination events were monitored using a panel of 15 markers spanning from 163.13 Mb to 181.32 Mb on distal chromosome 1 (average marker spacing of 1.3 Mb) (**Table S2, S3**). We first confirmed that each of the markers was polymorphic between B6J and BNJ substrains prior to genotyping the recombinant lines. Polymerase chain reaction (PCR) primers were designed for each SNP (at least 50 bp upstream and 20 bp downstream) using Primer3 (<http://bioinfo.ut.ee/primer3-0.4.0/primer3/>). We used 50 ng/µl DNA in the PCR reaction and ran it through a 1% agarose gel via electrophoresis. We observed a single band for each of the primer sets, excised it from the gel and extracted the PCR product using the QIAquick® gel extraction kit (Qiagen, Valencia, CA, USA). Samples were sent to Genewiz (Genewiz, South Plainfield, NJ, USA) for Sanger sequencing. Following successful sequencing and verification of polymorphic SNPs, custom fluorescent SNP assays were sometimes used to accelerate the pace of genotyping (Life Technologies, Carslbad, CA, USA).

**Statistical analysis of recombinant lines**

For N6-1, D2 and D4 (OXY, 1.25 mg/kg, i.p.) behaviors were analyzed using two-way ANOVA (Genotype, Treatment) and three-way ANOVA (Sex) to test for replication of OXY behavioral QTLs. Two-way ANOVAs were also run for D3 and D5 (SAL, i.p.). Both drug-free (D8) and state-dependent OXY-CPP (D9) were analyzed via two-way ANOVA (Genotype, Treatment) and three-way ANOVA (Sex), with change in time spent on the OXY-paired side between D1 and D8 or D9 [D8/D9-D1 (s)] as the dependent variable. Elevated plus maze behaviors [% Open Arm Time, Open Arm Entries (#), Open Arm Distance (m)] were analyzed using two-way ANOVA (Genotype, Treatment). Bonferroni or Tukey’s post-hoc comparisons were used to identify sources of interactions, as indicated. For the remaining recombinant lines, D1 (open access to CPP chambers following SAL i.p.) and D2 (closed access to OXY-paired side) were analyzed separately using unpaired t-tests and a Bonferroni-adjusted significance level of p < 0.00625 (0.05/8 recombinant lines). Finally, in addition to simple pair-wise t-test comparisons, because we found in the F2 analysis that the distal chromosome 1 QTL for OXY locomotor traits was driven primarily by the females, we also used two-way ANOVA (Genotype, Sex) of D1 (SAL) and D2 (OXY) locomotor traits in each of the recombinant lines and conducted follow-up sex-specific analyses in those lines showing a Genotype x Sex interaction for a given trait.

**RNA-seq**

Briefly, striatum punches were collected from 23 OXY-trained F2 mice at 24 h after behavioral testing on the EPM. Brains were rapidly removed, sectioned with a brain matrix to obtain a 3 mm thick section, and a 2.5 mm diameter-wide punch of the striatum was collected as described (Yazdani *et al.* 2015). Left and right striatum punches were pooled, placed in RNAlater (Life Technologies, Grand Island, NY, USA) for 48 h at 4°C prior to storage in a -80°C freezer. Total RNA was extracted as described (Yazdani *et al.* 2015) using the RNeasy kit (Qiagen, Valencia, CA, USA). RNA was shipped to the University of Chicago Genomics Core Facility for cDNA library preparation using the Illumina TruSeq (oligo-dT; 100 bp paired-end reads). Libraries were prepared according to Illumina's detailed instructions accompanying the TruSeq® Stranded mRNA LT Kit (Part# RS-122-2101). The purified DNA was captured on an Illumina flow cell for cluster generation and sample libraries were sequenced at 23 samples per lane over 5 lanes (technical replicates) according to the manufacturer’s protocols on the Illumina HiSeq 4000 machine, yielding an average of 69.4 million total reads per sample. FASTQ files were quality checked via FASTQC and possessed Phred quality scores > 30 (i.e., less than 0.1% sequencing error).

**Striatal expression QTL (eQTL) mapping and regression of gene expression onto behavior**

Expression QTL (**eQTL**) analysis can help identify candidate genes for complex traits by providing a functional intermediary link between DNA variant and behavior. However, like QTL analysis of behavioral traits, eQTL analysis in an F2 cross is limited in resolution by the low number of historical recombination events in the F1 gametes that are inherited in F2 offspring (typically 1-2 recombination events per inherited chromatid). Thus, cis-eQTLs are not assigned to a variant within or even near the transcript-encoding gene. Rather, causal variants influencing “nearby” gene expression are in linkage with the nearest marker which could be located tens of Mb away from the actual transcript. Because we were interested in cis-eQTLs that co-mapped to the same region as the behavioral QTLs, we focused on those transcripts located within the behavioral QTL intervals that also showed peak linkage with the same peak marker for the behavioral QTLs. We conducted eQTL analysis with the same marker panel used for behavioral QTL analysis which was limited to a total of 96 markers. Thus, each marker can serve as a guidepost linked to multiple causal variants that together influence expression of multiple transcripts and we hypothesize that at least one of these transcripts influences behavior. With the assumption that the causal gene has an eQTL, the eQTL should theoretically show the same peak association with the same marker as the behavioral QTL.

We aligned FASTQ files to the mm38 genome via TopHat (Trapnell *et al.* 2012) and used the Ensembl sequence and genome annotation. We used *featureCounts* for read alignment. We used the same marker panel as above for *cis*-eQTL analysis. Exons with low expression (less than 10 reads total across all 115 count files) were removed. A a *cis*-eQTL was liberally defined as any transcript with a genome-wide significant association between expression and a polymorphic marker that was within 70 Mb of a SNP, given the large linkage disequilibrium in a lowly recombinant F2 cross and given that this was the largest distance between any two SNP markers. Analysis was conducted using *limma* with default TMM normalization and VOOM transformation (Law *et al.* 2014; Ritchie *et al.* 2015). To account for the presence of replicates of each sample in the data, we used the duplicateCorrelation() function to estimate the within-sample correlation which we then included in the lmFit() function. An ANOVA test was conducted for gene expression, with Genotype as a fixed effect and Sex as a covariate. Gene-level tests were conducted using the Likelihood Ratio test. A false discovery rate of 5% was employed as the cut-off for statistical significance (Benjamini & Hochberg 1995).

In addition to eQTL analysis, we also examined Pearson’s correlation between gene expression (normalized read counts) and D2 OXY distance which was the behavior used for fine-mapping the distal chromosome 1 QTL. We also used the limma package and lmfit() to fit a linear mixed model regressing read counts onto D2 OXY distance which was the behavior used for fine-mapping.

**Antibodies for candidate gene/transcript/protein analysis**

Striatal tissue from the N6-1 recombinant line was homogenized in RIPA buffer (Thermo Scientific, Waltham, MA, USA) containing 1x HALT protease/phosphatase inhibitor cocktail (Thermo Scientific) with 3 s burst from an ultrasonic homogenizer. Samples were spun at 17200 RCF for 20 min at 4º C. Supernatants were collected, and protein concentrations were determined via BCA protein estimation (Thermo Scientific). 30 µg of sample protein and loading buffer (BioRad, Hercules, CA, USA) was loaded into 4-15% Criterion TGX gels (BioRad) and run at 120 V until fully migrated. Gels were transferred onto nitrocellulose membranes (GE Healthcare, Chicago, IL, USA) overnight at 25 V at 4º C. Blots were then blocked with 5% milk for 1 h. Blots were probed overnight for each protein for each candidate gene at 4º C, and then for 1 h in a 1:10,000 dilution of peroxidase conjugated donkey anti rabbit antibody (Jackson Immunoresearch; cat. # 711-035-152). Imaging was conducted using a ChemiDoc XRS+ imager (BioRad). Blots were then stripped (Thermo Scientific, 46430) at 55°C for 15 min, blocked, incubated in Beta-actin antibody (1:50,000; Sigma Aldrich, #A2228) in 5% milk for 1 h, then peroxidase conjugated donkey anti mouse antibody for 1 h (1:10,000; Jackson Immunoresearch, cat. # 715-035-151), and then imaged. Immunoblot bands were quantified using densitometry analysis in NIH ImageJ. Raw densitometry values for each lane were normalized to beta actin.

Information for each antibody is as follows: PCP4L1 (1:10,000; ProteinTech, rabbit, polyclonal antibody, cat. # 25933-1-AP); NDUFS2 (1:10,000; abcam, rabbit, monoclonal antibody, EPR16266, cat. # ab192022, lot # GR217388-11); ATP1A2 (1:3,000; ProteinTech, rabbit, polyclonal antibody, cat. # 16836-1-AP); KCNJ9 (1:5,000; “GIRK3”, Sigma, PB247, rabbit, affinity-isolated, 094K1728); IGSF9 (1:10,000; Novus, lot #: E113781, polyclonal antibody, rabbit affinity-purified, NBP1-93676); CADM3 (1:5,000; abcam, ab69604, rabbit polyclonal antibody, GR309281-2); AIM2 (1:5,000; Cell Signaling Technology, Lot 1, March 2017, 13095S, rabbit, mouse-specific); RGS7 (1:5,000; Sigma, SAB4502633, rabbit, affinity-isolated, Lot: 3118326).

**RESULTS**

**Distal chromosome 1 recombinant lines spanning 163 Mb to 181 Mb resulting from repeated backcrossing to B6J**

The markers, primer sequences, and positions that we used to scan for recombination events and define the recombinant lines are provided in **Table S2**. The pedigree of the recombinant lines, beginning with the F2 founder that was heterozygous throughout the 163-181 Mb region, is illustrated in **Fig.S9**. Because F2 individuals from a reduced complexity cross between C57BL/6 substrains are nearly isogenic, the mixed background of the recombinant lines was not a concern, considering the simple genetic architecture (typically one major QTL accounting for a large proportion of the variance) and that population structure cannot arise from a breeding design involving continual backcrossing.

We fine-mapped the distal chromosome 1 QTL for OXY locomotion to 2.45 Mb. Pedigree, genotypes and breakpoints for recombinations for each founder are provided in **Table S3**. A single, heterozygous F2 female (#6712) throughout the QTL (163-181 Mb) was backcrossed to B6J to generate N3 mice, from which four N4 offspring were backcrossed and propagated. Three of these families (N4-3, N4-19, and N4-20) again had a heterozygous breeder throughout the 163-181 Mb region (**Fig.S9**). The fourth family, N4-18, had a heterozygous founder that inherited a proximal recombination between 170.2-172.6 Mb and the remaining distal recombinant segment was heterozygous through the 181 Mb marker.

This N4-18 female founder (#6874) was backcrossed to B6J to generate a female N5 founder (N5-10; #7646) that inherited a recombination from 173.1-173.7 Mb and was heterozygous through the 181 Mb marker. The N5-10 founder was used to propagate the N5-10 family and several N6-10 families that were all direct descendants and contained the same heterozygous segment from 173-181 Mb (N6-13,-14,-15,-16,-17,-18,-19; **Fig.S9**).

The N5-11 male founder (7715) descended from N4-19 and inherited a proximal recombination yielding a distal segment spanning 173/174-181 Mb. This founder propagated the N5-11 family as well as direct, N6 descendant lines containing the same segment (N6-4,-6,-11,-12). Finally, the N5-8 male founder (#7604) descended from N4-20 and inherited a proximal recombination that resulted in an even shorter distal segment spanning 175/178 Mb-181 Mb (**Fig.S9**).

N4-3 descendants were also heterozygous throughout the 163-181 Mb region (N5-2). However, the N6-1 female founder (#7583) inherited two new recombination events at the proximal and distal ends, yielding a heterozygous segment spanning approximately 163/164-180/181 Mb. This individual propagated the N6-1 family and two N7-1 families that were direct descendants of N6-1 and contained the same recombinant segment (N7-2,-3,-4; **Fig.S9**). Two other N7 families from N6-1 propagated additional families containing newly truncated recombinant segments, including a more proximally localized segment comprising N7-15 (163/164-170/173 Mb) and a more distally localized segment arising from a recombinant offspring of N7-2 (170/173-180 Mb) that comprised N8-7 and its direct descendants (N9-2, -3, and -4; **Fig.S9**).

**Fine mapping the distal chromosome 1 behavioral QTL for D2 OXY locomotor traits in recombinant lines**

In examining D1 Distance following SAL (i.p.), none of the other lines besides N6-1 showed a significant difference (p_adjusted_<0.05/8 recombinant lines=0.00625): N9-8 (t49=2.19; p=0.033); N9-5 (t66<1); N8-7 (t44=2.33; p=0.024); N7-15 (t59<1); N5-8 (t51=1.45; p=0.15); N5-11 (t70<1); N5-10 (t55<1). In examining D1 Rotations following SAL (i.p.), none of the other lines besides N6-1 showed a significant genotypic difference (p_adjusted_<0.00625): N9-8 (t49=1.50;p=0.14); N9-5 (t66<1); N8-7 (t44=1.44;p=0.16); N7-15 (t59<1); N5-8 (t51<1); N5-11 (t70<1); N5-10 (t55<1). In examining D1 Spins following SAL (i.p.), none of the recombinant lines showed a significant genotypic difference (p_adjusted_<0.00625) (**Fig.4D**; vertically below panel C: N6-1 t48=2.13; p=0.038); N9-8 (t49=1.26; p=0.21); N9-5 (t66<1); N8-7 (t44=1.98; p=0.082); N7-15 (t59<1); N5-8 (t51<1); N5-11 (t70=1.03; p=0.31); N5-10 (t55=1.83; p=0.073).

In examining D2 OXY Distance, none of the other lines besides N6-1 and N6-8 showed a significant genotypic difference (p_adjusted_<0.05/8 recombinant lines=0.00625): N9-5 (t66=1.49; p=0.14); N8-7 (t44=2.72; p=0.0092); N7-15 (t59<1); N5-8 (t51=2.28; p=0.027); N5-11 (t70<1); N5-10 (t55<1). In examining D2 OXY rotations, again, none of the other lines besides N6-1 and N9-8 showed a significant genotypic difference (p_adjusted_<0.00625): N9-5 (t66=1.74; p=0.087); N8-7 (t44=1.58; p=0.12); N7-15 (t59<1); N5-8 (t51=1.72; p=0.092); N5-11 (t70<1); N5-10 (t55<1). In examining D2 OXY spins, none of the other lines besides N6-1 and N9-8 showed a significant genotypic difference (p_adjusted_<0.00625): N9-5 (t66=2.089; p=0.041); N8-7 (t44=1.88; p=0.066); N7-15 (t59<1); N5-8 (t51=1.70; p=0.095); N5-11 (t70<1); N5-10 (t55<1).

In addition to unpaired t-tests, because we subsequently discovered in the F2 analysis that the distal chromosome 1 QTL for OXY locomotor traits was driven by the females, we also ran two-way ANOVA analysis of D1 (SAL) and D2 (OXY) locomotor traits in each recombinant line and followed up specifically on those lines and traits showing a Genotype x Sex interaction. The only instances in which we identified recombinant lines showing Genotype x Sex interactions were for D1 locomotor traits (**Fig.S10)**.

Both N8-7 and N9-5 trended toward partially capturing the QTL for decreased OXY-induced behaviors but did not reach the Bonferroni-adjusted threshold of p<0.00625. Given the large sample sizes employed for each of the three lines (**Fig.S4B**) and that statistical power is not a limitation (see Power Analysis above), one explanation is that multiple variants within the locus influence OXY sensitivity and that N9-8 captures all of them, whereas the proximally shorter N8-7 and N9-5 lines capture only some of them. Unfortunately, despite our extensive efforts in surveying over 100 additional potential markers, we were unable to validate any polymorphic variants within the 170.16-172.61 Mb interval for reasons that included repeated failure in PCR amplification and Sanger sequencing and discovering that several reported variants were monomorphic. The widespread failure in Sanger sequencing could be due to uncharacterized structural variation segregating within this region (e.g., segmental duplication; see the final section in the Results, “Genetic variants between substrains within the causal QTL interval on distal chromosome 1”).

**Rgs7 as a candidate gene for OXY locomotion**

Although *Rgs7* (regulator of G-protein signaling, 7) is located distal to 170.16-172.61 Mb at 175.06 Mb, we would be remiss if we did not entertain *Rgs7* as a candidate gene for OXY locomotion and withdrawal. RGS proteins (regulators of g-protein signaling) serve as brakes on GPCR signaling and accordingly, *Rgs7* knockouts showed increase morphine locomotion, reward, reinforcement, and withdrawal (Sutton *et al.* 2016). Consistent with these observations, we found a non-significant increase in RGS7 protein in the J/N genotype (p = 0.08) that showed decreased OXY locomotion and withdrawal. Although *Rgs7* lies outside of the 2.45-Mb region, it is possible that one or more regulatory variants within the 2.45 Mb locus could modulate *Rgs7* expression (Smemo *et al.* 2014).

**SUPPLEMENTARY TABLES**

**Table S1: Polymorphic markers used for QTL mapping in B6J x B6NJ-F_2_ mice.** The marker dataset has previously been published (Kirkpatrick *et al.* 2017). Physical Map based on GRCm38 mouse reference build mm10.

| **Chr** | **Position (bp)** | **cM** | **B6J** | **B6NJ** | **Type** | **dbSNP** | **Target** | **% No Call** |
| --- | --- | --- | --- | --- | --- | --- | --- | --- |
| 1 | 29,549,273 | 11.33 | G | A | SNP | rs263412184 | Intergenic | 0.85 |
| 1 | 59,847,167 | 30.44 | G | A | SNP | rs227085647 | *Bmpr2* | 8.90 |
| 1 | 88,806,692 | 44.98 | C | T | SNP | rs242608911 | Intergenic | 0.61 |
| 1 | 116,037,566 | 51.85 | A | C | SNP | rs32180662 | *Cntnap5a* | 3.29 |
| 1 | 163,132,699 | 72.42 | A | T | SNP | rs6341208 | Intergenic | 0.37 |
| 1 | 181,318,003 | 84.58 | A | C | SNP | rs51237371 | Intergenic | 1.83 |
| 2 | 24,187,191 | 16.24 | A | G | SNP | rs254201911 | *Il1f9* | 6.46 |
| 2 | 38,946,612 | 24.43 | A | G | SNP | rs33064547 | Intergenic | 2.44 |
| 2 | 60,051,388 | 34.36 | A | T | SNP | rs256008253 | *Baz2b* | 0.49 |
| 2 | 78,799,176 | 47.11 | C | T | SNP | rs33162749 | Intergenic | 1.34 |
| 2 | 138,480,020 | 68.27 | T | C | SNP | rs13476801 | Intergenic | 3.66 |
| 2 | 152,781,403 | 75.36 | A | G | SNP | rs217443774 | *Bcl2l1* | 3.54 |
| 2 | 179,627,071 | 102.04 | T | C | SNP | rs3697302 | *Cdh4* | 38.41 |
| 3 | 5,370,727 | 1.85 | C | T | SNP | rs13476956 | Intergenic | 0.49 |
| 3 | 23,824,920 | 9.23 | T | A | SNP | rs13477019 | Intergenic | 0.61 |
| 3 | 54,914,906 | 25.91 | A | G | SNP | rs30751697 | *6030405A18Rik* | 10.12 |
| 3 | 95,734,876 | 41.34 | A | G | SNP | rs13474735 | *Ecm1* | 0.37 |
| 3 | 119,108,318 | 52.04 | A | G | SNP | rs50526143 | *Dpyd* | 0.85 |
| 3 | 159,811,101 | 82.64 | G | T | SNP | rs260058764 | Intergenic | 19.39 |
| 4 | 21,873,684 | 9.22 | C | G | SNP | rs13460207 | *Sfrs18* | 2.68 |
| 4 | 40,805,010 | 20.58 | A | G | SNP | rs46850777 | *B4galt1* | 2.20 |
| 4 | 65,944,235 | 34.50 | T | C | SNP | rs13477746 | Intergenic | 0.98 |
| 4 | 109,874,495 | 51.36 | A | G | SNP | rs3680956 | Intergenic | 0.24 |
| 4 | 124,654,566 | 57.85 | T | G | SNP | rs32865425 | *Pou3f1* | 2.93 |
| 4 | 138,221,673 | 70.22 | C | T | SNP | rs234702424 | *Hp1bp3* | 0.61 |
| 4 | 150,685,581 | 81.46 | T | C | SNP | rs32407463 | Intergenic | 2.56 |
| 5 | 18,710,388 | 8.37 | A | G | SNP | rs33367397 | Intergenic | 2.80 |
| 5 | 40,761,789 | 21.97 | C | T | SNP | rs33508711 | Intergenic | 19.15 |
| 5 | 59,833,642 | 31.78 | T | C | SNP | rs33209545 | Intergenic | 0.24 |
| 5 | 70,931,531 | 37.56 | T | C | SNP | rs29547790 | Intergenic | 11.10 |
| 5 | 89,775,351 | 44.34 | C | T | SNP | rs262569844 | *Adamts3* | 45.85 |
| 5 | 131,541,036 | 70.96 | A | G | SNP | rs3719791 | *Auts2* | 2.44 |
| 5 | 149,026,872 | 89.13 | T | C | SNP | rs33404968 | Intergenic | 8.90 |
| 6 | 39,400,456 | 18.21 | T | A | SNP | rs30899669 | *Mkrn1* | 1.34 |
| 6 | 60,591,379 | 29.06 | A | G | SNP | rs13478783 | Intergenic | 1.46 |
| 6 | 90,159,627 | 40.07 | A | C | SNP | rs245539697 | Intergenic | 1.10 |
| 6 | 117,470,880 | 55.04 | C | G | SNP | rs13478995 | Intergenic | 2.44 |
| 6 | 145,246,710 | 77.35 | A | G | SNP | rs30119689 | *Kras* | 1.71 |
| 7 | 4,006,930 | 2.30 | A | G | SNP | rs31995355 | Intergenic | 2.21 |
| 7 | 30,306,475 | 17.41 | A | G | SNP | rs3148686 | *Clip3* | 2.32 |
| 7 | 56,131,292 | 33.44 | G | A | SNP | rs50155421 | *Herc2* | 67.80 |
| 7 | 71,816,909 | 41.11 | C | G | SNP | rs32060039 | Intergenic | 0.98 |
| 7 | 102,973,309 | 54.80 | C | T | SNP | rs243575509 | Intergenic | 0.49 |
| 7 | 120,135,179 | 64.46 | G | A | SNP | rs235606231 | *Zp2* | 1.22 |
| 8 | 14,127,388 | 6.54 | C | T | SNP | rs236928053 | *Dlgap2* | 0.49 |
| 8 | 29,097,075 | 16.52 | T | C | SNP | rs13479672 | Intergenic | 2.68 |
| 8 | 59,125,561 | 30.78 | T | C | SNP | rs245638874 | Intergenic | 5.37 |
| 8 | 85,365,179 | 41.58 | A | T | SNP | rs245783224 | *Mylk3* | 4.27 |
| 8 | 119,945,761 | 68.67 | C | T | SNP | rs230879074 | *Gm20388* | 3.90 |
| 9 | 3,994,759 | 0.39 | C | T | SNP | rs226723272 | Intergenic | 0.98 |
| 9 | 25,130,622 | 10.16 | C | G | SNP | rs217366063 | *Herpud2* | 0.61 |
| 9 | 58,886,107 | 31.82 | G | A | SNP | rs238448868 | *Neo1* | 19.02 |
| 9 | 86,220,923 | 46.49 | A | C | SNP | rs30522453 | Intergenic | 1.83 |
| 9 | 120,016,907 | 71.41 | T | G | SNP | rs49019729 | *Xirp1* | 0.85 |
| 10 | 29,612,666 | 16.69 | C | A | SNP | rs29330320 | Intergenic | 0.49 |
| 10 | 67,238,174 | 34.92 | T | C | SNP | rs13480628 | *Jmjd1c* | 10.98 |
| 10 | 88,091,833 | 43.81 | T | C | SNP | rs246274290 | *4930547N16Rik* | 1.10 |
| 10 | 109,378,627 | 57.03 | C | T | SNP | rs13480759 | Intergenic | 71.95 |
| 11 | 30,675,691 | 18.15 | A | C | SNP | rs48169870 | Intergenic | 13.05 |
| 11 | 46,222,615 | 27.63 | G | A | SNP | rs240617401 | *Cyfip2* | 9.63 |
| 11 | 63,073,481 | 38.95 | T | C | SNP | rs6268547 | *Tekt3* | 24.15 |
| 11 | 90,480,671 | 55.09 | C | T | SNP | rs238893157 | *Stxbp4* | 0.98 |
| 11 | 120,306,788 | 84.05 | T | C | SNP | rs49027247 | Intergenic | 0.49 |
| 12 | 28,409,763 | 10.93 | G | T | SNP | rs29165094 | Intergenic | 0.61 |
| 12 | 55,758,373 | 24.10 | G | T | SNP | rs6385807 | Intergenic | 0.61 |
| 12 | 91,595,674 | 44.60 | A | C | SNP | rs235238684 | Intergenic | 0.61 |
| 12 | 117,167,625 | 62.92 | C | T | SNP | rs229717662 | *Ptprn2* | 0.49 |
| 13 | 27,037,150 | 12.24 | A | G | SNP | rs13481734 | Intergenic | 1.59 |
| 13 | 52,758,052 | 27.35 | T | C | SNP | rs29248623 | Intergenic | 17.93 |
| 13 | 72,630,777 | 39.25 | T | C | SNP | rs29957076 | *Irx2* | 34.27 |
| 13 | 88,426,920 | 45.27 | G | A | SNP | rs29622109 | Intergenic | 19.88 |
| 13 | 119,143,521 | 67.25 | C | T | SNP | rs29235721 | Intergenic | 1.22 |
| 14 | 9,937,385 | 5.80 | G | A | SNP | rs31187642 | Intergenic | 2.93 |
| 14 | 74,415,721 | 39.25 | T | A | SNP | rs30264676 | Intergenic | 2.07 |
| 14 | 90,847,095 | 45.14 | G | T | SNP | rs262164631 | Intergenic | 10.24 |
| 14 | 116,603,597 | 60.23 | A | C | SNP | rs236603300 | Intergenic | 1.10 |
| 15 | 11,336,383 | 5.58 | G | T | SNP | rs246033409 | *Adamts12* | 2.68 |
| 15 | 37,462,413 | 14.99 | A | T | SNP | rs31810918 | Intergenic | 0.85 |
| 15 | 71,632,551 | 32.31 | T | C | SNP | rs31858887 | Intergenic | 5.12 |
| 16 | 12,261,912 | 7.23 | T | C | SNP | rs4162529 | *Shisa9* | 0.61 |
| 16 | 35,291,544 | 25.15 | G | A | SNP | rs258334795 | *Adcy5* | 0.85 |
| 16 | 60,954,616 | 36.06 | C | G | SNP | rs4193066 | Intergenic | 3.66 |
| 16 | 90,181,009 | 51.59 | C | G | SNP | rs4217166 | Intergenic | 3.54 |
| 17 | 5,332,903 | 3.15 | T | C | SNP | rs4137196 | *Arid1b* | 0.12 |
| 17 | 26,854,246 | 13.56 | A | T | SNP | rs46703123 | Intergenic | 0.61 |
| 17 | 67,752,883 | 38.88 | G | A | SNP | rs33387924 | *Lama1* | 1.46 |
| 17 | 89,708,416 | 59.66 | C | T | SNP | rs50807504 | Intergenic | 38.41 |
| 18 | 9,450,891 | 4.87 | C | G | SNP | rs29560146 | Intergenic | 2.37 |
| 18 | 31,018,871 | 17.70 | G | A | SNP | rs257888249 | *Rit2* | 32.32 |
| 18 | 54,614,841 | 29.22 | A | C | SNP | rs13483369 | Intergenic | 20.85 |
| 18 | 90,448,757 | 59.64 | A | G | SNP | rs263687961 | Intergenic | 1.10 |
| 19 | 9,027,005 | 6.04 | G | C | SNP | rs31112038 | *Ahnak* | 1.34 |
| 19 | 16,602,218 | 11.66 | T | C | SNP | rs30709918 | Intergenic | 2.44 |
| 19 | 52,359,370 | 46.77 | A | G | SNP | rs30608930 | Intergenic | 0.12 |
| X | 64,312,439 | 34.62 | C | T | SNP | rs246222509 | Intergenic | 0.37 |
| X | 90,719,435 | 40.39 | A | C | SNP | rs221876655 | Intergenic | 4.63 |
| X | 120,419,803 | 49.63 | T | A | SNP | rs226119295 | *Pcdh11x* | 1.83 |
| X | 152,774,935 | 70.34 | A | G | SNP | rs226776950 | intergenic | 0.61 |

**Table S2. Polymorphic markers used to identify recombination events and fine map the 2.45 Mb distal chromosome 1 locus (170.16-172.61 Mb) underlying OXY behavioral sensitivity.**

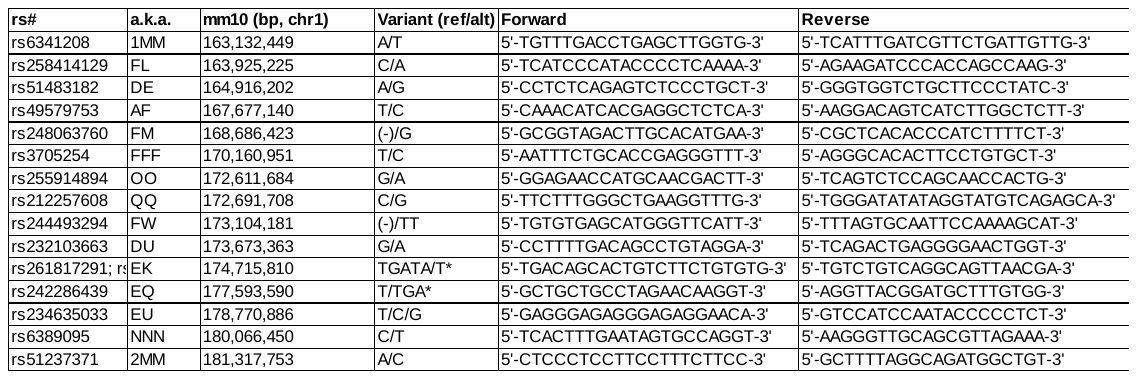

**Table S3. Genotypes at each marker for each recombinant line.**

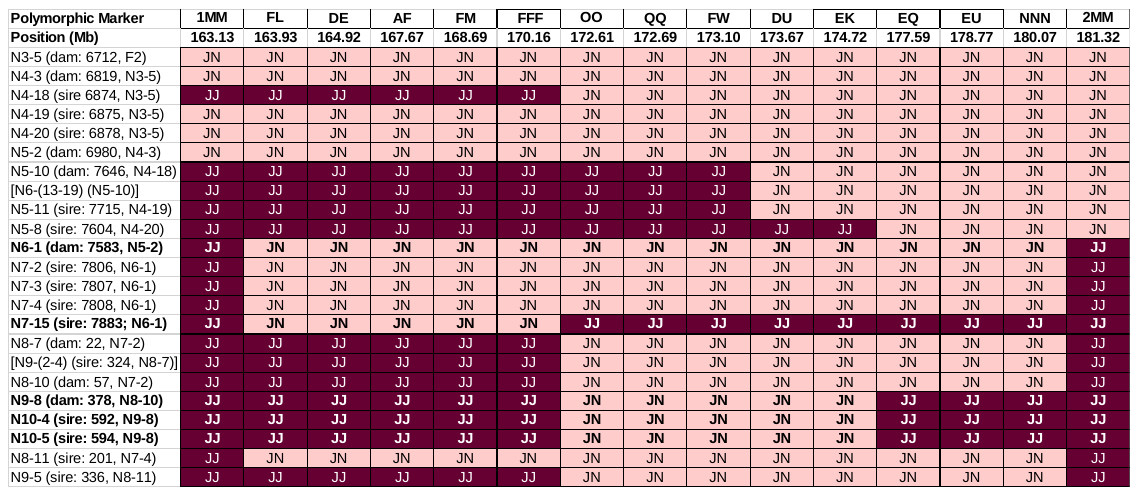

**Table S4: Cis-eQTLs associated with peak distal chromosome 1 marker for OXY-induced locomotor activity (rs51237371; 181,318,003 bp).**

| **geneID** | **chr** | **start** | **end** | **AvExp** | **F** | **P.Value** | **adj.P.Val** |
| --- | --- | --- | --- | --- | --- | --- | --- |
| Epb41l5 | 1 | 119545037 | 119649000 | 3.44 | 2.84 | 0.005263 | 0.044089 |
| Lypd1 | 1 | 125867622 | 125913101 | 4.45 | 3.51 | 0.000631 | 0.009675 |
| Tmem163 | 1 | 127486558 | 127679548 | 3.33 | -3.93 | 0.000138 | 0.003071 |
| R3hdm1 | 1 | 128103321 | 128237736 | 8.03 | -4.44 | 1.94E-05 | 0.00072 |
| Pfkfb2 | 1 | 130689182 | 130729253 | 5.94 | 3.26 | 0.001415 | 0.017428 |
| Rassf5 | 1 | 131176410 | 131245258 | 4.38 | -3.33 | 0.001151 | 0.015043 |
| Mfsd4a | 1 | 132022806 | 132067965 | 5.05 | -2.93 | 0.004075 | 0.036462 |
| Dstyk | 1 | 132417555 | 132466958 | 6.79 | 2.85 | 0.005084 | 0.042958 |
| Mdm4 | 1 | 132959484 | 133030561 | 7.22 | 6.19 | 7.64E-09 | 1.49E-06 |
| Btg2 | 1 | 134075170 | 134079120 | 3.84 | 3.97 | 0.000121 | 0.002823 |
| Ppfia4 | 1 | 134296783 | 134332928 | 6.78 | 7.62 | 5.25E-12 | 5.04E-09 |
| Rabif | 1 | 134494648 | 134508774 | 4.95 | -3.19 | 0.001795 | 0.020552 |
| Arl8a | 1 | 135146824 | 135156269 | 7.15 | 4.32 | 3.09E-05 | 0.001011 |
| Shisa4 | 1 | 135371057 | 135375237 | 6.47 | -3.68 | 0.000346 | 0.006122 |
| Ipo9 | 1 | 135382312 | 135430499 | 7.18 | -3.96 | 0.000123 | 0.002839 |
| Phlda3 | 1 | 135766119 | 135769136 | 4.10 | -2.85 | 0.00517 | 0.043467 |
| Rgs2 | 1 | 143999338 | 144004161 | 6.86 | 2.81 | 0.005747 | 0.046866 |
| Ptgs2 | 1 | 150100031 | 150108227 | 2.76 | -3.06 | 0.002741 | 0.027671 |
| Ivns1abp | 1 | 151344477 | 151364422 | 8.58 | 5.32 | 4.65E-07 | 4.33E-05 |
| Trmt1l | 1 | 151428542 | 151458161 | 4.28 | 2.85 | 0.005134 | 0.043284 |
| Nmnat2 | 1 | 152954998 | 153119261 | 7.06 | -4.30 | 3.34E-05 | 0.001067 |
| Rgs8 | 1 | 153653025 | 153700323 | 8.16 | 3.79 | 0.000231 | 0.00449 |
| Glul | 1 | 153899944 | 153909723 | 10.84 | -5.29 | 5.18E-07 | 4.72E-05 |
| Cacna1e | 1 | 154390731 | 154864067 | 9.34 | 3.40 | 0.000897 | 0.012569 |
| Cep350 | 1 | 155844964 | 155973255 | 6.29 | 4.82 | 4.02E-06 | 0.000221 |
| Brinp2 | 1 | 158245269 | 158356326 | 6.39 | -5.11 | 1.18E-06 | 9.08E-05 |
| Tnr | 1 | 159523769 | 159931729 | 7.68 | -3.22 | 0.001644 | 0.019351 |
| Rc3h1 | 1 | 160906418 | 160974978 | 6.10 | 3.13 | 0.002171 | 0.023654 |
| Kifap3 | 1 | 163779583 | 163917109 | 7.63 | -4.38 | 2.51E-05 | 0.000863 |
| Gpr161 | 1 | 165295789 | 165326745 | 3.50 | 3.27 | 0.00137 | 0.017042 |
| Pou2f1 | 1 | 165865154 | 166002678 | 5.37 | 4.96 | 2.24E-06 | 0.000144 |
| Ildr2 | 1 | 166254139 | 166316823 | 6.66 | -3.48 | 0.000698 | 0.010434 |
| Nuf2 | 1 | 169497934 | 169531464 | 1.53 | 3.01 | 0.003179 | 0.030795 |
| Pcp4l1 | 1 | 171173262 | 171196268 | 8.64 | 4.81 | 4.17E-06 | 0.000227 |
| Ncstn | 1 | 172066013 | 172082795 | 5.72 | -3.48 | 0.00068 | 0.010214 |
| Atp1a2 | 1 | 172271709 | 172298064 | 9.74 | -4.03 | 9.55E-05 | 0.002328 |
| Kcnj9 | 1 | 172320501 | 172329318 | 6.74 | -3.34 | 0.001086 | 0.014392 |
| Igsf9 | 1 | 172481788 | 172498878 | 3.78 | 3.10 | 0.002408 | 0.025474 |
| Cadm3 | 1 | 173333258 | 173367695 | 8.63 | -4.43 | 2.05E-05 | 0.000747 |
| Aim2 | 1 | 173350879 | 173466040 | 1.84 | -3.16 | 0.001987 | 0.022229 |
| Rgs7 | 1 | 175059087 | 175492500 | 4.93 | -3.25 | 0.001475 | 0.017949 |
| Cep170 | 1 | 176733653 | 176814067 | 6.25 | -3.05 | 0.002753 | 0.027765 |
| Zbtb18 | 1 | 177442351 | 177450764 | 7.94 | -5.38 | 3.51E-07 | 3.45E-05 |
| Desi2 | 1 | 178187417 | 178257301 | 5.49 | 3.04 | 0.002889 | 0.028582 |
| Hnrnpu | 1 | 178324343 | 178337797 | 8.94 | -4.17 | 5.67E-05 | 0.00159 |
| Kif26b | 1 | 178529125 | 178939200 | 5.10 | 5.22 | 7.27E-07 | 6.18E-05 |
| Smyd3 | 1 | 178951960 | 179518041 | 4.68 | 3.88 | 0.000167 | 0.00351 |
| Cdc42bpa | 1 | 179960472 | 180165603 | 8.39 | 2.98 | 0.003488 | 0.0328 |
| H3f3a | 1 | 180800832 | 180813943 | 5.35 | 3.31 | 0.001229 | 0.015733 |
| C130074G19Rik | 1 | 184871926 | 184883218 | 5.72 | -3.75 | 0.000266 | 0.005041 |
| Slc30a10 | 1 | 185454848 | 185468762 | 5.40 | 4.50 | 1.53E-05 | 0.000595 |
| Mfsd7b | 1 | 191005847 | 191026158 | 3.97 | 3.08 | 0.002566 | 0.026583 |
| Dtl | 1 | 191537365 | 191575544 | 2.32 | -2.93 | 0.003997 | 0.036134 |
| Lpgat1 | 1 | 191717834 | 191784255 | 8.63 | -3.39 | 0.000947 | 0.013121 |
| Nek2 | 1 | 191821444 | 191833050 | 2.10 | 3.18 | 0.001845 | 0.020961 |
| Kcnh1 | 1 | 192190774 | 192510159 | 6.64 | -4.30 | 3.41E-05 | 0.001082 |
| Syt14 | 1 | 192891233 | 193035775 | 5.39 | 4.61 | 9.80E-06 | 0.000433 |
| Cd34 | 1 | 194938819 | 194961279 | 5.51 | -3.81 | 0.000219 | 0.004333 |

**Table S5. Chromosome 5 cis-eQTL transcripts showing peak association with the peak behavioral QTL EPM marker for EPM behaviors during spontaneous OXY withdrawal (rs33209545; chromosome 5: 59,833,642 bp).** Bolded genes indicate candidate genes flanking the peak QTL (59 Mb) associated with spontaneous OXY withdrawal.

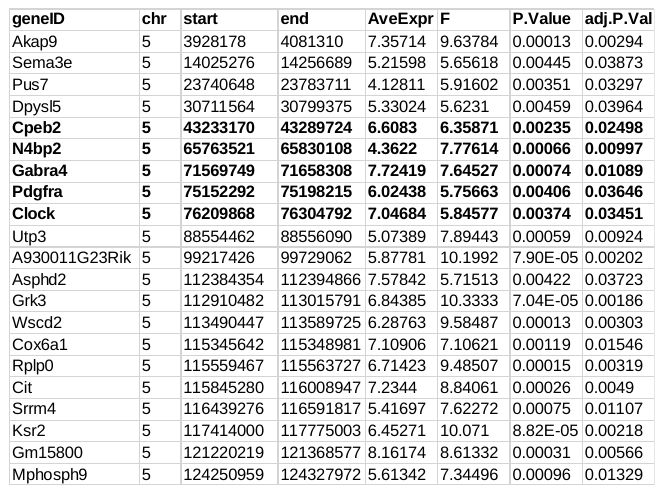

**Table S6. Genetic variants distinguishing C57BL/6J and C57BL/6NJ within the 2.45 Mb interval on distal chromosome 1 spanning 170.16-172.61 Mb.** REF = reference allele; ALT=alternate allele; QUAL=Phred-scaled variant probability; INFO = variant type (e.g., SNP or INDEL) and read depth (DP); FORMAT=format of genotype calls for each sample; GT = genotype of variant (*e.g*., 0/0, 0/1, or 1/1); PL: Normalized Phred-scaled likelihood for each possible genotypes (e.g., 0/0, 0/1, 1/1) with the most likely genotype given a score of 0 and the other genotypes scaled relative to the most likely genotype. 0/0=homozygous reference allele; 1/1=homozygous alternate allele. (please zoom in to view text. **In case the resolution is not sufficient once the PDF has been built, please also see Table S6 supplied as an excel spreadsheet**).

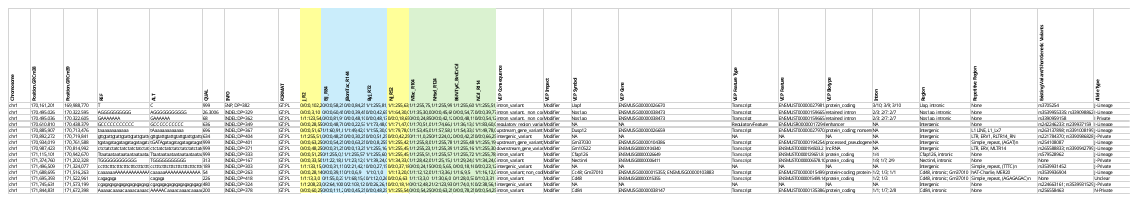

**SUPPLEMENTARY FIGURES**

**Fig.S1. D3 and D5 distance traveled (SAL training trials) in B6 substrains. (A):** For D3, there was an effect of Dose (F3,293=11.84; ****p<0.0001), but no effect of Substrain (F1,293=1.03; p=0.311) and no interaction (F3,293=2.24; p=0.084). The effect of Dose was explained by mice with prior exposure to the two highest OXY doses during training (1.25, 5 mg/kg) showing greater locomotor activity in response to SAL injections compared to mice receiving 0 mg/kg (***p<0.001; ****p<0.0001, respectively) (Bonferroni). The fact that these observations appear to show a dose-dependency and were only observed at the two highest doses suggest that there is a learned, conditioned locomotion effect. **(B):** To confirm a lack of Substrain effect in response to SAL injection on D3, we examined the time course in 5-min bins over 30 min in the group with prior exposure to 1.25 mg/kg OXY. Although there was an effect of Time (F5,150=96.30; p<0.0001), there was no effect of Substrain (F1,30)=1.95; p=0.17) and no interaction (F5,150=2.01; p=0.081). **(C):** In examining locomotor activity following the second SAL training day on D5, there was again a Dose effect (F3,292=4.43; p=0.0047) but no Substrain effect (F1,292=1.06; p=0.30) and no interaction (F3,292<1). The Dose effect was explained by an increase at 1.25 mg/kg vs. 0 mg/kg (**p<0.01), an increase at 1.25 mg/kg vs. 0.625 mg/kg (*p<0.05) (Bonferroni). These data indicate a persistent conditioned locomotion effect with the 1.25 mg/kg dose on drug-free training days and provide further support that the Substrain effect on locomotor activity on D2 and D4 is specific to OXY treatment. **(D):** Further support for this contention comes from time course analysis of D5 distance in the group with 2x prior exposure to the 1.25 mg/kg OXY. Here, there was no main effect of Substrain (F1,30<1). However, there was an effect of Time (F5,150 = 84.36; ****p < 0.0001) and a Substrain x Time interaction (F5,150 = 4.34; **p < 0.001) that was explained by a small decrease in B6NJ vs. B6J that was limited to the first 5-min bin of the 30 min assessment (*p<0.05) (Bonferroni).

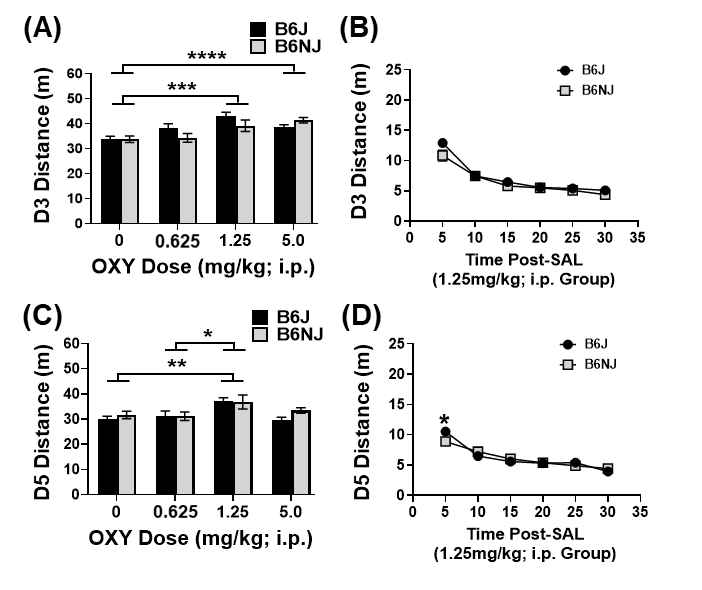

**Fig.S2. Effect plots for D2 and D4 OXY locomotor traits.** **(A-E):** Effect plots at rs51237371 (181.32 Mb) for distal chromosome 1 phenotypes showing significant QTLs. Note the dominant allelic effect of N, in which one copy of the N allele is sufficient to full decrease the OXY locomotor trait. Note also that the effect of the N allele is specific to OXY-treated mice and is not observed in SAL-treated mice.

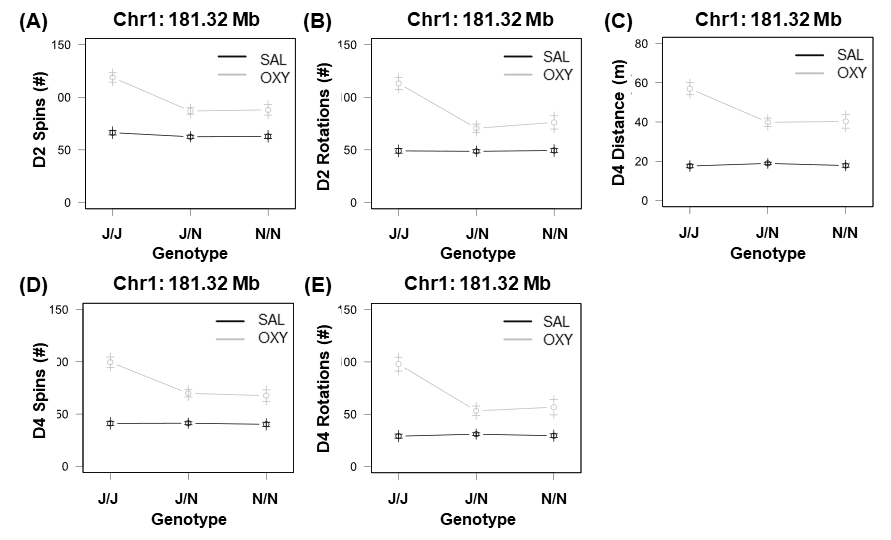

**Fig.S3. Additional QTL analyses to demonstrate specificity for OXY treatment. (A):** No significant chr.1 QTL for D3 and D5 Distance (SAL training trials). The suggestive peak for D3 Distance is likely explained by conditioned locomotor effects from D2 OXY injection (Fig.S1). **(B):** No genome-wide significant QTLs for D1 Distance (N=425 F2 mice; treatment-collapsed). **(C,D):** Analysis of SAL F2 mice only revealed no genome-wide significant QTLs for D2 or D4 OXY locomotor traits (C) or EPM traits (D).

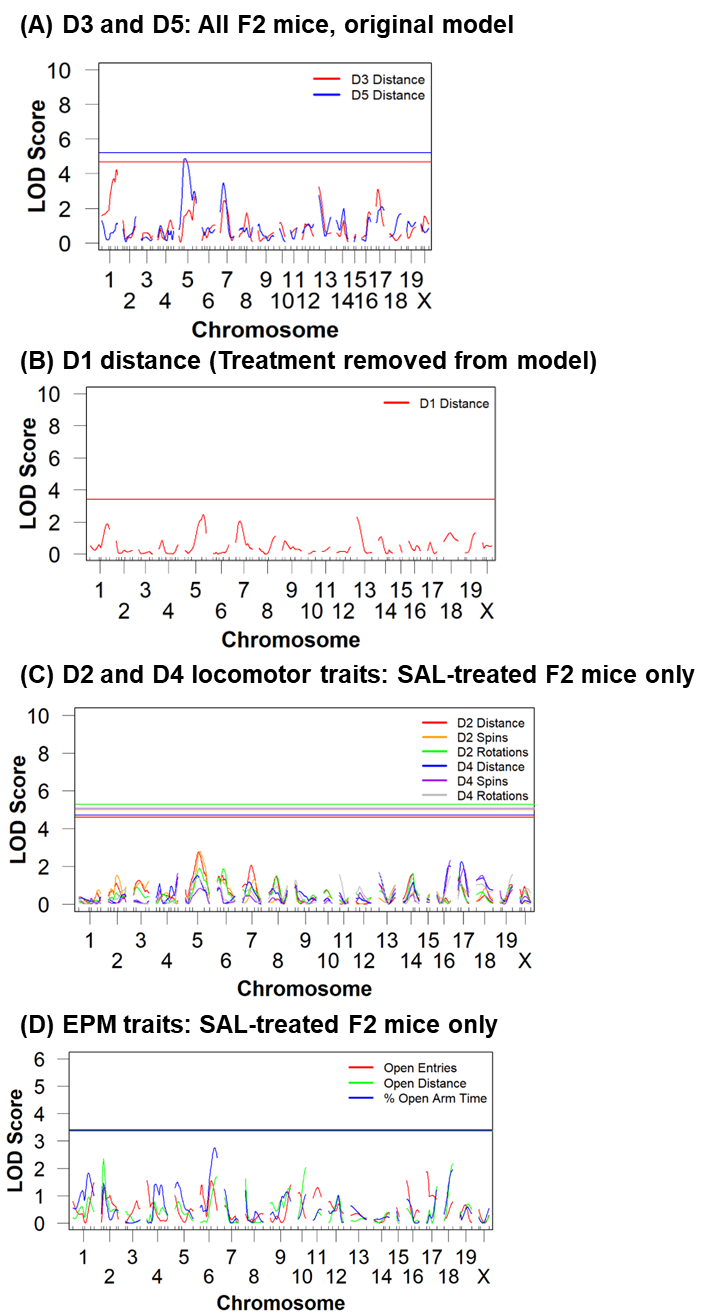

**Fig.S4. Antinociceptive tolerance in B6 parental substrains but not F2 mice following 4x 20 mg/kg OXY (i.p.) treatment and a challenge dose of OXY (5.0 mg/kg, i.p.).** **(A):** Experimentally-naïve C57BL/6 substrains underwent 4x 20 mg/kg OXY on Monday-Thurs of Week 3 (Fig.2A) and 16 h later were tested for baseline hot plate latency. Thirty min later, mice were injected with OXY (5 mg/kg, i.p.) and thirty min later, were then tested for post-injection antinociceptive latencies. There was a Treatment effect (F1,40=16.48; ***p<0.001) but no Genotype or Sex effects (F1,40<1) and no interactions (ps>0.05), indicating significant tolerance irrespective of Substrain or Sex. **(B):** CPP-experienced F2 mice were treated during Week 3 (Fig.1A) with 4X 20 mg/kg OXY (i.p.) on Monday-Thursday and 16 h later on Friday, were assessed identically as described in panel A for baseline nociception and OXY-induced antinociception. Unlike the C57BL/6 substrains, F2 mice showed less OXY-induced antinociception did not exhibit significant antinociceptive tolerance, as there were no Treatment effect (F1,156<1), Sex effect (F1,156=1.53; p=0.22), or interaction (F1,156=2.15; p=0.15).

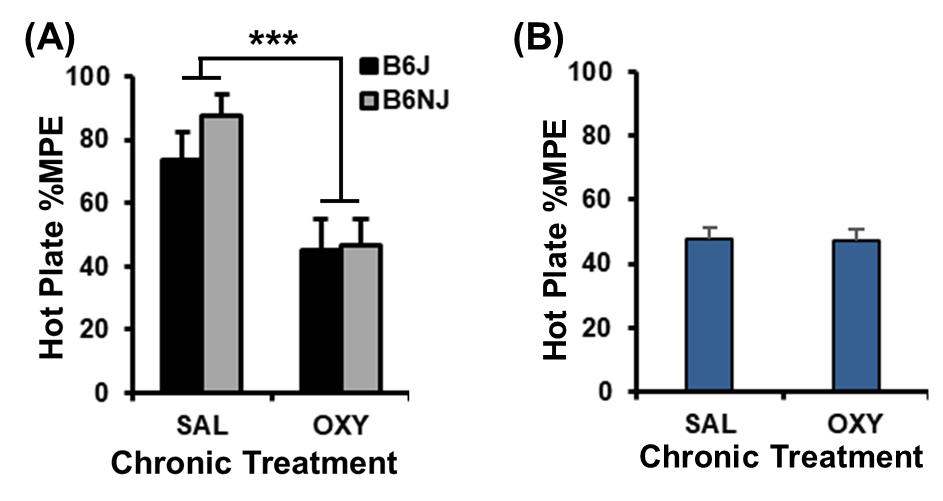

**Fig.S5. OXY withdrawal traits in F2 mice and effect plots at distal chromosome 1 QTL.** Means and S.E.M. for EPM traits. **(A):** For Open Arm Entries (#), there was no Treatment effect (F1,228=3.45; p=0.065), an effect of Sex (F1,228=3.95; p=0.048), and no interaction (F1,228<1). **(B):** For Open Arm Distance (m), there was no effect of Treatment (F1,228<1), Sex (F1,228<1), or interaction (F1,228<1). **(C):** For Open Arm Time (%; panel B), there was no effect of Treatment (F1,228<1), Sex (F1,228<1), or interaction (F1,228<1). **(D,E):** Effect plots for distal chromosome 1 EPM withdrawal traits showing significant QTLs (see Figure 2B,F-H).

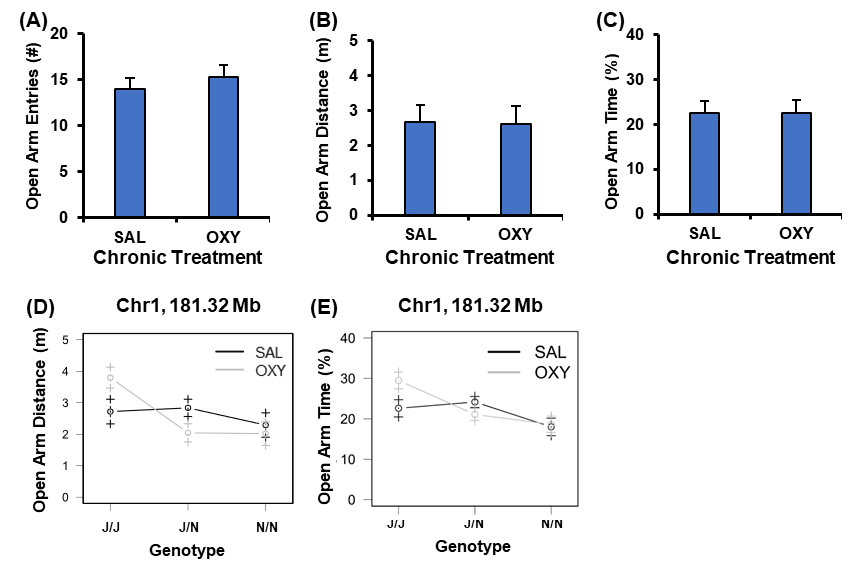

**Fig.S6. QTL analysis of EPM withdrawal traits in hot plate-exposed F2 mice only.** A genome-wide significant QTL was once again identified on distal chromosome 1. While the medial chromosome 5 QTL for Open Arm Entries (Fig.2B,I,J) was not significant, the peak is still visible without inclusion of the hot plate-naïve mice.

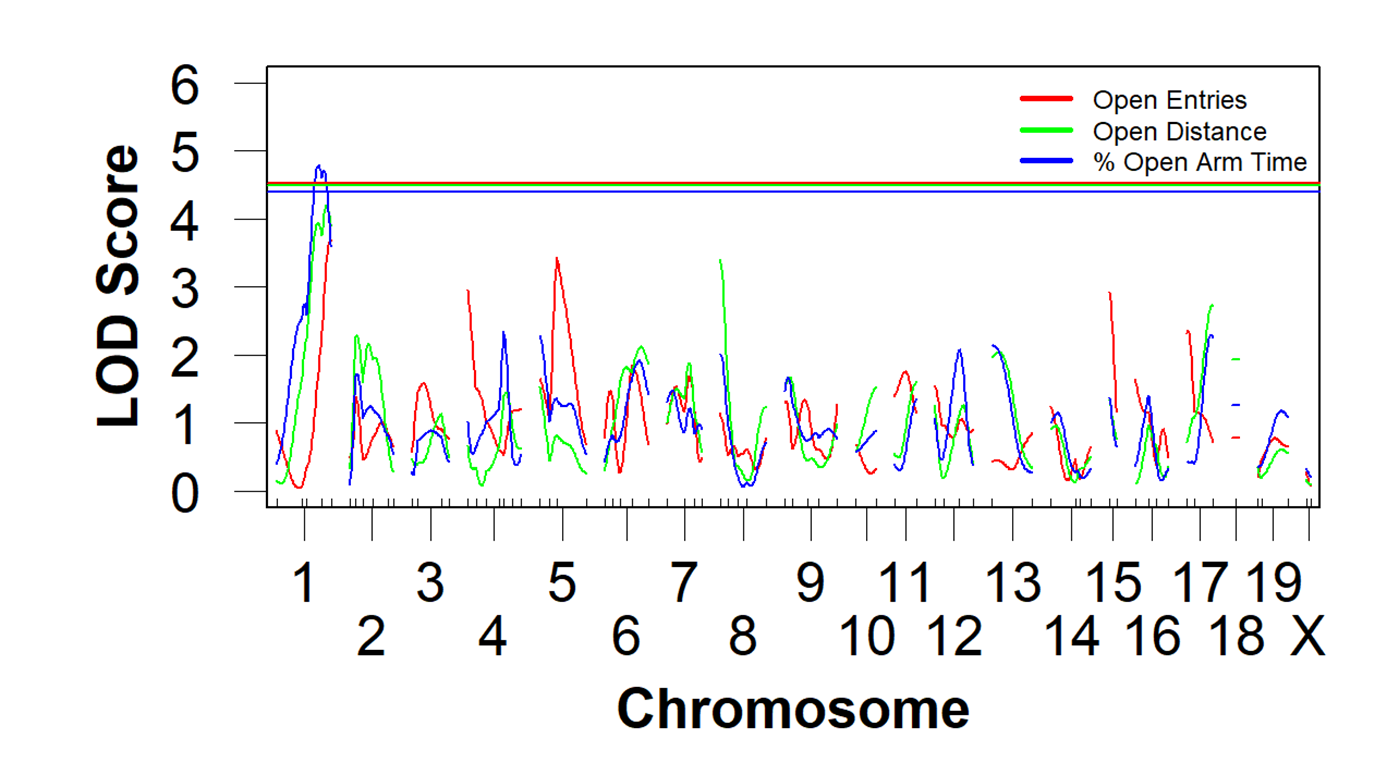

**Fig.S7. QTL analysis by Sex for OXY locomotor traits and EPM withdrawal traits**

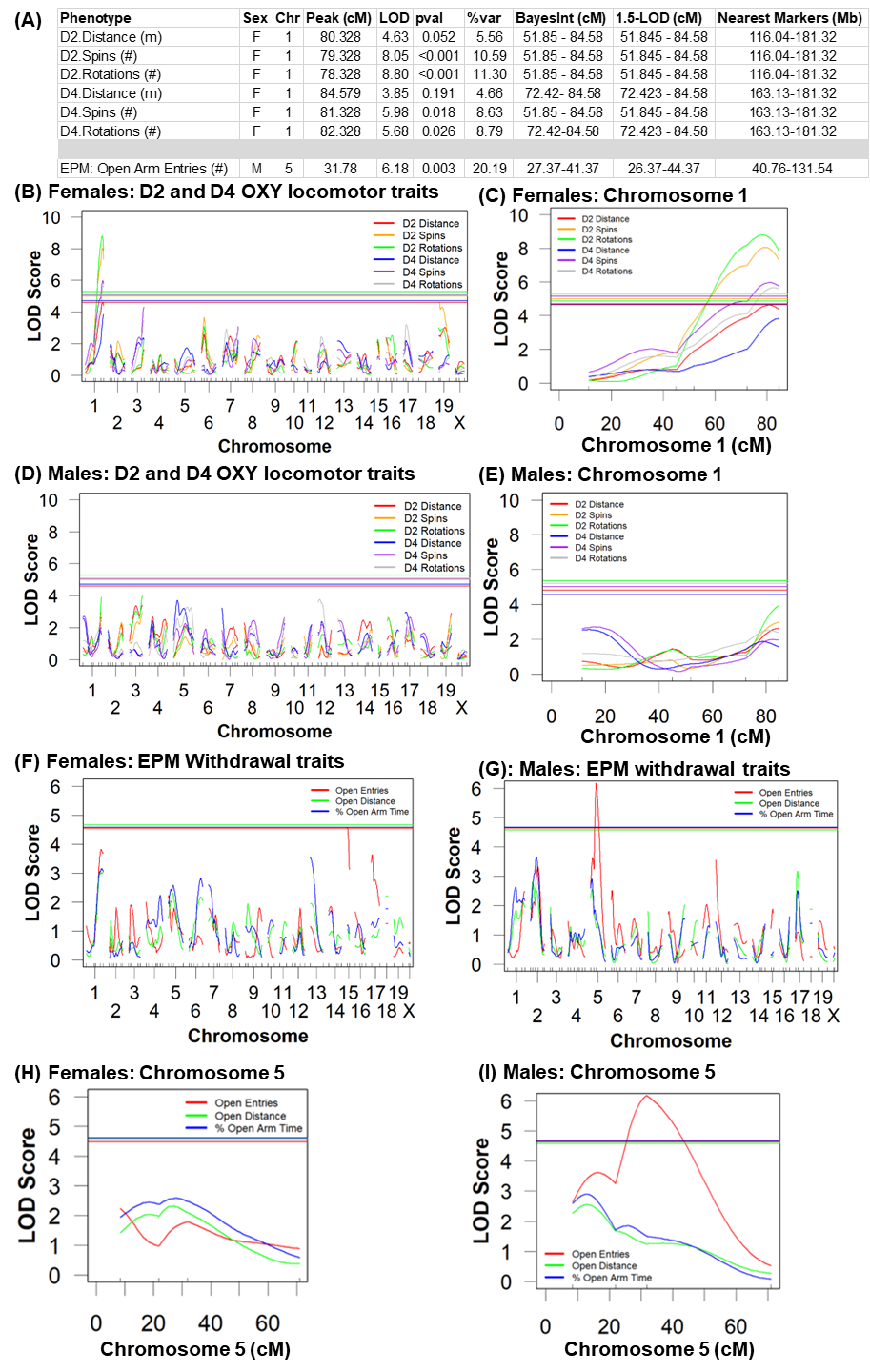

**Fig.S8. Effect plots by Sex for D2 and D4 OXY locomotor traits and EPM withdrawal traits.**

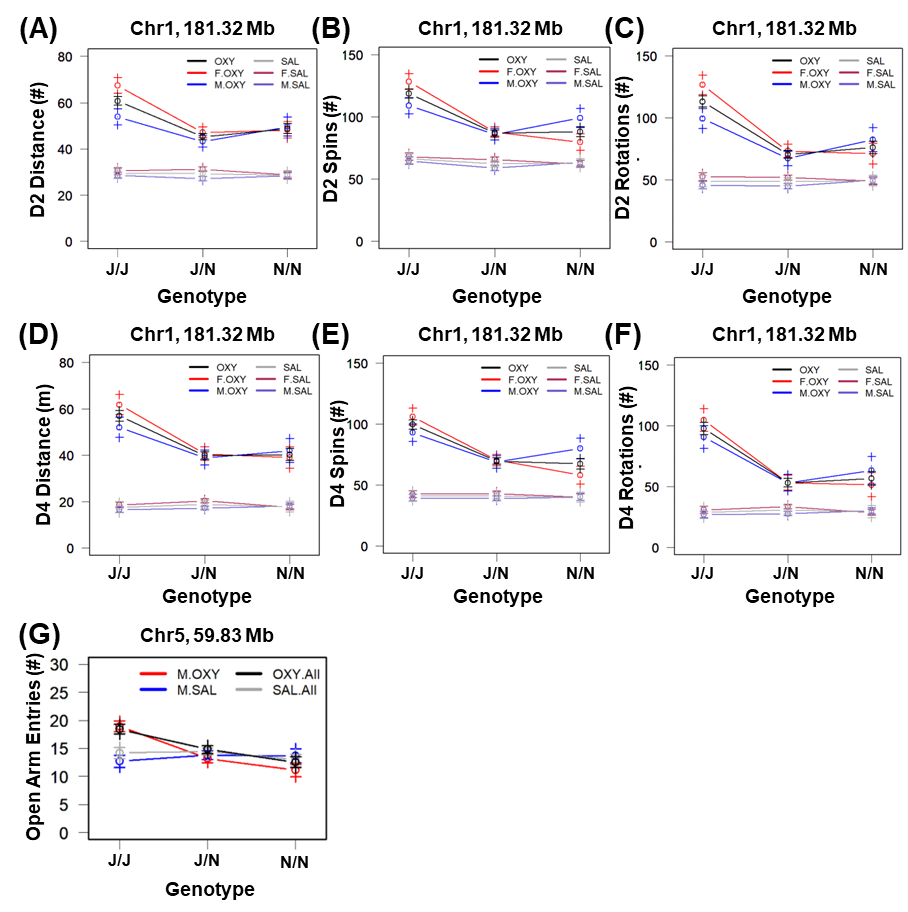

**Fig.S9. Pedigree for recombinant lines used to fine-map the distal chromosome 1 QTL for OXY behavioral sensitivity. (A):** Pedigree for generating and propagating recombinant lines. Beginning with the single F2 founder, mice were continually backrossed at each generation to propagate informative sample sizes for a particular recombinant event within a line and to screen for new informative recombination events that could also then also be propagated and aid in fine-mapping the QTL interval. **Underlined, bolded:** Recombinant lines that were phenotyped. N7-15, N9-5, and N5-8 all produced sufficient offspring that further propagation with additional families was not necessary. N9-8, N8-7, N5-10, and N5-11 were all backcrossed to continue making mice with the same recombinant interval within the 170.16-170.61 Mb region and thus, within each line, these data were combined for the analysis. **(B):** Schematic (same as in Figure 4a) of the eight congenic lines used to deduce the 2.45 Mb region spanning 170.16-172.61 Mb. Burgundy color: homozygous J/J genotype. Mauve color: heterozygous J/N genotype “+” = captured the QTL for reduced D2 OXY distance. “-” = failed to capture the QTL for reduced D2 OXY distance.

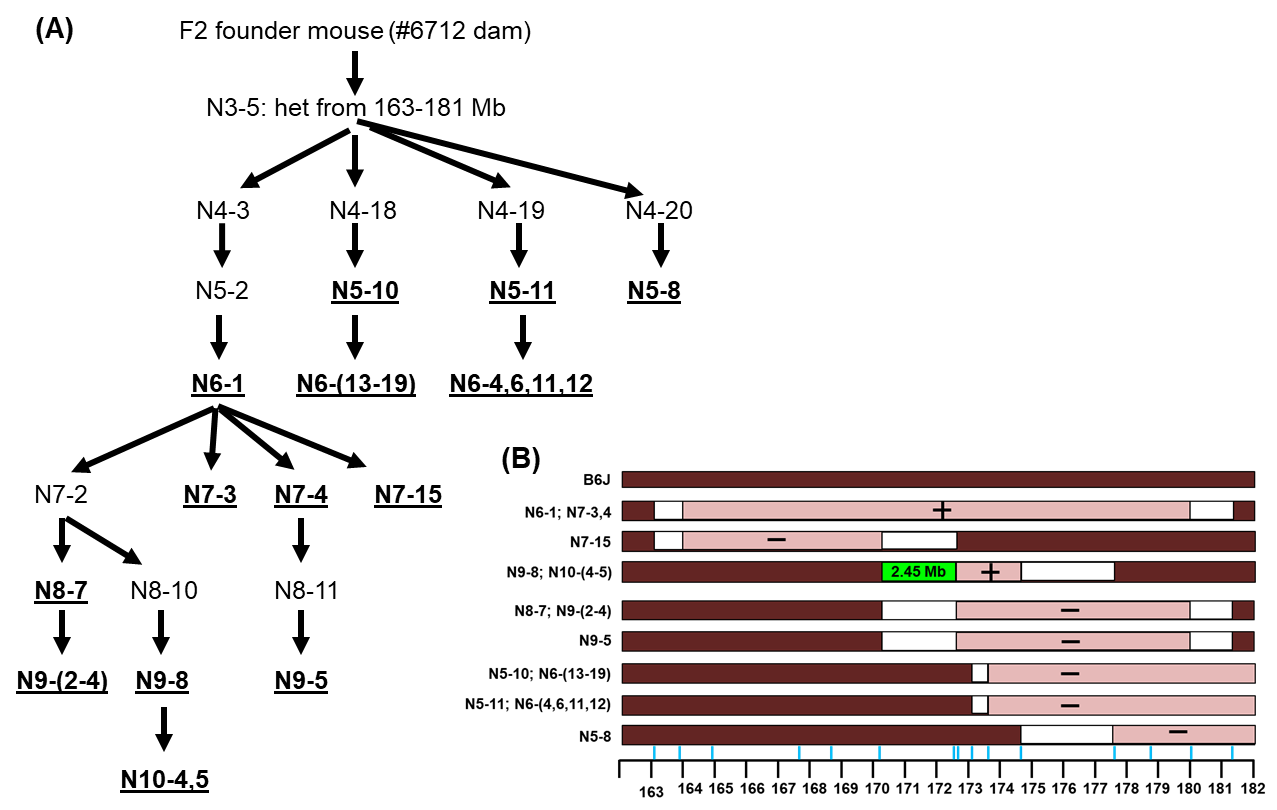

**Fig.S10. Genotype x Sex interactions and post-hoc analyses of D1 locomotor traits in reocombinant lines. (A): D1 Distance, N8-7:** Genotype x Sex: F1,42=6.19; p=0.017. Tukey’s: J/J females vs. J/N females: padj=0.0059. Males: n.s. **(B): D1 Spins, N8-7:** Genotype x Sex: F1,42=4.15; p=0.048. Tukey’s: J/J females vs. J/N females: padj=0.029. Males: n.s. **(C):** **D1 Spins, N9-5**: Genotype x Sex: F1,64=11.68 ; p=0.0011. Tukey’s: J/J females vs. J/N females: *padj=0.013. J/N females vs. J/N males: *p=0.013.

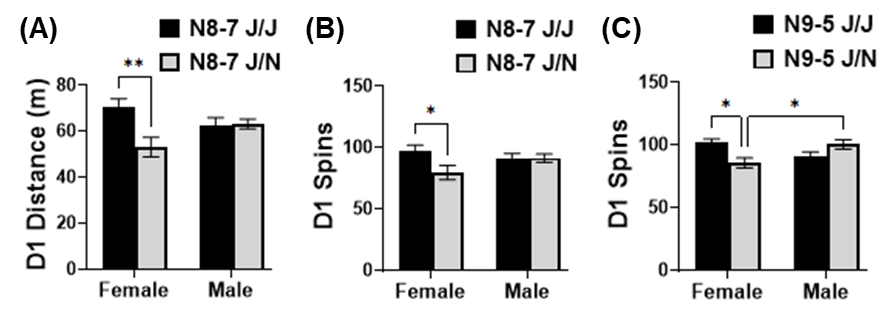

**Figure S11. Genes located within the 2.45 Mb interval on distal chromosome 1 spanning 170.16-172.63 Mb (mm10, UCSC Genome Browser).**

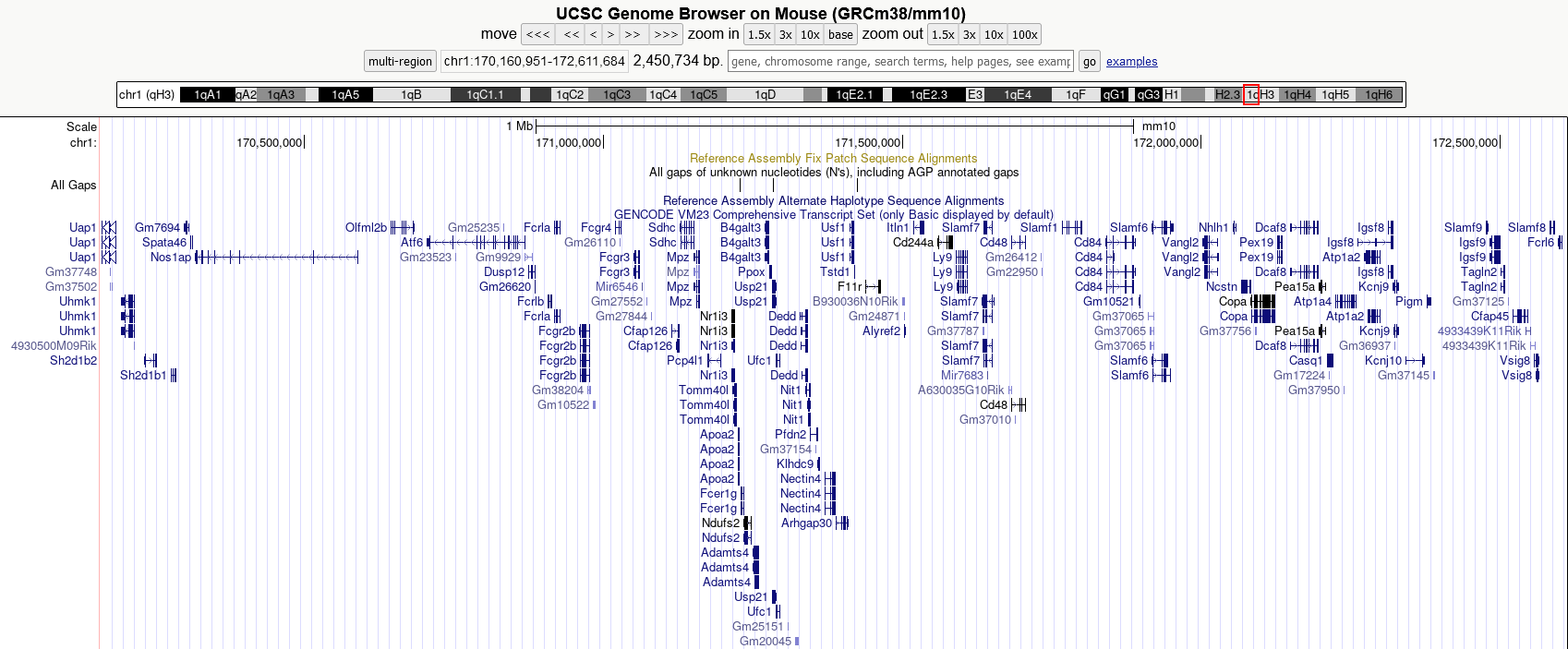

**REFERENCES**

Benjamini, Y. & Hochberg, Y. (1995) Controlling false discovery rate: A practical and powerful approach to multiple testing. *Journal of the Royal Statistical Society* **57**, 289–300.

Broman, K.W. & Sen, S. (2009) *A Guide to QTL Mapping with R/qtl*. , Statistics for Biology and Health. 1st edn. Springer-Verlag, Inc., New York.

Broman, K.W., Wu, H., Sen, S. & Churchill, G.A. (2003) R/qtl: QTL mapping in experimental crosses. *Bioinformatics (Oxford, England)* **19**, 889–90.

Bryant, C.D., Bagdas, D., Goldberg, L.R., Khalefa, T., Reed, E.R., Kirkpatrick, S.L., Kelliiher, J.C., Chen, M.M., Johnson, W.E., Mulligan, M.K. & Damaj, M.I. (2019) C57BL/6 substrain differences in inflammatory and neuropathic nociception and genetic mapping of a major quantitative trait locus underlying acute thermal nociception. *Mol Pain* 1744806918825046.

Goldberg, L.R., Yao, E.J., Kelliher, J.C., Reed, E.R., Wu Cox, J., Parks, C., Kirkpatrick, S.L., Beierle, J.A., Chen, M.M., Johnson, W.E., Homanics, G.E., Williams, R.W., Bryant, C.D. & Mulligan, M.K. (2021) A quantitative trait variant in Gabra2 underlies increased methamphetamine stimulant sensitivity. *BioRxiv* **450337**.

Keane, T.M., Goodstadt, L., Danecek, P., White, M.A., Wong, K., Yalcin, B., Heger, A., Agam, A., Slater, G., Goodson, M., Furlotte, N.A., Eskin, E., Nellaker, C., Whitley, H., Cleak, J., Janowitz, D., Hernandez-Pliego, P., Edwards, A., Belgard, T.G., Oliver, P.L., McIntyre, R.E., Bhomra, A., Nicod, J., Gan, X., Yuan, W., van der Weyden, L., Steward, C.A., Bala, S., Stalker, J., Mott, R., Durbin, R., Jackson, I.J., Czechanski, A., Guerra-Assuncao, J.A., Donahue, L.R., Reinholdt, L.G., Payseur, B.A., Ponting, C.P., Birney, E., Flint, J. & Adams, D.J. (2011) Mouse genomic variation and its effect on phenotypes and gene regulation. *Nature* **477**, 289–294.

Kirkpatrick, S.L. & Bryant, C.D. (2015) Behavioral architecture of opioid reward and aversion in C57BL/6 substrains. *FrontBehavNeurosci* **8**, 450.

Kirkpatrick, S.L., Goldberg, L.R., Yazdani, N., Babbs, R.K., Wu, J., Reed, E.R., Jenkins, D.F., Bolgioni, A.F., Landaverde, K.I., Luttik, K.P., Mitchell, K.S., Kumar, V., Johnson, W.E., Mulligan, M.K., Cottone, P. & Bryant, C.D. (2017) Cytoplasmic FMR1-Interacting Protein 2 Is a Major Genetic Factor Underlying Binge Eating. *BiolPsychiatry* **81**, 757–769.

Law, C.W., Chen, Y., Shi, W. & Smyth, G.K. (2014) voom: Precision weights unlock linear model analysis tools for RNA-seq read counts. *Genome Biol* **15**, R29-2014-15-2-r29.

Ritchie, M.E., Phipson, B., Wu, D., Hu, Y., Law, C.W., Shi, W. & Smyth, G.K. (2015) limma powers differential expression analyses for RNA-sequencing and microarray studies. *Nucleic Acids Res* **43**, e47.

Smemo, S., Tena, J.J., Kim, K.-H., Gamazon, E.R., Sakabe, N.J., Gómez-Marín, C., Aneas, I., Credidio, F.L., Sobreira, D.R., Wasserman, N.F., Lee, J.H., Puviindran, V., Tam, D., Shen, M., Son, J.E., Vakili, N.A., Sung, H.-K., Naranjo, S., Acemel, R.D., Manzanares, M., Nagy, A., Cox, N.J., Hui, C.-C., Gomez-Skarmeta, J.L. & Nóbrega, M.A. (2014) Obesity-associated variants within FTO form long-range functional connections with IRX3. *Nature* **507**, 371–375.

Sutton, L.P., Ostrovskaya, O., Dao, M., Xie, K., Orlandi, C., Smith, R., Wee, S. & Martemyanov, K.A. (2016) Regulator of G-Protein Signaling 7 Regulates Reward Behavior by Controlling Opioid Signaling in the Striatum. *BiolPsychiatry* **80**, 235–245.

Trapnell, C., Roberts, A., Goff, L., Pertea, G., Kim, D., Kelley, D.R., Pimentel, H., Salzberg, S.L., Rinn, J.L. & Pachter, L. (2012) Differential gene and transcript expression analysis of RNA-seq experiments with TopHat and Cufflinks. *NatProtoc* **7**, 562–578.

Yalcin, B., Wong, K., Agam, A., Goodson, M., Keane, T.M., Gan, X., Nellaker, C., Goodstadt, L., Nicod, J., Bhomra, A., Hernandez-Pliego, P., Whitley, H., Cleak, J., Dutton, R., Janowitz, D., Mott, R., Adams, D.J. & Flint, J. (2011) Sequence-based characterization of structural variation in the mouse genome. *Nature* **477**, 326–329.

Yazdani, N., Parker, C.C., Shen, Y., Reed, E.R., Guido, M.A., Kole, L.A., Kirkpatrick, S.L., Lim, J.E., Sokoloff, G., Cheng, R., Johnson, W.E., Palmer, A.A. & Bryant, C.D. (2015) Hnrnph1 Is A Quantitative Trait Gene for Methamphetamine Sensitivity. *PLoS Genet* **11**, e1005713.
